## Supplementary Information for "Fast-exchanging spirocyclic rhodamine probes for aptamer-based super-resolution RNA imaging"

#### Supplementary figures

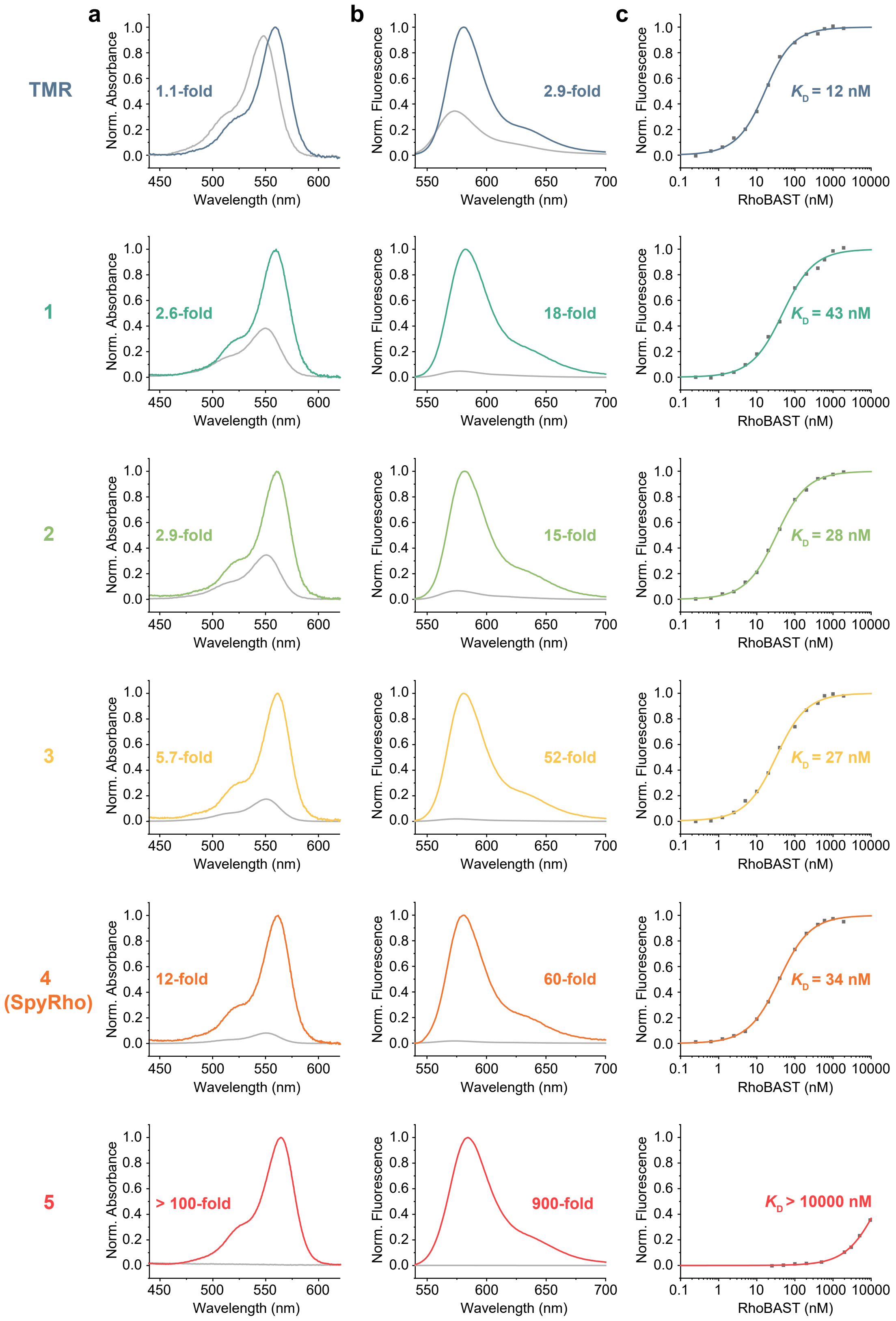


**Supplementary Figure 1. Properties of TMR and amide-substituted TMR derivatives. a)** Normalized absorption spectra of rhodamines in the presence (colored lines) and absence (grey lines) of RhoBAST. Given absorption increase corresponds to the ratio of the excitation maxima of the bound and unbound probe. **b)** Normalized fluorescence emission spectra of rhodamines in the presence (colored lines) and absence (grey lines) of RhoBAST. Absorbance and fluorescence measurements were performed in ASB at 25 °C using 1 µM dye and 5 µM RhoBAST. **c)** Isotherms of rhodamines binding to RhoBAST. Measurements were performed in ASB supplemented with 0.05% Tween20 at 25 °C using 10 nM probe (50 nM for **5**).

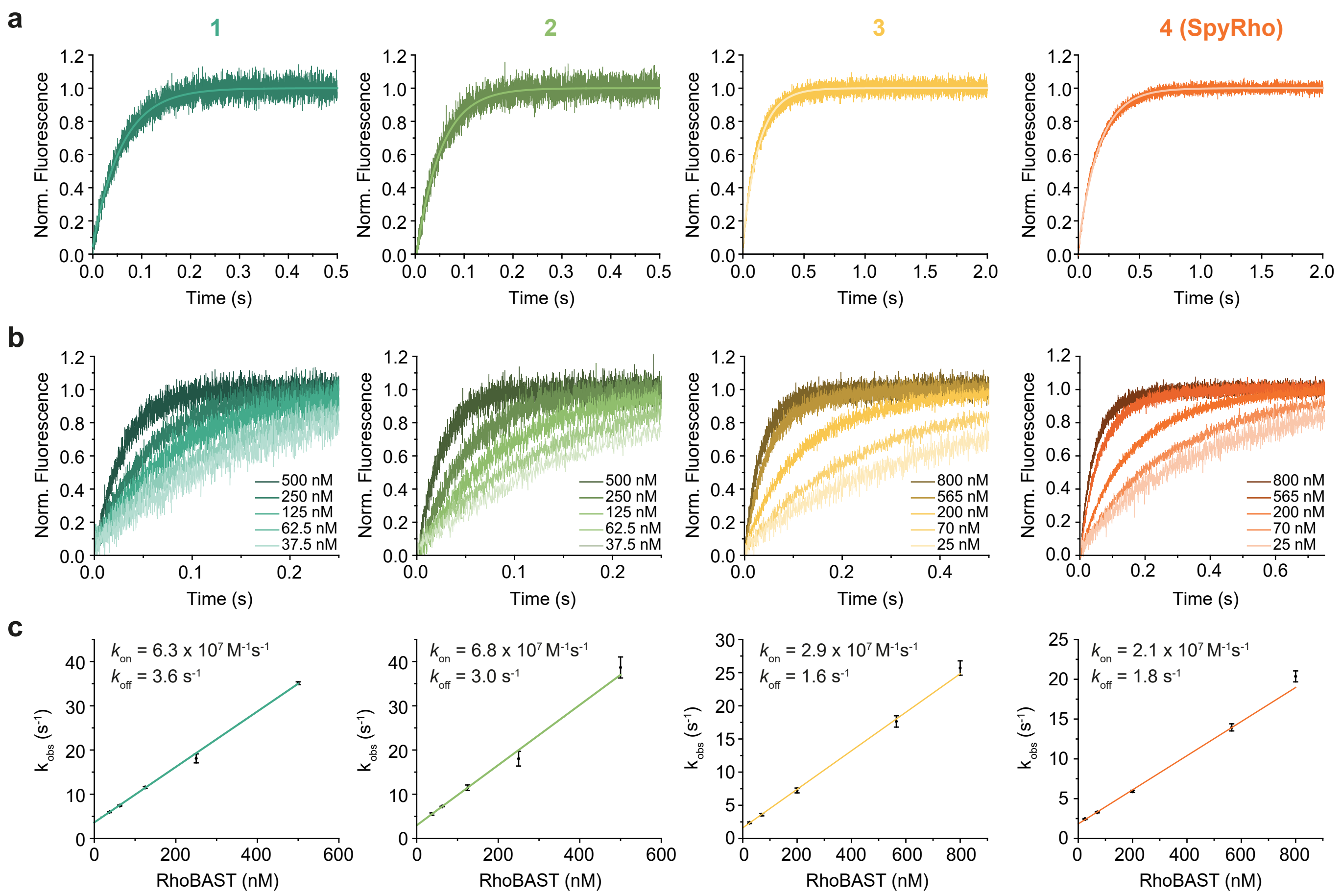


**Supplementary Figure 2: Stopped-flow binding kinetics of RhoBAST:dye complexes. a)** Normalized fluorescence increase over time upon mixing of dye (5 nM) and RhoBAST (for **1** or **2**: 250 nM RhoBAST; for **3** or **4 (SpyRho)** 200 nM RhoBAST) and single exponential fitting. **b)** Normalized fluorescence increase due to complex formation over time upon mixing of 5 nM dye with different RhoBAST concentrations at 25 °C. **c)** Observed kinetic rates ($k_{obs}$) were obtained by single-exponential fitting of the data shown in b). Linear fitting of the observed rates versus the RhoBAST concentration yields the dissociation ($k_{off}$) and association ($k_{on}$) rate coefficient. Data points represent mean ± s.d of three independent measurements.

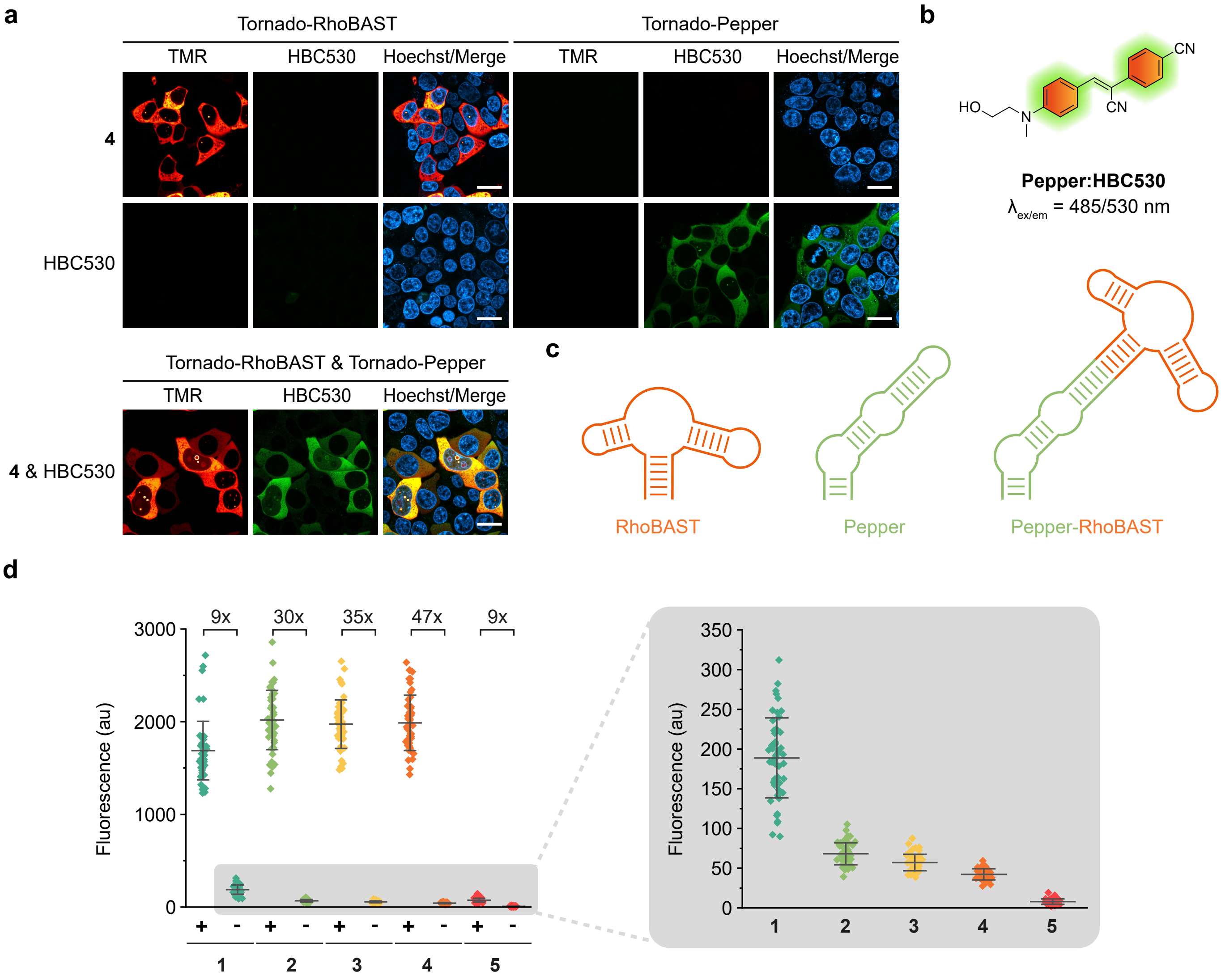


**Supplementary Figure 3. Comparison of amide-substituted TMR derivatives in live cells. a)** Orthogonality of RhoBAST and Pepper FLAP system. Confocal imaging of live HEK293T cells expressing circular RhoBAST and/or circular Pepper aptamer incubated with **4** (100 nM) and/or HBC530 (1 µM) for 30 min. **b)** Chemical structure of HBC530. **c)** Schematic drawing of RhoBAST, Pepper and the designed Pepper-RhoBAST tandem aptamer construct. **d)** Left: Figure 2e reproduced; right: background fluorescence in negative control cells (-) shown with an expanded scale. Scale bars, 20 µm.

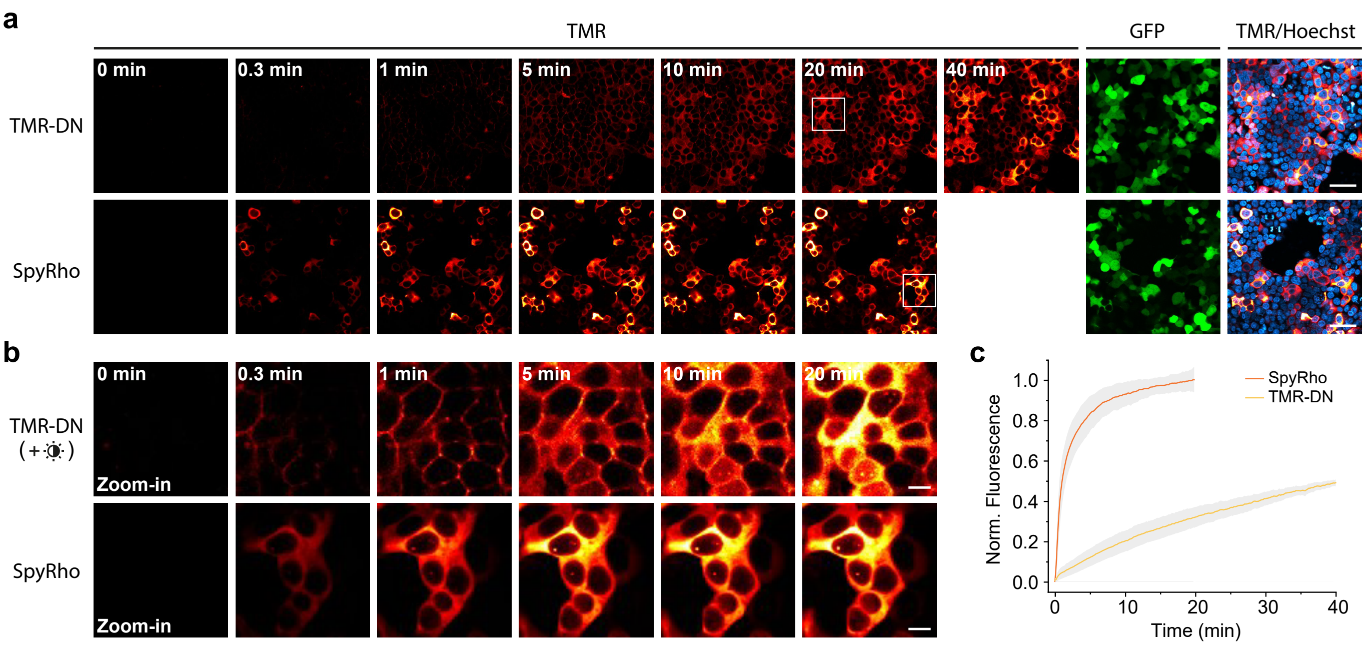


**Supplementary Figure 4. Cellular uptake of SpyRho and TMR-DN. a)** Confocal imaging of live HEK293T cells expressing circular RhoBAST and mEGFP (as transfection control) at different time points after addition of the dyes (100 nM). Scale bars, 50 µm. **b)** Zoom-ins of the highlighted regions of a). Scale bars, 10 µm. **c)** Quantification of the fluorescence increase over time (mean ± s.d.) in the cytosol of individual transfected cells (*N* ≥ 100) shown in a) The fluorescence was normalized to the average fluorescence intensity of SpyRho at *t* = 20 min.

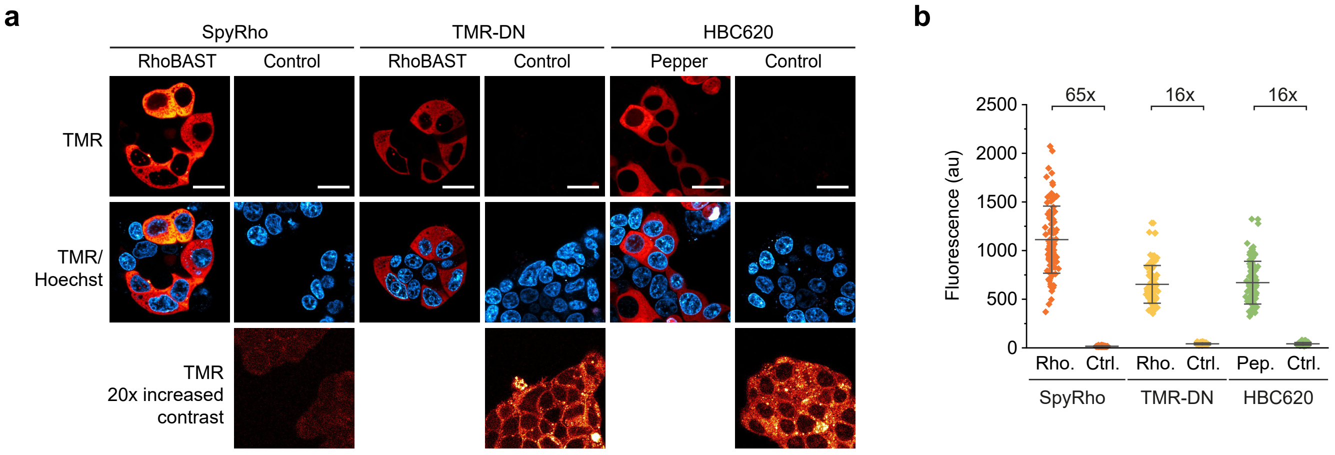


**Supplementary Figure 5.** Confocal imaging of live HEK293T cells expressing circular RhoBAST or circular Pepper aptamers, incubated with the corresponding fluorogenic probes (100 nM for SpyRho and TMR-DN, 1 µM HBC620) for 1 h. As a control, cells expressing the orthogonal aptamer (Pepper for SpyRho and TMR-DN, RhoBAST for HB620) were used. **b)** Quantification of normalized TMR fluorescence (mean ± s.d.) in the cytosol of individual transfected cells (*N* = 100) of images in a). Scale bars, 20 µm.

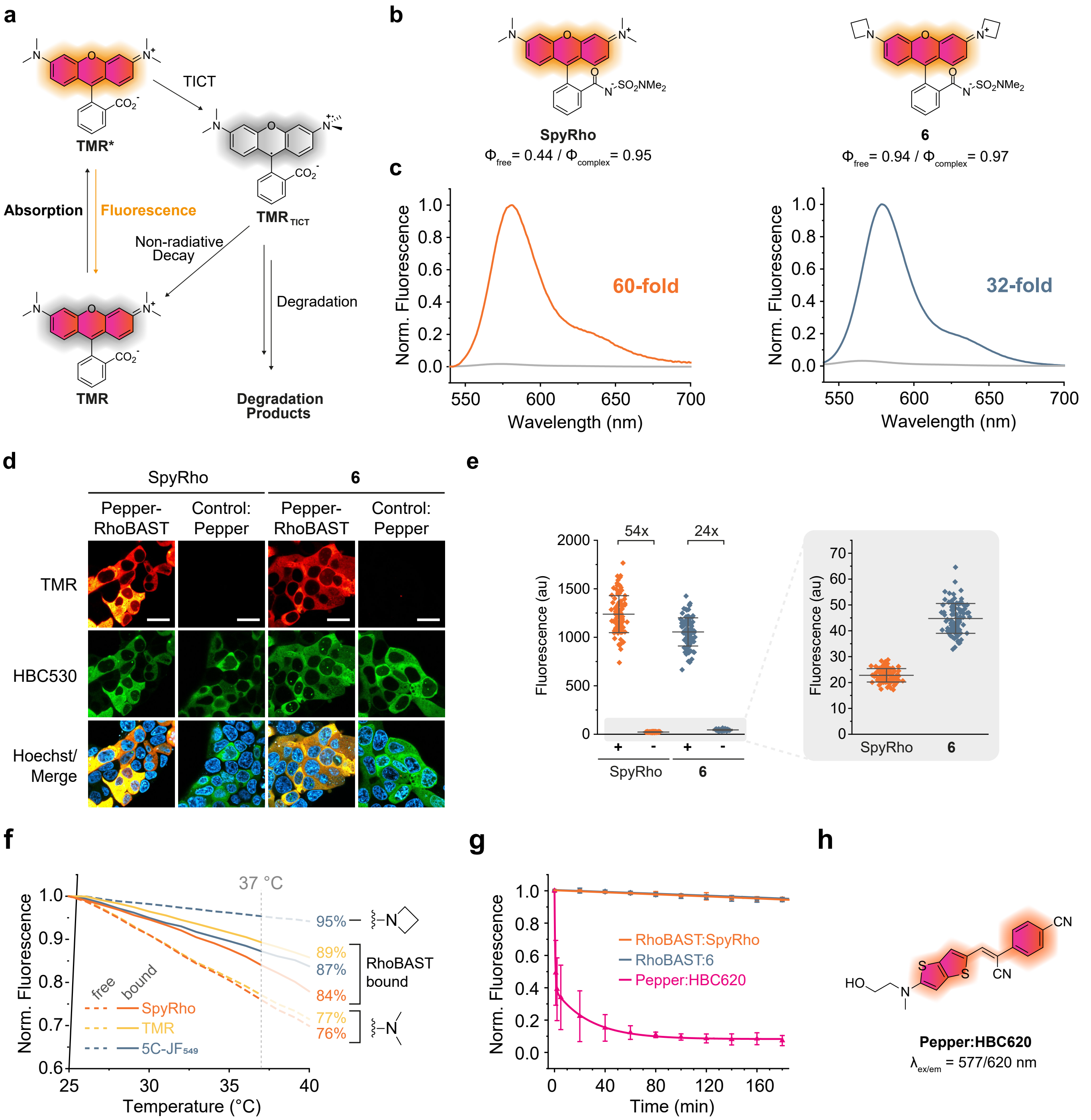


**Supplementary Figure 6. Suppression of the twisted intramolecular charge transfer (TICT) mechanism. a)** Schematic illustrating the TICT process. **b)** Chemical structure of SpyRho and the corresponding azetidinyl rhodamine **6**. **c)** Normalized emission spectra of SpyRho and **6** in the presence (colored line) and absence (grey line) of RhoBAST. Measurements were performed in ASB at 25 °C using 1 µM dye and 5 µM RhoBAST. **d)** Confocal imaging of *live* HEK293T cells expressing circular Pepper-RhoBAST or circular Pepper aptamer incubated with SpyRho or **6** (100 nM) and HBC530 (1 µM) for 1 h. **e)** Left: Quantification of normalized TMR fluorescence (mean ± s.d.) in the cytosol of individual transfected (Pepper-RhoBAST (+), Pepper (-)) cells ($N$ = 50) from images as those shown in d); right: background fluorescence in negative control cells (-), shown with an expanded scale. **f)** Temperature dependence of the fluorescence of tetramethyl and azetidinyl rhodamines (500 nM) in the absence and presence of RhoBAST (2 µM) in ASB supplemented with 0.05% Tween 20. **g)** Fluorescence decrease of aptamer:dye complexes (20 nM dye, 500 nM aptamer) under constant irradiation with an LED ( $_{max}$ = 567 nm, 680 µW mm$^{-2}$) in ASB supplemented with 0.05% Tween 20 at 25 °C. **h)** Chemical structure of HBC620. Scale bars, 20 µm.

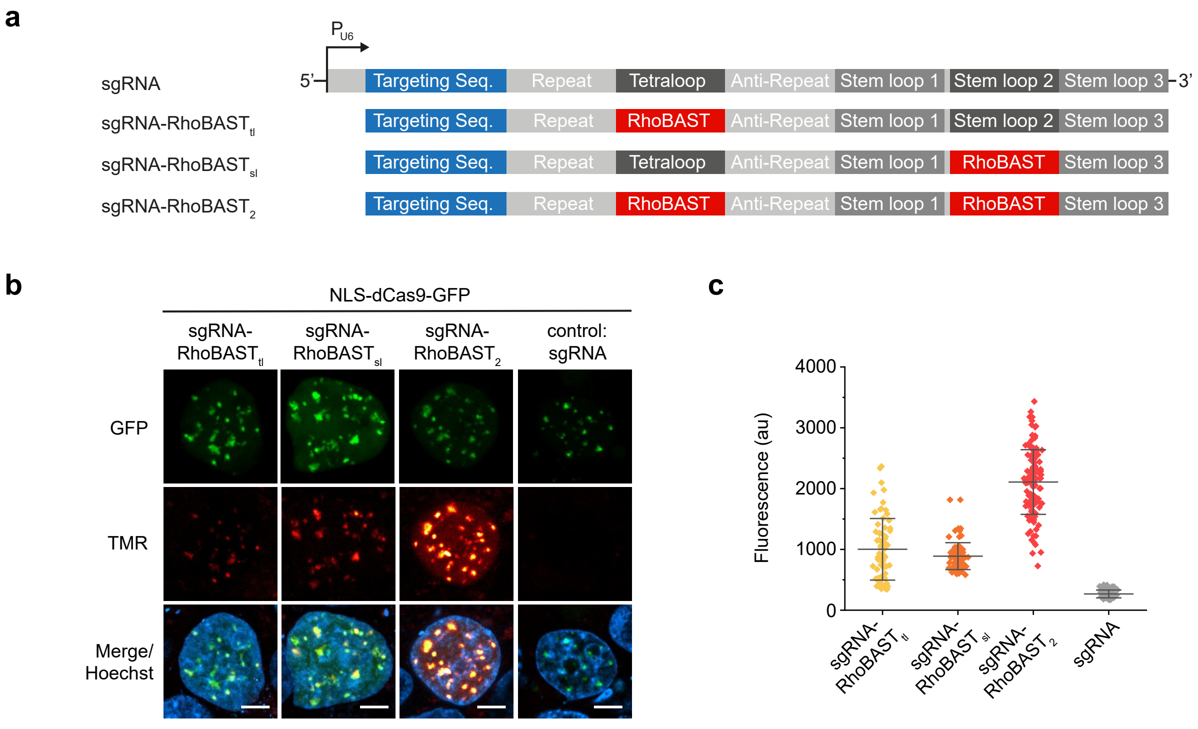


**Supplementary Figure 7. Visualization of genomic loci using RhoBAST-modified sgRNAs. a)** Schematic illustration of gene constructs for expression of unmodified and RhoBAST-modified sgRNA. **b)** Confocal imaging of HEK293T cells co-expressing NLS-dCas9-GFP and different sgRNAs constructs (plasmid ratio 1:10) targeting centromeres (targeting sequence: GAATCTGCAAGTGGATATT). Cells were incubated with SpyRho (100 nM) for 1 h prior to imaging. Maximum intensity projections of acquired z-stacks (500 nm step-size, 5 μm total) are shown for the GFP and TMR channels. **c)** Quantification of TMR fluorescence (mean ± s.d.) of nuclear foci (*N* ≥ 70) in individual transfected cells (*N* ≥ 10) of images in (c). Scale bars, 5 μm.

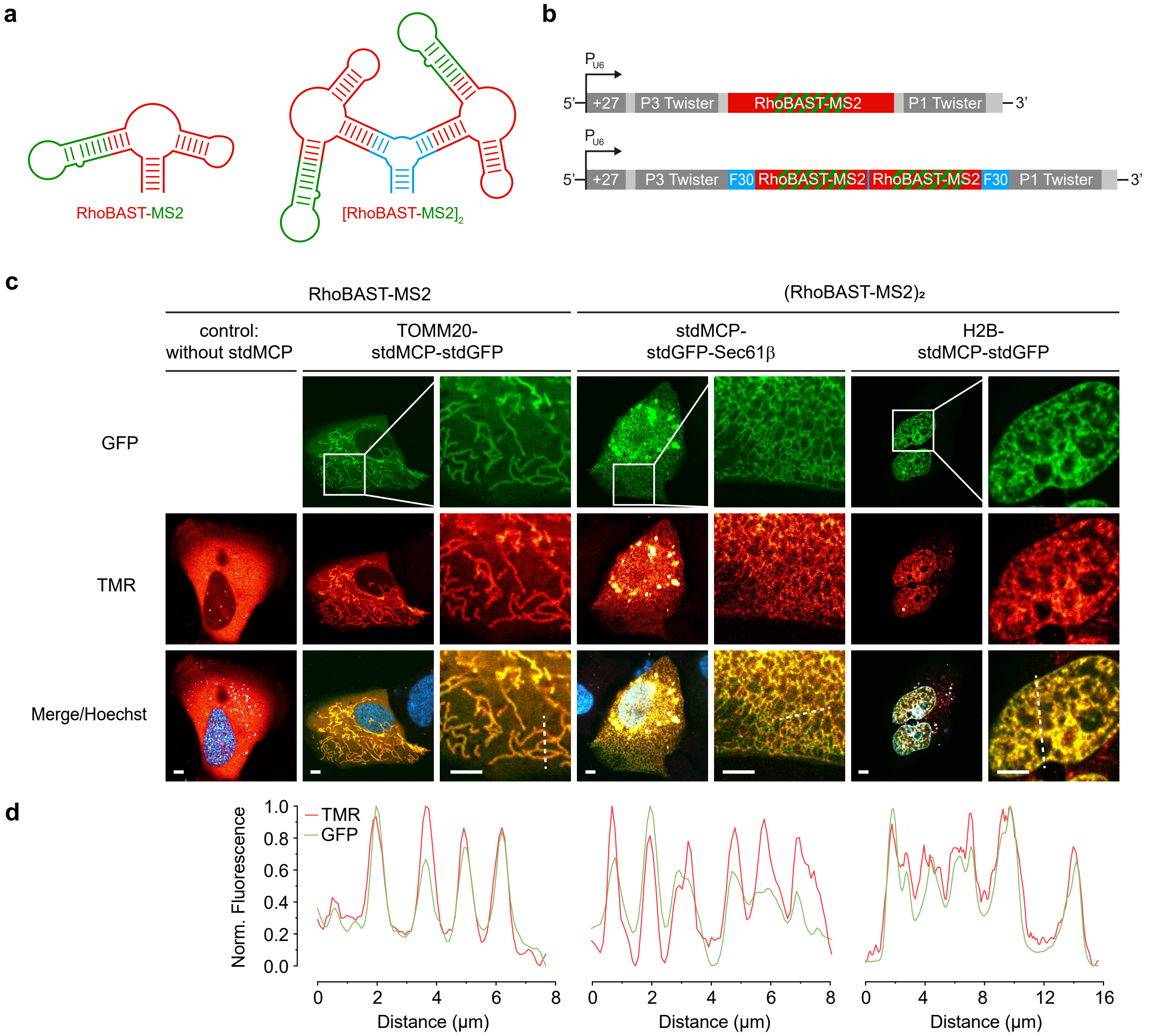


**Supplementary Figure 8. Visualization of proteins using RhoBAST-MS2 tandem constructs. a)** Schematic drawing of RhoBAST-MS2 and [RhoBAST-MS2]$_2$. **b)** Schematics of gene constructs for expressing circular *RhoBAST-MS2* and *[RhoBAST-MS2]$_2$* imager RNA. **c)** Confocal imaging of live Cos7 cells co-expressing circular *RhoBAST-MS2* or *[RhoBAST-MS2]$_2$* and different stdMCP fusion proteins (plasmid ratio 10:1). Cells were incubated with SpyRho (100 nM) for 1 h prior to imaging. For clarity, the merged zoom-ins are shown without Hoechst. **d)** Normalized fluorescence profiles of the indicated dashed lines in c). Scale bars, 5 μm.

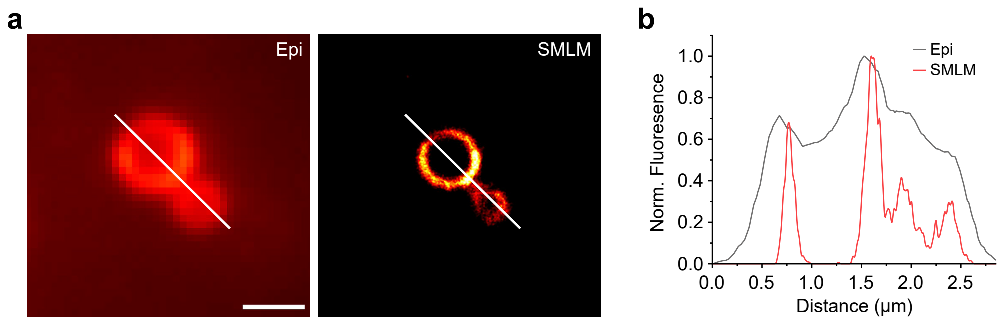


**Supplementary Figure 9. Single-molecule localization microscopy (SMLM) of circular RhoBAST. a)** Epifluorescence (Epi) and SMLM image of the nucleus of a fixed Cos7 cell expressing circular RhoBAST incubated with SpyRho (50 nM) for 30 min. The SMLM image was reconstructed using 30,000 frames (30 ms exposure time). **b)** Normalized intensity profile along the white line in a). Scale bar, 1 µm.

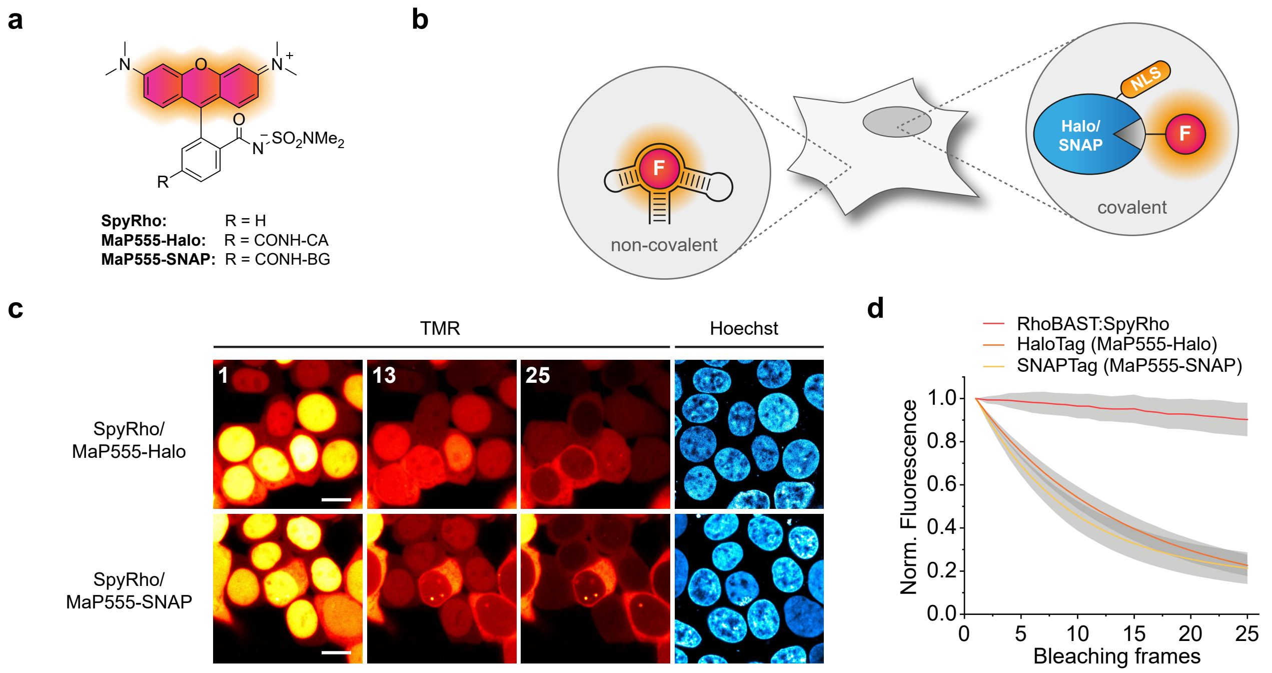


**Supplementary Figure 10. Comparison of photostability of RhoBAST:SpyRho with MaP dyes. a)** Chemical structure of SpyRho and employed MaP dyes (CA = chloroalkane; BG = benzyl-guanine). **b)** Illustration of localization of fluorescence arising from circular RhoBAST (cytosol) and HaloTag-SNAP-Tag-NLS fusion protein (nucleus) after labeling with the corresponding dyes (SpyRho, MaP555-Halo or MaP555-SNAP). **c)** Confocal imaging of HEK293T cells expressing circular RhoBAST and HaloTag-SNAP-Tag-NLS fusion protein (plasmid ratio 10:1). Cells were first incubated with the corresponding fluorogenic MaP555 derivative (MaP555-Halo: 50 nM, MaP555-SNAP: 500 nM) for 30 min;excess dye was removed by washing. Next, cells were incubated with SpyRho (100 nM) for 15 min and cells were imaged for 4 min (0.1 frames/s, 2× line accumulation, 530 µW). Numbers in the images correspond to the position in the sequence of acquired frames. **d)** Quantification of TMR fluorescence normalized to the initial value (mean ± s.d.) in the cytosol and nucleus of individual transfected cells ($N$ = 30) of images as those shown in c). Scale bars, 10 µm.

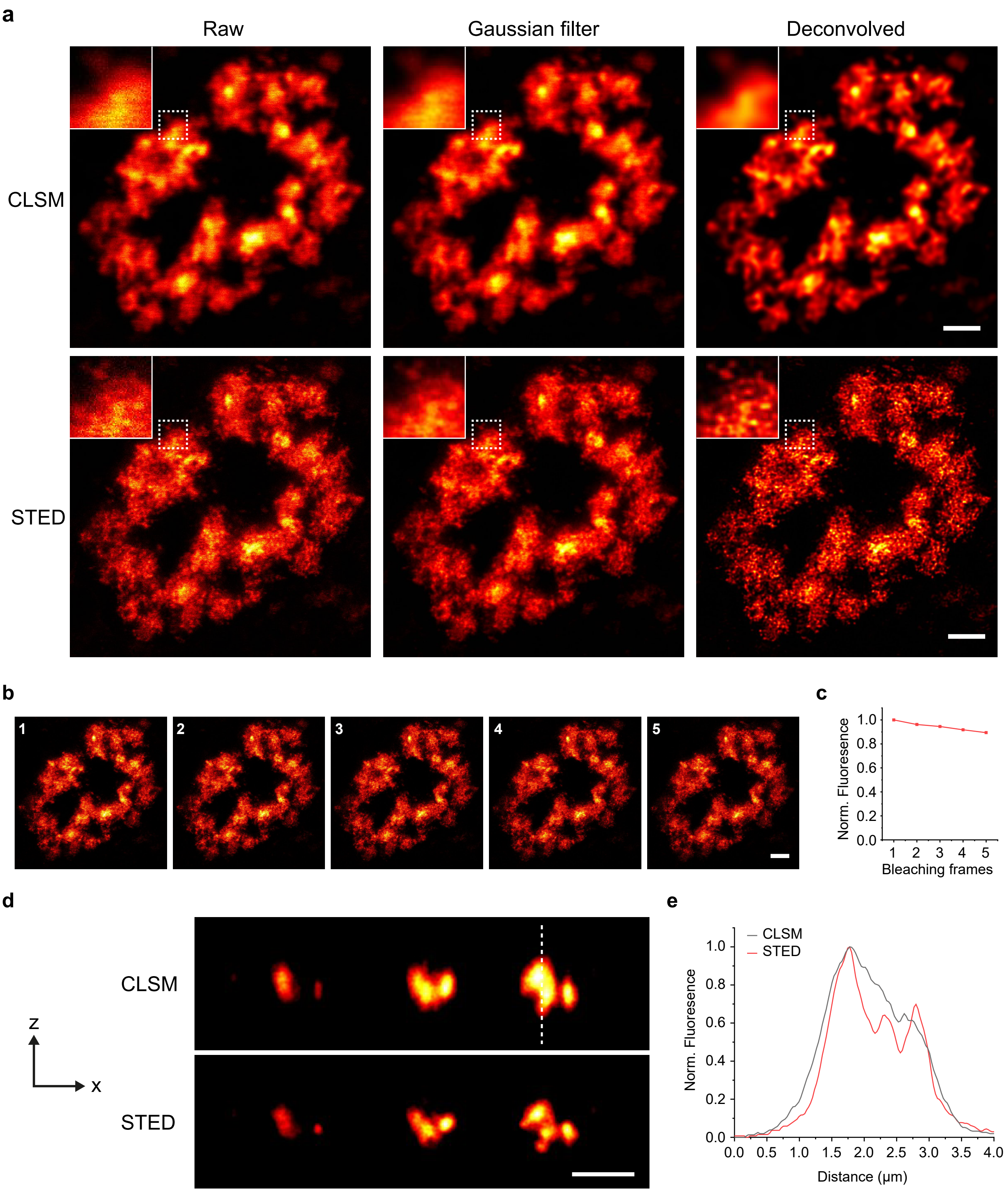


**Supplementary Figure 11. Stimulated emission depletion (STED) RNA imaging. a)** Raw, Gaussian-filtered and deconvolved confocal laser scanning microscopy (CLSM) and 2D-STED images of a nucleus of a live Cos7 cells expressing *CGG₉₉-FMR1-GFP-RhoBAST₁₆* mRNA incubated with SpyRho (100 nM) for 1 h. Magnification of indicated region (dashed box) is shown as inset. **b)** Time-lapse STED imaging of live Cos7 cells expressing *CGG₉₉-FMR1-GFP-RhoBAST₁₆* mRNA incubated with SpyRho (100 nM) for 1 h. Each frame was acquired using 16× line accumulation. Frames were acquired continuously. **c)** Quantification of the normalized TMR fluorescence of the images shown in a). **d)** XZ scans using 3D-STED mode of the nucleus of a live Cos7 cells expressing *CGG₉₉-FMR1-GFP-RhoBAST₁₆* mRNA incubated with SpyRho (100 nM) for 1 h. **e)** Normalized fluorescence profile along the dashed line shown in d). Scale bars, 2 µm.

### Supplementary Tables

**Supplementary Table 1. Properties of fluorogenic rhodamines and the corresponding RhoBAST:dye complexes.** The excitation wavelength ($\lambda_{ex}$), emission wavelength ($\lambda_{em}$), peak extinction coefficient ($\varepsilon$), fluorescence quantum yield ($\Phi_F$), and fluorescence turn-on (1 µM probe, 5 µM RhoBAST) were measured in ASB at 25 °C. Absolute quantum yields were measured for rhodamines; values for RhoBAST:dye complexes were determined relative to sulforhodamine 101. The brightness was calculated as the product of $\varepsilon$ and $\Phi_F$. Equilibrium dissociation constants ($K_D$) were measured in ASB supplemented with 0.05% Tween 20 at 25°C using 10 nM probe.

| | Free dye | | | | | | RhoBAST:dye complex | | | | | | | |
|---|---|---|---|---|---|---|---|---|---|---|---|---|---|---|
| | $\lambda_{ex}$ [nm] | $\lambda_{em}$ [nm] | $\varepsilon$ [$M^{-1}cm^{-1}$] | $\Phi_F$ | Brightness [$M^{-1}cm^{-1}$] | $D_{50}$ | $\lambda_{ex}$ [nm] | $\lambda_{em}$ [nm] | $\varepsilon$ [$M^{-1}cm^{-1}$] | $\Phi_F$ | Brightness [$M^{-1}cm^{-1}$] | Abs. increase [n-fold] | Turn-on [n-fold] | $K_D$ [nM] |
| **TMR** | 549 | 573 | 75000 ± 3000 | 0.48 | 36000 | 12 | 560 | 581 | 80000 ± 3000 | 0.92 ± 0.02 | 74000 | 1.1 ± 0.1 | 2.9 ± 0.2 | 12 ± 1 |
| **1** | 551 | 578 | 23000 ± 2000 | 0.46 | 11000 | 43 | 560 | 581 | 60000 ± 1000 | 0.98 ± 0.03 | 59000 | 2.6 ± 0.3 | 18 ± 4 | 43 ± 4 |
| **2** | 552 | 575 | 24000 ± 1000 | 0.47 | 11000 | 55 | 561 | 582 | 69000 ± 1000 | 0.95 ± 0.02 | 66000 | 2.9 ± 0.2 | 15 ± 1 | 28 ± 1 |
| **3** | 551 | 575 | 12000 ± 2000 | 0.45 | 5400 | 70 | 561 | 581 | 69000 ± 1000 | 0.97 ± 0.03 | 67000 | 5.7 ± 0.9 | 52 ± 19 | 27 ± 2 |
| **4 (SpyRho)** | 551 | 573 | 5300 ± 600 | 0.44 | 2300 | 70 | 562 | 581 | 65000 ± 1000 | 0.95 ± 0.03 | 62000 | 12 ± 2 | 60 ± 18 | 34 ± 2 |
| **5** | 549 | 571 | <200 | - | - | >70 | 565 | 584 | ND[a] | 0.98 ± 0.02 | ND[a] | >100 | >900 | >10000[b] |
| **6** | 552 | 573 | 6700 ± 100 | 0.94 | 6300 | >70 | 562 | 580 | 74000 ± 3000 | 0.97 ± 0.03 | 72000 | 11 ± 1 | 32 ± 7 | 100 ± 8 |

[a] Not determined (ND) due to incomplete complexation.
[b] Dissociation constant was determined using 50 nM probe.

**Supplementary Table 2. Kinetic parameters.** Association ($k_{on}$) and dissociation ($k_{off}$) rate coefficients of the RhoBAST:dye complexes were obtained from stopped-flow measurements in ASB supplemented with 0.05% Tween20 at 25 °C.

| | $k_{on}$ [$M^{-1}s^{-1}$] | $k_{off}$ [$s^{-1}$] |
|---|---|---|
| RhoBAST:**1** | $6.3 \pm 0.1 \times 10^7$ | $3.6 \pm 0.1$ |
| RhoBAST:**2** | $6.8 \pm 0.3 \times 10^7$ | $3.0 \pm 0.2$ |
| RhoBAST:**3** | $2.9 \pm 0.1 \times 10^7$ | $1.6 \pm 0.1$ |
| RhoBAST:SpyRho (**4**) | $2.1 \pm 0.1 \times 10^7$ | $1.8 \pm 0.1$ |
| RhoBAST:**6** | $1.2 \pm 0.1 \times 10^7$ | $2.6 \pm 0.1$ |

**Supplementary Table 3. DNA Sequences.**

<table>
<tr><td rowspan="2">pAV-U6+27-tornado-RhoBAST</td><td>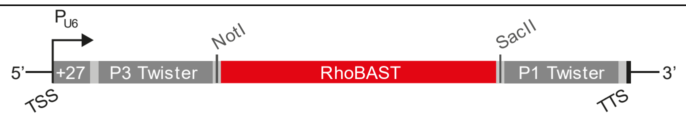
</td></tr>
<tr><td>GTCGACGGGCCGCACTCGCCGGTCCCAAGCCCGGATAAAATGGGAGG<br>GGGCGGGAAACCGCCTAACCATGCCGAGTGCGGCCGCACCTCCGCGA<br>AAGCGGTGAAGGAGAGGCGCAAGGTTAACCGCCTCAGGTGTGGCCGC<br>GGTCGGCGTGGACTGTAGAACACTGCCAATGCCGGTCCCAAGCCCGG<br>ATAAAAGTGGAGGGTACAGTCCACGCTCTAGA</td></tr>
<tr><td rowspan="2">pAV-U6+27-tornado-Pepper</td><td>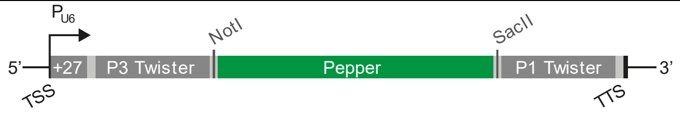
</td></tr>
<tr><td>GTCGACGGGCCGCACTCGCCGGTCCCAAGCCCGGATAAAATGGGAGG<br>GGGCGGGAAACCGCCTAACCATGCCGAGTGCGGCCGCCCCAATCGTG<br>GCGTGTCGGCCTGCTTCGGCAGGCACTGGCGCCGGGTGGCCGCGGTC<br>GGCGTGGACTGTAGAACACTGCCAATGCCGGTCCCAAGCCCGGATAA<br>AAGTGGAGGGTACAGTCCACGCTCTAGA</td></tr>
<tr><td rowspan="2">pAV-U6+27-tornado-Pepper-RhoBAST</td><td>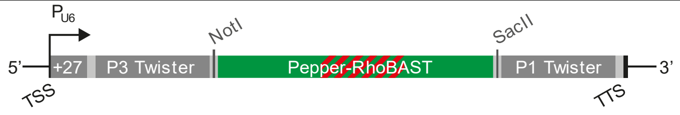
</td></tr>
<tr><td>GTCGACGGGCCGCACTCGCCGGTCCCAAGCCCGGATAAAATGGGAGG<br>GGGCGGGAAACCGCCTAACCATGCCGAGTGCGGCCGCCCCAATCGTG<br>GCGTGTCGGCCTGCAACCTCCGCGAAAGCGGTGAAGGAGAGGCGCAA<br>GGTTAACCGCCTCAGGTTGCAGGCACTGGCGCCGGGTGGCCGCGGTC<br>GGCGTGGACTGTAGAACACTGCCAATGCCGGTCCCAAGCCCGGATAA<br>AAGTGGAGGGTACAGTCCACGCTCTAGA</td></tr>
<tr><td rowspan="2">pAV-U6+27-tornado-RhoBAST-MS2</td><td>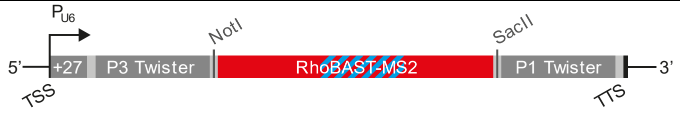
</td></tr>
<tr><td>GTCGACGGGCCGCACTCGCCGGTCCCAAGCCCGGATAAAATGGGAGG<br>GGGCGGGAAACCGCCTAACCATGCCGAGTGCGGCCGCACCTCGGCGG<br>CCAACATGAGGATCACCCATGTCTGCAGGGCCGCCGTGAAGGAGAGG<br>CGCAAGGTTAACCGCCTCAGGTGTGGCCGCGGTCGGCGTGGACTGTA<br>GAACACTGCCAATGCCGGTCCCAAGCCCGGATAAAAGTGGAGGGTAC<br>AGTCCACGCTCTAGA</td></tr>
<tr><td rowspan="2">pAV-U6+27-tornado-[RhoBAST-MS2]$_2$</td><td>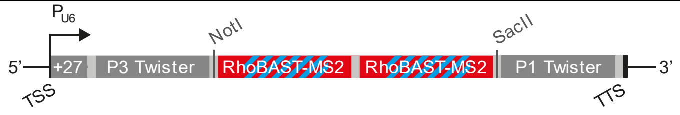
</td></tr>
<tr><td>GTCGACGGGCCGCACTCGCCGGTCCCAAGCCCGGATAAAATGGGAGG<br>GGGCGGGAAACCGCCTAACCATGCCGAGTGCGGCCGCTTGCCATGTG</td></tr>
</table>

<table>
<tr><td></td><td>TATCGGAAGGCCTCCGCGCCCAGGTGGAGGATCACCCCACCCTGCAG<br>GGGCGCGGTGAAGGAGCGGCACAAGGTTAACTGCCGCAGGCCTTCCG<br>ATACTCTGATGATCCGGAACCTCCGGGGCCAACATGAGGATCACCCA<br>TGTCTGCAGGGCCCCGGTGAAGGAGAGGCGCAAGGTTAACCGCCTCA<br>GGTTCCGGATCATTCATGGCAAGTGGCCGCGGTCGGCGTGGACTGTA<br>GAACACTGCCAATGCCGGTCCCAAGCCCGGATAAAAGTGGAGGGTAC<br>AGTCCACGCTCTAGA</td></tr>
<tr><td rowspan="2">pcDNA3-HaloTag7-Sam68</td><td>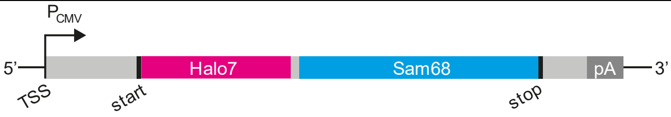
</td></tr>
<tr><td>AAGCTTCGCCGCCACCATGGATCCGAAATCGGTACTGGCTTTCCAT<br>TCGACCCCCATTATGTGGAAGTCCTGGGCGAGCGCATGCACTACGTC<br>GATGTTGGTCCGCGCGATGGCACCCCTGTGCTGTTCCTGCACGGTAA<br>CCCGACCTCCTCCTACGTGTGGCGCAACATCATCCCGCATGTTGCAC<br>CGACCCATCGCTGCATTGCTCCAGACCTGATCGGTATGGGCAAATCC<br>GACAAACCAGACCTGGGTTATTTCTTCGACGACCACGTCCGCTTCAT<br>GGATGCCTTCATCGAAGCCCTGGGTCTGGAAGAGGTCGTCCTGGTCA<br>TTCACGACTGGGGCTCCGCTCTGGGTTTCCACTGGGCCAAGCGCAAT<br>CCAGAGCGCGTCAAAGGTATTGCATTTATGGAGTTCATCCGCCCTAT<br>CCCGACCTGGGACGAATGGCCAGAATTTGCCCGCGAGACCTTCCAGG<br>CCTTCCGCACCACCGACGTCGGCCGCAAGCTGATCATCGATCAGAAC<br>GTTTTTATCGAGGGTACGCTGCCGATGGGTGTCGTCCGCCCGCTGAC<br>TGAAGTCGAGATGGACCATTACCGCGAGCCGTTCCTGAATCCTGTTG<br>ACCGCGAGCCACTGTGGCGCTTCCCAAACGAGCTGCCAATCGCCGGT<br>GAGCCAGCGAACATCGTCGCGCTGGTCGAAGAATACATGGACTGGCT<br>GCACCAGTCCCCTGTCCCGAAGCTGCTGTTCTGGGGCACCCCAGGCG<br>TTCTGATCCCACCGGCCGAAGCCGCTCGCCTGGCCAAAAGCCTGCCT<br>AACTGCAAGGCTGTGGACATCGGCCCGGGTCTGAATCTGCTGCAAGA<br>AGACAACCCGGACCTGATCGGCAGCGAGATCGCGCGCTGGCTGTCTA<br>CTCTGGAGATTTCCGGTGGTTCCGGCTCAGGATCTAGAGGATCC [...]<br>ATCCATATGGACGTTATTAAAAACAAACAGGAGGGA</td></tr>
<tr><td rowspan="2">pcDNA5-FRT/TO-NLS-dCas9-NLS-GFP</td><td>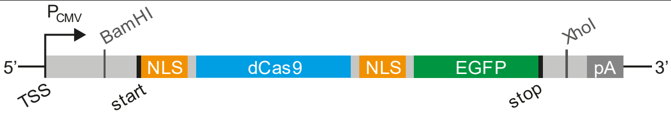
</td></tr>
<tr><td>CCGAGCTCGGATCCTCGCCACCATGGCTAGCCCCAAAAAGAAGAGGA<br>AAGTGGACAAGAAGTATTCT [...] GCTCGGAGGGGATGAGGGAGCTCC<br>CAAGAAAAAGCGCAAGGTAGGTAGTTCCGTGAGCAAGGGCGAGGAGC<br>TGTTCACCGGGGTGGTGCCCATCCTGGTCGAGCTGGACGGCGACGTA<br>AACGGCCACAAGTTCAGCGTGTCCGGCGAGGGCGAGGGCGATGCCAC<br>CTACGGCAAGCTGACCCTGAAGTTCATCTGCACCACCGGCAAGCTGC<br>CCGTGCCCTGGCCCACCCTCGTGACCACCCTGACCTACGGCGTGCAG<br>TGCTTCAGCCGCTACCCCGACCACATGAAGCAGCACGACTTCTTCAA<br>GTCCGCCATGCCCGAAGGCTACGTCCAGGAGCGCACCATCTTCTTCA<br>AGGACGACGGCAACTACAAGACCCGCGCCGAGGTGAAGTTCGAGGGC<br>GACACCCTGGTGAACCGCATCGAGCTGAAGGGCATCGACTTCAAGGA<br>GGACGGCAACATCCTGGGGCACAAGCTGGAGTACAACTACAACAGCC<br>ACAACGTCTATATCATGGCCGACAAGCAGAAGAACGGCATCAAGGTG<br>AACTTCAAGATCCGCCACAACATCGAGGACGGCAGCGTGCAGCTCGC<br>CGACCACTACCAGCAGAACACCCCCATCGGCGACGGCCCCGTGCTGC<br>TGCCCGACAACCACTACCTGAGCACCCAGTCCGCCCTGAGCAAAGAC<br>CCCAACGAGAAGCGCGATCACATGGTCCTGCTGGAGTTCGTGACCGC<br>CGCCGGGATCACTCTCGGCATGGACGAGCTGTACAAGTAAGCTCGAG<br>TCTAG</td></tr>
<tr><td>pAV-U6-centromer-sgRNA</td><td>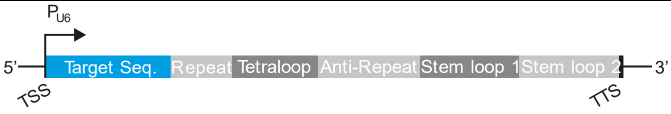
</td></tr>
</table>

| | |
|---|---|
| | GCC***GGATCC***AAGGTCGGGCAGGAAGAGGGCCTATTTCCCATGATTCC<br>TTCATATTTGCATATACGATACAAGGCTGTTAGAGAGATAATTGGAA<br>TTAATTTGACTGTAAACACAAAGATATTAGTACAAAATACGTGACGT<br>AGAAAGTAATAATTTCTTGGGTAGTTTGCAGTTTTAAAATTATGTTT<br>TAAAATGGACTATCATATGCTTACCGTAACTTGAAAGTATTTCGATT<br>TCTTGGCTTTATATATCTTGTGGAAAGGACGAAACACC**GAATCTGCA**<br>**AGTGGATATTGTTTGAGAGCTAGAAATAGCAAGTTCAAATAAGGCTA**<br>**GTCCGTTATCAACTTGAAAAAGTGGCACCGAGTCGGTGC**TTTTTTGT<br>TTTA***CTCGAG***CCGC |
| pAV-U6-centromer-sgRNA- RhoBAST-1 | 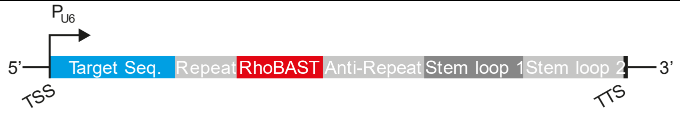 |
| | GCC***GGATCC***AAGGTCGGGCAGGAAGAGGGCCTATTTCCCATGATTCC<br>TTCATATTTGCATATACGATACAAGGCTGTTAGAGAGATAATTGGAA<br>TTAATTTGACTGTAAACACAAAGATATTAGTACAAAATACGTGACGT<br>AGAAAGTAATAATTTCTTGGGTAGTTTGCAGTTTTAAAATTATGTTT<br>TAAAATGGACTATCATATGCTTACCGTAACTTGAAAGTATTTCGATT<br>TCTTGGCTTTATATATCTTGTGGAAAGGACGAAACACC**GAATCTGCA**<br>**AGTGGATATTGTTTGAGAGCTAGGAACCTCCGCGAAAGCGGTGAAGG**<br>**AGAGGCGCAAGGTTAACCGCCTCAGGTTCCTAGCAAGTTCAAATAAG**<br>**GCTAGTCCGTTATCAACTTGAAAAAGTGGCACCGAGTCGGTGC**TTTT<br>TTGTTTTA***CTCGAG***CCGC |
| pAV-U6-centromer-sgRNA-RhoBAST-2 | 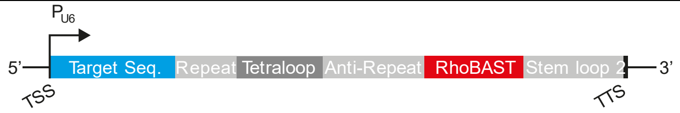 |
| | GCC***GGATCC***AAGGTCGGGCAGGAAGAGGGCCTATTTCCCATGATTCC<br>TTCATATTTGCATATACGATACAAGGCTGTTAGAGAGATAATTGGAA<br>TTAATTTGACTGTAAACACAAAGATATTAGTACAAAATACGTGACGT<br>AGAAAGTAATAATTTCTTGGGTAGTTTGCAGTTTTAAAATTATGTTT<br>TAAAATGGACTATCATATGCTTACCGTAACTTGAAAGTATTTCGATT<br>TCTTGGCTTTATATATCTTGTGGAAAGGACGAAACACC**GAATCTGCA**<br>**AGTGGATATTGTTTGAGAGCTAGAAATAGCAAGTTCAAATAAGGCTA**<br>**GTCCGTTATCAACTTGGAACCTCCGCGAAAGCGGTGAAGGAGAGGCG**<br>**CAAGGTTAACCGCCTCAGGTTCCAAGTGGCACCGAGTCGGTGC**TTTT<br>TTGTTTTA***CTCGAG***CCGC |
| pAV-U6-centromer-sgRNA-RhoBAST$_2$ | 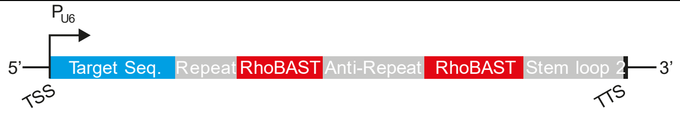 |
| | GCC***GGATCC***AAGGTCGGGCAGGAAGAGGGCCTATTTCCCATGATTCC<br>TTCATATTTGCATATACGATACAAGGCTGTTAGAGAGATAATTGGAA<br>TTAATTTGACTGTAAACACAAAGATATTAGTACAAAATACGTGACGT<br>AGAAAGTAATAATTTCTTGGGTAGTTTGCAGTTTTAAAATTATGTTT<br>TAAAATGGACTATCATATGCTTACCGTAACTTGAAAGTATTTCGATT<br>TCTTGGCTTTATATATCTTGTGGAAAGGACGAAACACC**GAATCTGCA**<br>**AGTGGATATTGTTTGAGAGCTAGGAACCTCCGCGAAAGCGGTGAAGG**<br>**AGAGGCGCAAGGTTAACCGCCTCAGGTTCCTAGCAAGTTCAAATAAG**<br>**GCTAGTCCGTTATCAACTTAAGGCCTCCGCGAAAGCGGTGAAGGAGC**<br>**GGCACAAGGTTAACTGCCGCAGGCCTTAAGTGGCACCGAGTCGGTGC**<br>TTTTTTGTTTTA***CTCGAG***CCGC |
| pcDNA5-FRT-H2B-stdMCP-stdGFP | 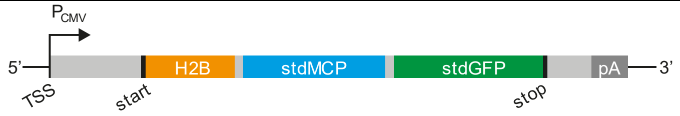 |
| | AAGCTTGGCCACC**ATGCCAGAGCCAGCGAAGTCTGCTCCCGCCCCGA**<br>**AAAAGGGCTCCAAGAAGGCGGTGACTAAGGCGCAGAAGAAAGGCGGC**<br>**AAGAAGCGCAAGCGCAGCCGCAAGGAGAGCTATTCCATCTATGTGTA**<br>**CAAGGTTCTGAAGCAGGTCCACCCTGACACCGGCATTTCGTCCAAGG** |

<table>
<tr><td></td><td>CCATGGGCATCATGAATTCGTTTGTGAACGACATTTTCGAGCGCATC<br>GCAGGTGAGGCTTCCCGCCTGGCGCATTACAACAAGCGCTCGACCAT<br>CACCTCCAGGGAGATCCAGACGGCCGTGCGCCTGCTGCTGCCTGGGG<br>AGTTGGCCAAGCACGCCGTGTCCGAGGGTACTAAGGCCATCACCAAG<br>TACACCAGCGCTAAGCTGTTATTAATTAACGCTTCTAACT [...] ATGG<br>ATGAATTGTACAAATAATCTAGAG</td></tr>
<tr><td rowspan="2">pcDNA5-FRT-TOMM20-stdMCP-stdGFP</td><td>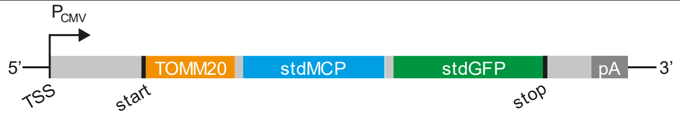
</td></tr>
<tr><td>AAGCTTGGCCACCATGGGTCGGAACAGCGCCATCGCCGCGGGCGTGT<br>GCGGTGCCCTCTTCATAGGGTACTGCATCTACTTTGACCGCAAAAGA<br>CGAAGTGACCCCAACTTCCTGTTATTAATTAACGCTTCTAACT [...] A<br>TGGATGAATTGTACAAATAATCTAGAG</td></tr>
<tr><td rowspan="2">pcDNA5-FRT-stdMCP-stdGFP- Sec61β</td><td>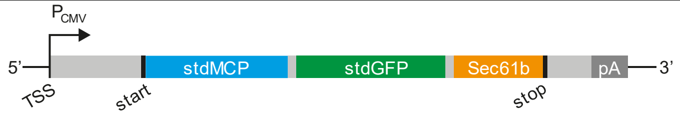
</td></tr>
<tr><td>AAGCTTGGCCACCATGGCTTCTAACT [...] ATGGATGAATTGTACAAA<br>TCCGGACTCAGATCTGGCTCCAGCGCAGGCAGCGCATCCGGCGGAAG<br>CGGAAGCCCTGGTCCGACCCCCAGTGGCACTAACGTGGGATCCTCAG<br>GGCGCTCTCCCAGCAAAGCAGTGGCCGCCCGGGCGGCGGGATCCACT<br>GTCCGGCAGAGGAAAAATGCCAGCTGTGGGACAAGGAGTGCAGGCCG<br>CACAACCTCGGCAGGCACCGGGGGATGTGGCGATTCTACACAGAAG<br>ATTCACCTGGGCTCAAAGTTGGCCCTGTTCCAGTATTGGTTATGAGT<br>CTTCTGTTCATCGCTTCTGTATTTATGTTGCACATTTGGGGCAAGTA<br>CACTCGTTCGTAGTCTAGAG</td></tr>
</table>

**Supplementary Table 4. Properties of fluorophore-quencher conjugates and the corresponding RhoBAST:dye complexes.** The excitation wavelength ($\lambda_{ex}$), emission wavelength ($\lambda_{em}$), extinction coefficient ($\varepsilon$), fluorescence quantum yield ($\Phi_F$), and fluorescence turn-on (1 µM probe, 5 µM RhoBAST) were measured in ASB at 25 °C. Absolute quantum yields were measured for rhodamines; values for RhoBAST:dye complexes were determined relative to sulforhodamine 101. The brightness was calculated as product of $\varepsilon$ and $\Phi_F$. $K_D$ values were measured in ASB supplemented with 0.05% Tween 20 at 25 °C using 10 nM probe.

| | free dye | | | | | RhoBAST:dye complex | | | | | | |
|---|---|---|---|---|---|---|---|---|---|---|---|---|
| | $\lambda_{ex}$ [nm] | $\lambda_{em}$ [nm] | $\varepsilon$ [$M^{-1}cm^{-1}$] | $\Phi_F$ | Brightness [$M^{-1}cm^{-1}$] | $\lambda_{ex}$ [nm] | $\lambda_{em}$ [nm] | $\varepsilon$ [$M^{-1}cm^{-1}$] | $\Phi_F$ | Brightness [$M^{-1}cm^{-1}$] | Turn-on [n-fold] | $K_D$ [nM] |
| **TMR** | 549 | 573 | 75000 ± 3000 | 0.48 | 36000 | 560 | 581 | 80000 ± 3000 | 0.92 ± 0.02 | 74000 | 2.9 ± 0.2 | 12 ± 1 |
| **5C-TMR** | 551 | 578 | 77000 ± 1000 | 0.43 | 33000 | 563 | 587 | - | - | - | - | 32 ± 6 |
| **TMR-DN[1]** | 553 | 582 | 64000 | 0.027 | 1700 | 564 | 590 | 96000 | 0.57 ± 0.04 | 55000 | 26 | 15 |
| **5C-JF$_{549}$** | 552 | 578 | 81000 ± 1000 | 0.92 | 75000 | 564 | 584 | - | - | - | - | 35 ± 1 |
| **JF$_{549}$-DN** | 554 | 581 | 62000 ± 1000 | 0.061 ± 0.002[a] | 3800 | 565 | 586 | 88000 ± 4000 | 0.55 ± 0.02 | 48000 | 30 ± 10 | 15 ± 1 |

[a] Quantum yield was determined relative to sulforhodamine 101.

### Supplementary Note 1:

#### Limitations of fluorophore-quencher conjugates due to residual quenching

Initially, we aimed to enhance the brightness of the established FLAP RhoBAST:TMR-DN by increasing the fluorescence quantum yield, $_{F}$, of the complex. Accordingly, we synthesized the fluorophore-quencher conjugate $JF_{549}$-DN, in which the dimethylamino groups of the TMR fluorophore were replaced by strained azetidine heterocycles (Supplementary Fig. N1a,b). This small structural modification is known to substantially increase the quantum yield of tetramethyl rhodamines from about 0.4 to 0.9 *via* suppression of the twisted intramolecular charge transfer (TICT) non-radiative decay channel.[2]

The prepared $JF_{549}$-DN probe displayed similar binding properties to RhoBAST than TMR-DN, specifically, an equilibrium dissociation coefficient $K_D$ = 15 nM and a 30-fold fluorescence turn-on (Supplementary Note Fig. 1c-e, Supplementary Table 4). Surprisingly, the RhoBAST:$JF_{549}$-DN ( $_{F}$ = 0.55) complex had a similar low quantum yield as RhoBAST:TMR-DN ( $_{F}$ = 0.57), despite TICT inhibition. Thus, we hypothesized that the fluorescence quenching by the conjugated DN is not completely abolished through the binding of RhoBAST. Indeed, the quantum yield of RhoBAST:TMR (without quencher moiety) was very high ( $_{F}$ =0.92), whereas the free dye has a quantum yield of 0.4.

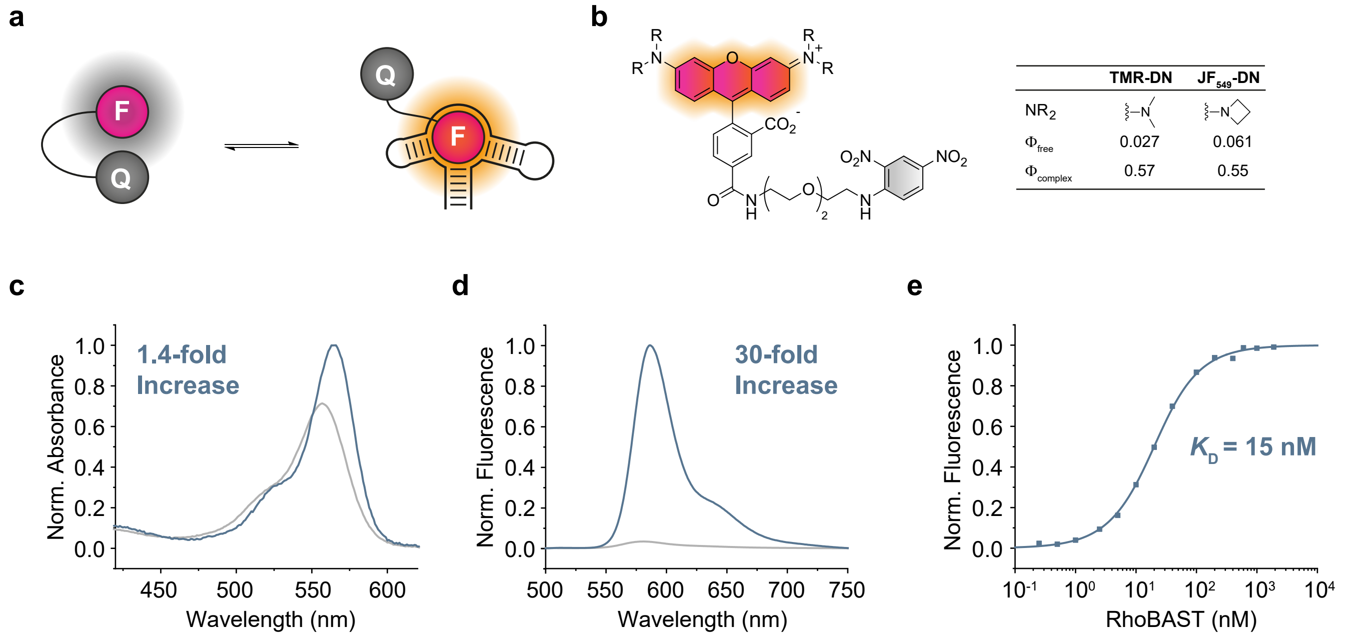


**Supplementary Figure N1: Properties of the fluorophore-quencher conjugate $JF_{549}$-DN. a)** Schematic illustration of the light-up mechanism of fluorophore-quencher conjugates. Binding to an aptamer leads to disruption of the fluorophore-quencher interaction and consequently a fluorescence increase. **b)** Chemical structure and fluorescence quantum yields of TMR-DN and $JF_{549}$-DN. **c)** Normalized absorption spectra of $JF_{549}$-DN in the presence (blue line) and absence (grey line) of RhoBAST. **d)** Normalized fluorescence emission spectra of $JF_{549}$-DN in the presence (blue line) and absence (grey line) of RhoBAST. Absorbance and fluorescence measurements were performed in ASB at 25 °C using 1 µM dye and 5 µM RhoBAST. **e)** Binding isotherm of $JF_{549}$-DN using RhoBAST. Measurements were performed in ASB supplemented with 0.05% Tween 20 at 25 °C using 10 nM probe.

### Supplementary Note 2:

#### Shifting the open-closed equilibrium by increasing the electrophilicity of the xanthene

The electrophilicity of the xanthene core can be increased by the introduction of fluorine substituents at the 2′- and 7′-position and/or the use of amine substituents bearing electron-withdrawing groups (Supplementary Fig. N2a).[3-7] Notably, the decreased electron density does not only lead to a shift of the open-closed equilibrium to the non-fluorescent spirolactone form, but also to an undesired hypsochromic shift of the excitation and emission maxima,[3] making the dye less suitable for excitation with a common 561 nm laser. However, due to the high brightness, photostability and structural similarity to the high-affinity binder 5C-JF$_{549}$ ($K_D$ = 35 nM), we decided to synthesize the fluorinated version 5C-JF$_{525}$ and test its applicability as a ligand for RhoBAST.[3]

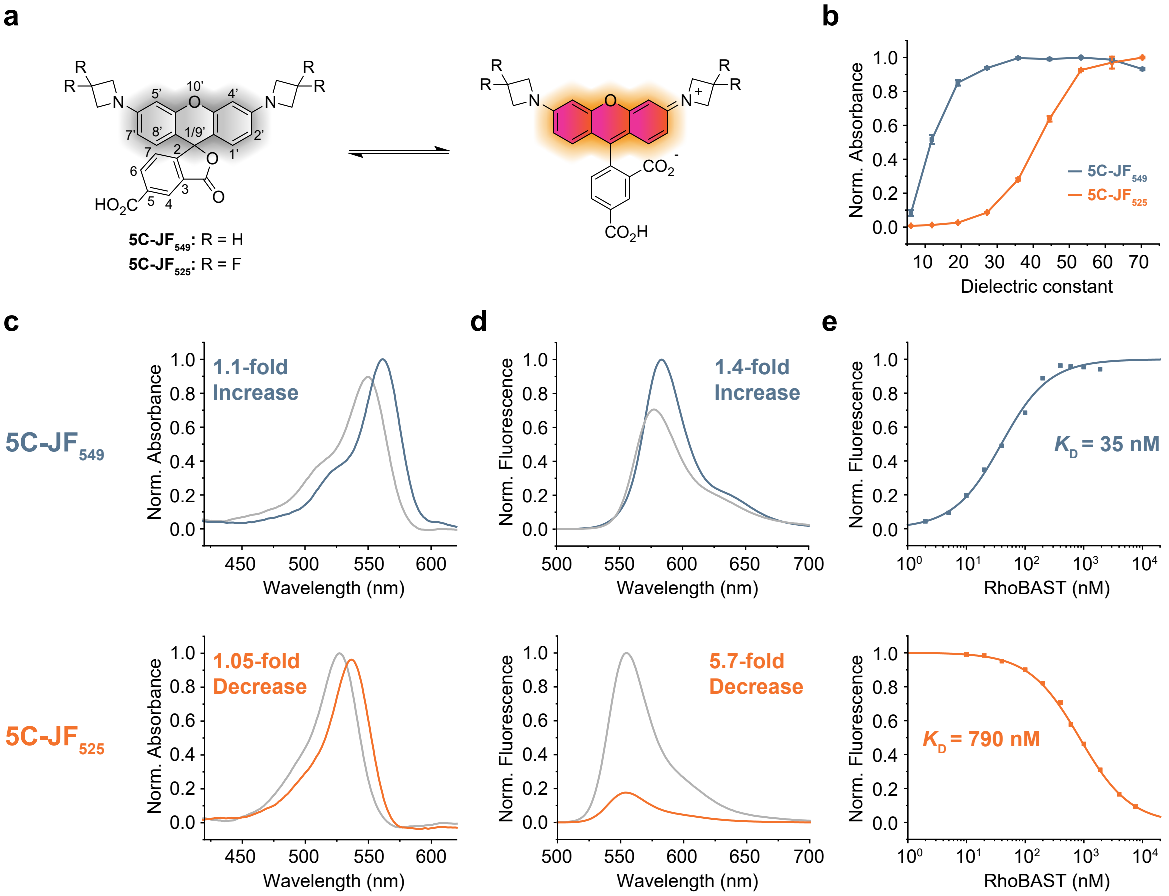


**Supplementary Figure N2. Properties of azetidinyl rhodamines. a)** Chemical structure and open-closed equilibrium of 5C-JF$_{549}$ and 5C-JF$_{525}$. **b)** Absorption of 5C-JF$_{549}$ (5 µM) and 5C-JF$_{525}$ (5 µM) as a function of the dielectric constant of the solvent mixture (water/dioxane). The absorbance was normalized to the highest measured point of each derivative. **c)** Normalized absorption spectra of 5C-JF$_{549}$ and 5C-JF$_{525}$ in the presence (colored lines) and absence (grey lines) of RhoBAST. **d)** Normalized fluorescence emission spectra of 5C-JF$_{549}$ and 5C-JF$_{525}$ in the presence (colored lines) and absence (grey lines) of RhoBAST. Absorbance and fluorescence spectra were measured in ASB at 25 °C using 1 µM dye and 5 µM RhoBAST. **e)** Binding isotherms of 5C-JF$_{549}$ and 5C-JF$_{525}$ using RhoBAST, measured in ASB supplemented with 0.05% Tween 20 at 25 °C using 10 nM probe.

First, the $D_{50}$ values of azetidinyl rhodamine 5C-JF$_{549}$ and 5C-JF$_{525}$ were measured to investigate the position of the open-closed equilibrium (Supplementary Fig. N2a,b). Compared to the non-fluorinated parent dye 5C-JF$_{549}$ ($D_{50}$ = 12), 5C-JF$_{525}$ displayed the desired shift of the equilibrium towards the spirolactone form ($D_{50}$ = 41). However, despite the structural similarity, 5C-$JF_{525}$ bound to RhoBAST with a substantially lower affinity ($K_D$ = 793 nM) than 5C-$JF_{549}$ ($K_D$ = 35 nM) and displayed a fluorescence decrease instead of the desired light-up (Supplementary Fig. N2c-e). A similar turn-off behavior was previously observed for the xanthene dyes 9-aminoacridine and oxazine 1, which possess a relatively electron-poor conjugated system, upon binding to sulforhodamine B-binding aptamer SRB-2, the predecessor to RhoBAST.[1] Consequently, we reasoned that reducing the electron density in the xanthene system would not lead to the desired advanced fluorogenic probes for the RhoBAST system.

### Supplementary Note 3:

**Chemical Synthesis**

All commercial reagents were purchased from suppliers Sigma-Aldrich, ABCR, Acros, TCI and Alfa Aesar and used without further purification. Moisture and/or oxygen-sensitive reactions were carried out under an argon atmosphere using standard Schlenk techniques.

Silica column chromatography was performed using silica gel (high-quality, pore size 60 Å, 40-63 µm particle size) purchased from Sigma-Aldrich. All fluorophores were purified *via* reverse phase High Performance Liquid Chromatography (HPLC) before use in live-cell or *in vitro* experiments. HPLC was performed on an AGILENT 1100 Series HPLC system equipped with a multi wavelength detector and a fraction collector using a Phenomenex Luna 5 µm C-18(2) 100 Å (250 x 21.2 mm) column. All purification runs were carried out with a constant flow rate of 6 ml/min using mixtures of water and acetonitrile (containing 0.1% TFA each) as solvent system. Appropriate fractions were freeze-dried using a lyophilizer Alpha 2-4 LDplus (Christ).

Nuclear Magnetic Resonance Spectroscopy (NMR) spectra were recorded on Varian Mercury Plus 300 or Mercury Plus 500 systems. Deuterated solvents were purchased from Euriso-Top. Chemical shifts ($\delta$) are given in ppm (with respect to tetramethylsilane) and coupling constants ($J$) in Hz. All recorded $^{1}H$ and $^{13}C$ spectra were referenced to the protio impurity or the $^{13}C$ signal of the deuterated solvent. $^{13}C$ measurements were recorded as APT spectra.

High resolution mass spectra (HR-MS) were measured on a Bruker micrOTOF QII-ESI system using sodium formiate as internal calibrant. The reported mass to charge ratios (m/z) refer to the isotopic peak with the highest intensity.

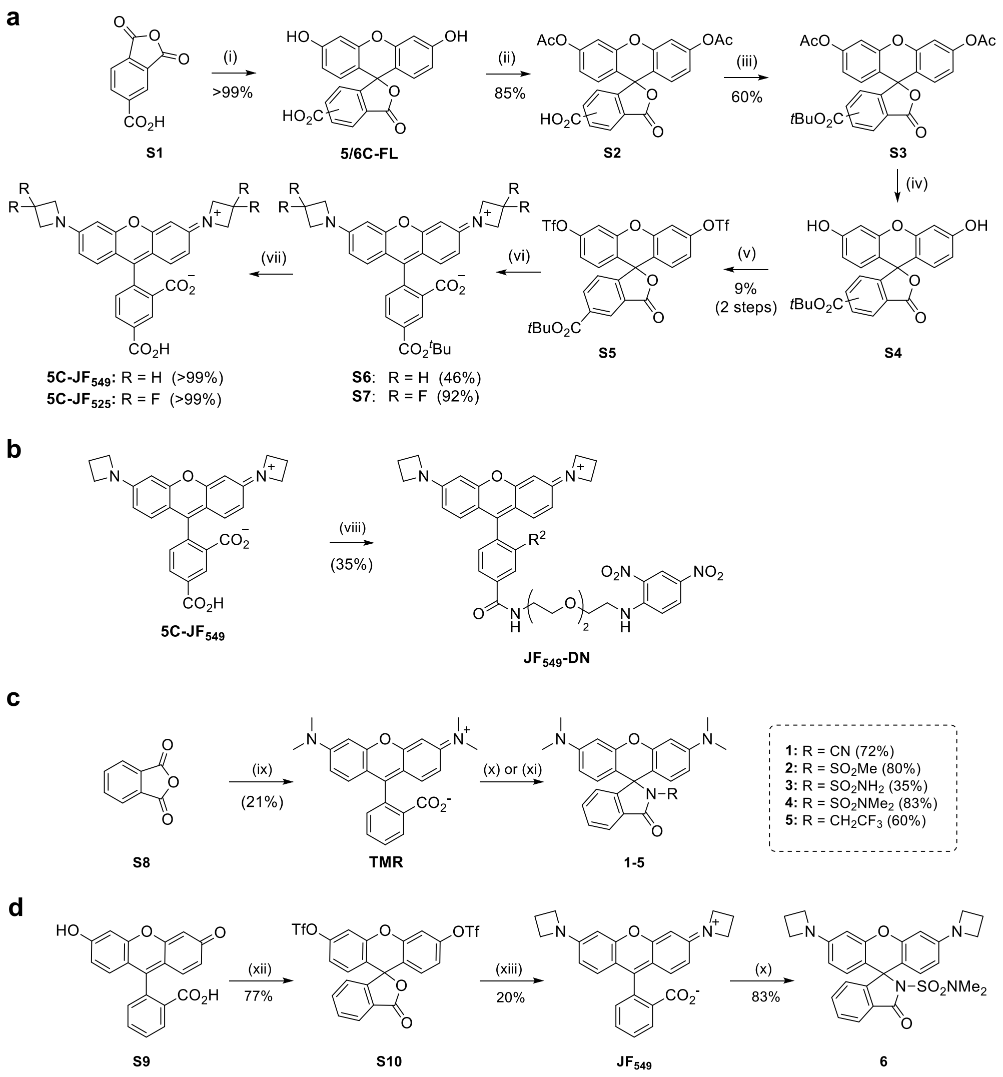


**Supplementary Figure N3: Synthesis of rhodamine dyes.** (i) Resorcinol, $MeSO_3H$; (ii) Acetic anhydride; (iii) Dimethylformamide di-tert-butyl acetal, toluene; (iv) NaOH, methanol, THF; (v) Trifluoromethane sulfonic anhydride, pyridine, DCM; (vi) Azetidine/3,3-difluoroazetidine hydrochloride, $Pd_2(dba)_3$, XPhos, $Cs_2CO_3$, dioxane; (vii) TFA, DCM. (viii) TSTU, **DN-NH₂**, DIPEA, DMF. (ix) 3-Dimethylaminophenol; (x) amine, EDC, DMAP (for **1**,**2**,**4**,**5**,**6**); (xi) $POCl_3$, amine, DIPEA (for **3**); (xii) $Tf_2O$, pyridine; (xiii) azetidine, $Pd_2dba_3$, XPhos, $Cs_2CO_3$.

**5/6-Carboxy fluorescein (5/6-C-FL)**

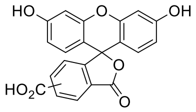

$C_{21}H_{12}O_7$
376.32 g/mol

Trimellitic anhydride (6.00 g, 31.2 mmol, 1.00 eq) and resorcinol (6.88 g, 62.5 mmol, 2.00 eq) were suspended in methanesulfonic acid (40 mL). The mixture was stirred at 120 °C for 2 h. The solution was poured into ice water (500 mL) and the resulting precipitate was isolated *via* filtration. The precipitate was washed with water (2 × 100 mL) and dissolved in aqueous 1.25 M NaOH (200 mL). The solution was reacidified with conc. HCl (~30 mL) to pH 2-3 and the resulting precipitate was isolated by filtration and washed with water (2 × 100 mL). Next, the residue was dried in an oven at 160 °C for 10 h to yield the product as an orange powder (11.7 g, >99%). The recorded spectrum was in accordance with literature.[8]

**$^1$H NMR** (300 MHz, $(CD_3)_2SO$): *δ* = 8.41 – 8.37 (m, 1H, 5-isomer), 8.29 (dd, *J* = 8.1, 1.5 Hz, 1H, 5-isomer), 8.22 (dd, *J* = 8.0, 1.3 Hz, 1H, 6-isomer), 8.14 – 8.07 (m, 1H, 6-isomer), 7.67 – 7.61 (m, 1H, 6-isomer), 7.42 – 7.35 (m, 1H, 5-isomer), 6.74 – 6.68 (m, 4H, 5-isomer, 6-isomer), 6.64 – 6.51 (m, 8H, 5-isomer, 6-isomer) ppm.

**5/6-Carboxy fluorescein diacetate (S2)**

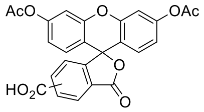

$C_{25}H_{16}O_9$
460.39 g/mol

First, 5/6-Carboxy fluorescein (1.50 g, 3.99 mmol, 1.00 eq) was suspended in $Ac_2O$ (10 mL) and the mixture was refluxed for 4 h. Next, water was added until a white precipitate was obtained. The supernatant was discarded and the obtained solid was dissolved in EtOAc (50 mL). The organic phase was washed with brine (20 mL), dried over magnesium sulfate, filtered and the solvent was removed under reduced pressure. The desired product was obtained as a light yellowish solid (1.41 g, 85%). The recorded spectrum was in accordance with literature.[9]

**$^1$H NMR** (300 MHz, $(CD_3)_2SO$): *δ* = 8.46 – 8.43 (m, 1H, 5-isomer), 8.32 (dd, *J* = 8.1, 1.5 Hz, 1H, 5-isomer), 8.27 (dd, *J* = 8.0, 1.3 Hz, 1H, 6-isomer), 8.17 (d, *J* = 7.8 Hz, 1H, 6-isomer), 7.84 (s, 1H, 6-isomer), 7.55 (d, *J* = 8.0 Hz, 1H, 5-isomer), 7.30 (d, *J* = 1.3 Hz, 3H, 5/6-isomer), 6.95 (m, 7H, 5/6-isomer), 2.29 (s, 12H, 5/6-isomer) ppm.

**5/6-*tert*-Butoxycarbonyl fluorescein diacetate (S3)**

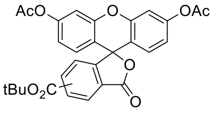

$C_{25}H_{16}O_9$
516.50 g/mol

A *Schlenk* flask was charged with 5/6-Carboxy fluorescein diacetate (1.20 g, 2.61 mmol, 1.00 eq) and evacuated/backfilled with Argon three times. The solid was suspended in anhydrous toluene (10 mL) and *N*,*N*-Dimethylformamide di-*tert*-butylacetal (2.50 mL, 2.12 g, 10.4 mmol, 4.00 eq) was added to the suspension. The mixture was stirred at 85 °C overnight. Next, the reaction mixture was diluted with toluene (50 mL) and washed with saturated aqueous $NaHCO_3$ solution (2 × 30 mL). The organic phase was dried over magnesium sulfate, filtered and the solvent removed under reduced pressure to yield **S3** as light yellowish solid (773 mg, 57%).

**$^1$H NMR** (300 MHz, $CDCl_3$): *δ* = 8.62 (dd, *J* = 1.4, 0.7 Hz, 1H, 5-isomer), 8.31 (dd, *J* = 8.0, 1.5 Hz, 1H, 5-isomer), 8.26 (dd, *J* = 8.0, 1.3 Hz, 1H, 6-isomer), 8.07 (dd, *J* = 8.0, 0.7 Hz, 1H, 6-isomer), 7.73 (dd, *J* = 1.2, 0.8 Hz, 1H, 6-isomer), 7.23 (m, 1H, 5-isomer), 7.13 – 7.10 (m, 4H, 5/6-isomer), 6.85 – 6.77 (m, 8H, 5/6-isomer), 2.32 (m, 12H, 5/6-isomer), 1.64 (s, 9H, 5-isomer), 1.56 (s, 9H, 5/6-isomer) ppm.

**$^{13}$C NMR** (75 MHz, $CDCl_3$): *δ* = 169.0, 168.9, 168.4, 168.4, 164.0, 164.0, 156.2, 152.8, 152.4, 152.3, 151.7, 151.7, 138.8, 136.4, 134.6, 131.4, 129.4, 129.2, 129.1, 129.0, 128.4, 126.8, 126.6, 125.4, 125.3, 125.2, 124.3, 118.0, 116.0, 115.9, 110.7, 110.6, 83.0, 82.6, 82.2, 81.9, 28.3, 28.1, 21.3 ppm.

**MS** (HR-ESI, pos): meas. *m/z* = 539.1313, calc. *m/z* = 539.1313 for $C_{25}H_{16}NaO_9$ [M+Na]$^+$.

**5-*tert*-Butoxycarbonyl fluorescein ditriflate (S5)**

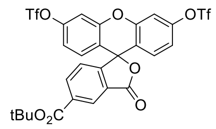


$C_{27}H_{18}F_6O_{11}S_2$
696.54 g/mol

First, 5/6-carboxyfluorescein diacetate *tert*-butyl ester (1.62 g, 3.14 mmol, 1.00 eq) was dissolved in a 1:1 mixture of THF and MeOH (25 mL) and aqueous 1 M NaOH (6 mL) was added. The mixture was stirred at room temperature for 1 h. The solution was acidified with aqueous 1 M HCl (7 mL) and diluted with water (30 mL). The aqueous phase was extracted with EtOAc (2 × 60 mL) and the combined organic layers were washed with brine (40 mL). The organic layer was dried over $MgSO_4$, filtered and the solvent removed *in vacuo*. The resulting orange solid was taken up in DCM (15 mL) and cooled to 0 °C. Pyridine (2.02 mL, 1.98 g, 25.1 mmol, 8.00 eq) was added to the suspension, followed by $Tf_2O$ (2.11 mL, 3.54 g, 12.6 mmol, 4.00 eq). The mixture was allowed to warm up to room temperature and was stirred for an additional 1 h. The resulting red solution was diluted with water (30 mL) and extracted with DCM (2 × 60 mL). The combined organic layers were dried over magnesium sulfate, filtered and put on celite. The crude mixture was purified by silica column chromatography (cyclohexane:EtOAc = 10:0 → 7:1). The isomerically pure product was obtained as an off-white solid (200 mg, 9%).

**$^1$H NMR** (500 MHz, $CDCl_3$): *δ* = 8.66 (s, 1H), 8.36 (dd, *J* = 8.0, 1.2 Hz, 1H), 7.31 (d, *J* = 2.4 Hz, 2H), 7.24 (d, *J* = 8.0 Hz, 1H), 7.03 (dd, *J* = 8.8, 2.4 Hz, 2H), 6.94 (d, *J* = 8.8 Hz, 2H), 1.64 (s, 9H) ppm.

**$^{13}$C NMR** (126 MHz, $CDCl_3$): *δ* = 167.8, 163.7, 155.4, 151.4, 150.5, 136.9, 135.3, 130.0, 127.2, 126.0, 123.9, 120.9, 118.9, 118.0, 111.0, 82.9, 80.3 28.3 ppm.

**MS** (HR-ESI, pos): meas. *m/z* = 719.0088, calc. *m/z* = 719.0087 for $C_{27}H_{18}F_6NaO_{11}S_2$ [M+Na]$^+$.

**General procedure for *Buchwald Hartwig* coupling of ditriflates (GP1)**

The following procedure is exemplary for the *Buchwald-Hartwig amination* of fluorescein ditriflates to the corresponding azetidine-substituted rhodamines:

A reaction vessel was charged with fluorescein ditriflate (1.00 eq), $Cs_2CO_3$ (2.80 eq), $Pd_2(dba)_3$ (0.20 eq) and 2-dicyclohexyl-phosphino-2′,4′,6′-triisopropylbiphenyl (XPhos, 0.30 eq), sealed, evacuated and backfilled with argon (3×). The solids were suspended in dry dioxane (35 mM solution). Azetidine or 3,3-difluoroazetidine hydrochloride (2.40 eq) was added and the mixture was stirred at 100 °C overnight. The mixture was diluted with MeOH and the solvent was evaporated under reduced pressure. The crude product was purified by silica column chromatography.

##### 5-*tert*-Butoxycarbonyl Janelia Fluor 549 (S6)

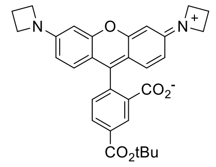


$C_{31}H_{30}N_2O_5$
510.59 g/mol

Rhodamine **S6** was synthesized from ditriflate **S5** and azetidine according to general procedure 1. The crude mixture was purified by silica column chromatography ($CHCl_3$:2 M $NH_3$ in MeOH = 10:0 → 9:1) and the product was obtained as violet solid (33.8 mg, 46%).

**$^1$H NMR** (300 MHz, $CDCl_3$): $\delta$ = 8.60 (s, 1H), 8.25 (dd, $J$ = 8.0, 1.3 Hz, 1H), 7.21 (d, $J$ = 8.0 Hz, 1H), 6.56 (d, $J$ = 8.6 Hz, 2H), 6.20 (d, $J$ = 1.9 Hz, 2H), 6.09 (dd, $J$ = 8.6, 2.0 Hz, 2H), 3.93 (t, $J$ = 7.3 Hz, 8H), 2.39 (p, $J$ = 7.2 Hz, 4H), 1.63 (s, 9H) ppm.

**$^{13}$C NMR** (75 MHz, $CDCl_3$): $\delta$ = 169.0, 164.5, 155.0, 154.0, 153.2, 135.3, 133.8, 129.1, 128.6, 126.9, 124.8, 108.2, 107.7, 97.4, 82.3, 70.7, 52.2, 28.3, 16.7 ppm.

**MS** (HR-ESI, pos): meas. $m/z$ = 511.2225, calc. $m/z$ = 511.2227 for $C_{31}H_{31}N_2O_5$ $[M+H]^+$.

##### 5-*tert*-Butoxycarbonyl Janelia Fluor 525 (S7)

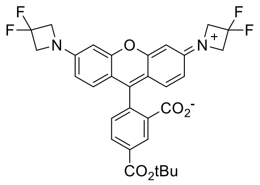


$C_{31}H_{26}F_4N_2O_5$
582.55 g/mol

Rhodamine **S7** was synthesized from ditriflate **S5** and 3,3-difluoroazetidine hydrochloride according to general procedure 1. The crude mixture was purified by silica column chromatography ($CHCl_3$:MeOH = 10:0 → 60:1) and the product was obtained as violet solid (80.5 mg, 92%).

**$^1$H NMR** (500 MHz, $CDCl_3$): $\delta$ = 8.65 – 8.54 (m, 1H), 8.28 (dd, $J$ = 8.0, 1.4 Hz, 1H), 7.20 (d, $J$ = 8.0 Hz, 1H), 6.61 (d, $J$ = 8.6 Hz, 2H), 6.30 (d, $J$ = 2.3 Hz, 2H), 6.16 (dd, $J$ = 8.6, 2.3 Hz, 2H), 4.25 (t, $^3J_{HF}$ = 11.7 Hz, 8H), 1.64 (s, 9H) ppm.

**$^{19}$F NMR** (282 MHz, $CDCl_3$) $\delta$ = −99.5 (p, $^3J_{HF}$ = 11.7 Hz) ppm.

**$^{13}$C NMR** (126 MHz, $CDCl_3$): $\delta$ = 168.8, 164.3, 156.6, 152.5, 151.5 (t, $^4J_{CF}$ = 2.77 Hz), 136.0, 134.2, 129.3, 127.5, 126.6, 124.1, 115.7 (t, $^1J_{CF}$ = 275 Hz), 109.0, 108.9, 99.4, 84.2, 82.5, 63.4 (t, $^2J_{CF}$ = 26.5 Hz), 28.3 ppm.

**MS** (HR-ESI, pos): meas. $m/z$ = 605.1670; calc. $m/z$ = 605.1770 for $C_{31}H_{26}F_4N_2NaO_5$ $[M+Na]^+$.

**General procedure for the deprotection of *tert*-butyl ester of rhodamines (GP2)**

The following procedure is exemplary for the deprotection of *tert*-butyl esters of azetidine-substituted rhodamines.

The rhodamine was dissolved in DCM (1.5 mL) and TFA (150 µL) was added. The mixture was stirred at room temperature for 6 h. The resulting solution was diluted with benzene upon which a dark solid precipitated. The solvents were removed under reduced pressure and the resulting crude product was co-evaporated with MeOH (3×).

**5-Carboxy Janelia Fluor 549 (5C-JF$_{549}$)**

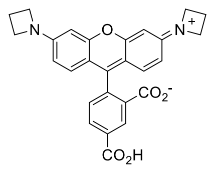


$C_{27}H_{22}N_2O_5$
454.48 g/mol

The deprotected **5C-JF$_{549}$** fluorophore was synthesized from **S6** according to general procedure 2. The TFA salt of the product was obtained as dark violet solid (32.9 mg, 98%).

**$^1$H NMR** (500 MHz, $CD_3OD$): $\delta$ = 8.88 (d, *J* = 1.4 Hz, 1H), 8.39 (dd, *J* = 7.9, 1.6 Hz, 1H), 7.48 (d, *J* = 7.9 Hz, 1H), 7.06 (d, *J* = 9.2 Hz, 2H), 6.60 (dd, *J* = 9.2, 2.1 Hz, 2H), 6.53 (d, *J* = 2.0 Hz, 2H), 4.30 (t, *J* = 7.6 Hz, 8H), 2.56 (p, *J* = 7.6 Hz, 4H) ppm.

**$^{13}$C NMR** (126 MHz, $CD_3OD$): $\delta$ = 168.2, 167.8, 160.5, 158.8, 158.1, 139.3, 134.4, 134.1, 133.8, 133.3, 132.2, 131.8, 114.7, 113.6, 95.1, 52.8, 16.8 ppm.

**MS** (HR-ESI, pos): meas. *m/z* = 455.1612, calc. *m/z* = 455.1601 for $C_{27}H_{22}N_2O_5$ $[M+H]^+$.

**5-Carboxy Janelia Fluor 525 (5C-JF$_{525}$)**

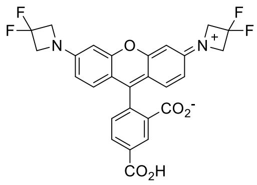


$C_{27}H_{15}F_4N_2O_5$
526.44 g/mol

The deprotected **5C-JF$_{525}$** fluorophore was synthesized from **S7** according to general procedure 2. The TFA salt of the product was obtained as dark violet solid (86.0 mg, 98%).

**$^1$H NMR** (500 MHz, $CD_3OD$): $\delta$ = 8.92 (d, *J* = 1.3 Hz, 1H), 8.45 (dd, *J* = 7.9, 1.6 Hz, 1H), 7.54 (d, *J* = 7.9 Hz, 1H), 7.20 (d, *J* = 9.1 Hz, 2H), 6.81 (d, *J* = 2.1 Hz, 2H), 6.78 (dd, *J* = 9.1, 2.2 Hz, 2H), 4.69 (t, $^3J_{HF}$ = 11.6 Hz, 8H) ppm.

**$^{19}$F NMR** (282 MHz, $CD_3OD$) $\delta$ = −77.04 (s), −102.55 (p, $^3J_{HF}$ = 11.7 Hz) ppm.

**$^{13}$C NMR** (126 MHz, $CD_3OD$): $\delta$ = 167.8, 167.3, 158.7, 157.5 (t, $^4J_{CF}$ = 4.16 Hz), 134.8, 134.5, 133.2, 132.6, 132.5, 131.5, 116.5 (t, $^1J_{CF}$ = 285 Hz), 115.6, 115.0, 97.5, 64.2 (t, $^2J_{CF}$ = 29.0 Hz) ppm.

**MS** (HR-ESI, pos): meas. *m/z* = 527.1224; calc. *m/z* = 527.1225 [*M*+H]$^+$.

##### ***N*-(2-(2-(2-aminoethoxy)ethoxy)ethyl)-2,4-dinitroaniline (DN-NH$_2$)**

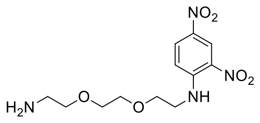

$C_{12}H_{18}N_4O_6$
314.30 g/mol

First, 2,2′-(Ethylenedioxy)bis(ethylamine) (8.58 mL, 58.8 mmol, 10.0 eq) was dissolved in DCM (40 mL). The resulting solution was cooled to 0 °C and 1-fluoro-2,4-dinitrobenzene (675 µL, 5.88 mmol, 1.00 eq) was added dropwise under stirring. The reaction mixture was allowed to warm up to room temperature and was stirred for 45 min. Next, water (200 mL) was added and the organic phase was recovered. The organic phase was extracted with 0.1 M aqueous HCl (200 mL). The pH of the aqueous phase was adjusted to pH = 12 using 1 M aqueous NaOH (~20 mL) and the aqueous layer was extracted with DCM (2 × 200 mL). The combined organic phases were washed with brine (100 mL), dried over $MgSO_4$, filtered and the solvent was removed under reduced pressure. The product was obtained as a brown oil (1.25 g, 74%). The obtained $^1$H NMR spectrum was in accordance with the cited literature.[10]

**$^1$H NMR** (300 MHz, $CDCl_3$): *δ* = 9.14 (d, *J* = 2.6 Hz, 1H), 8.81 (s, 1H), 8.27 (dd, *J* = 9.5, 2.5 Hz, 1H), 6.94 (d, *J* = 9.5 Hz, 1H), 3.84 (t, *J* = 5.2 Hz, 2H), 3.75 – 3.64 (m, 4H), 3.60 (q, *J* = 5.1 Hz, 2H), 3.53 (t, *J* = 5.2 Hz, 2H), 2.89 (s, 2H), 1.65 (s, 2H) ppm.

##### **5-(((2-(2-((2,4-dinitrophenyl)amino)ethoxy)ethoxy)methyl)carbamoyl) Janelia Fluor 549 (JF$_{549}$-DN)**

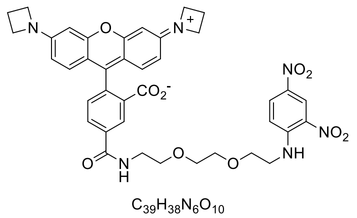

$C_{39}H_{38}N_6O_{10}$
750.77 g/mol

A *Schlenk* flask was charged with 5C-JF$_{549}$ (30.0 mg, 66.0 µmol, 1.00 eq) and TSTU (33.8 mg, 112 µmol, 1.70 eq), evacuated and backfilled with argon (3×). The solids were dissolved in anhydrous DMF (1 mL) and DIPEA (34.5 µL, 198 µmol, 5.00 eq) was added. The mixture was stirred at room temperature for 1.5 h. Next, DN-NH$_2$ (24.9 mg, 19.2 µmol, 1.20 eq) was dissolved in anhydrous DMF (0.5 mL) and added to the mixture. The reaction was stirred at room temperature overnight. The solvent was removed under reduced pressure and the crude product was purified by preparative reversed-phase HPLC (48 → 60% MeCN + 0.1% TFA in 40 min, $R_t$ = 25 min) and the product was obtained as violet solid (16.9 mg, 35%).

**$^1$H NMR** (500 MHz, $CD_3OD$): *δ* = 8.88 (d, *J* = 2.7 Hz, 1H), 8.70 (d, *J* = 1.6 Hz, 1H), 8.21 (dd, *J* = 7.9, 1.7 Hz, 1H), 8.18 (dd, *J* = 9.6, 2.7 Hz, 1H), 7.42 (d, *J* = 7.9 Hz, 1H), 7.15 (d, *J* = 9.6 Hz, 1H), 7.01 (d, *J* = 9.2 Hz, 2H), 6.56 (dd, *J* = 9.2, 2.0 Hz, 2H), 6.50 (d, *J* = 2.0 Hz, 2H), 4.29 (t, *J* = 7.6 Hz, 8H), 3.84 (t, *J* = 5.2 Hz, 2H), 3.77 – 3.70 (m, 6H), 3.65 (q, *J* = 5.7 Hz, 4H), 2.56 (p, *J* = 7.6 Hz, 4H) ppm.

**$^{13}$C NMR** (126 MHz, $CD_3OD$): *δ* = 168.2, 167.3, 160.2, 158.7, 158.0, 149.8, 138.2, 137.6, 137.0, 132.8, 132.3, 132.1, 131.9, 131.4, 131.3, 131.0, 124.6, 116.1, 114.6, 113.6, 95.1, 71.7, 71.5, 70.6, 69.8, 52.9, 44.2, 41.2, 16.8 ppm.

**MS** (HR-ESI, pos): meas. *m/z* = 751.2713, calc. *m/z* = 751.2722 for $C_{39}H_{39}N_6O_{10}$ [M+H]$^+$.

**Tetramethyl rhodamine (TMR)**

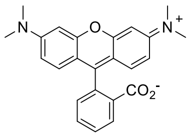

$C_{24}H_{22}N_2O_3$
386.45 g/mol

Phthalic anhydride (300 mg, 2.03 mmol, 1.00 eq) and 3-dimethylaminophenol (556 mg, 4.05 mmol, 2.00 eq) were placed inside a flask and stirred at 150 °C for 24 h. After allowing the mixture to cool to room temperature, the resulting solid was taken up in $CHCl_3$/MeOH (10:1) and purified by silica column chromatography ($CHCl_3$:MeOH = 10:1 → 5:1). The product was obtained as dark violet solid (164 mg, 21%). The measured spectrum was in accordance with literature.[11]

**$^1$H NMR** (300 MHz, $CD_3OD$): *δ* = 8.15 – 8.07 (m, 1H), 7.71 – 7.59 (m, 2H), 7.30 – 7.21 (m, 3H), 6.99 (dd, *J* = 9.5, 2.5 Hz, 2H), 6.89 (d, *J* = 2.5 Hz, 2H), 3.26 (s, 12H) ppm.

**MS** (HR-ESI, pos): meas. *m/z* = 387.1714, calc. *m/z* = 387.1703 for $C_{24}H_{23}N_2O_3$ $[M+H]^+$.

**General procedure for the amide coupling reaction of 3-carboxy group of rhodamines (GP3)**

The following procedure is exemplary for the amide coupling of the 3-carboxy functionality of rhodamine derivatives with the corresponding amines to obtain the respective cyclic lactams.

A reaction vessel was charged with the rhodamine derivative (1.00 eq), EDC hydrochloride (8.00 eq), DMAP (8.00 eq) and corresponding amine (1.20 eq), evacuated and backfilled with argon (3×). The solids were dissolved in anhydrous DCM (25 mM solution) and stirred at 60 °C overnight. The solvent was removed under reduced pressure and the crude product was purified by silica column chromatography.

**3-(Cyanocarbamoyl) tetramethyl rhodamine (1)**

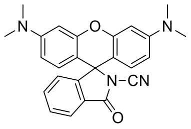

$C_{25}H_{22}N_4O_2$
410.48 g/mol

Rhodamine **1** was synthesized from TMR and cyanamide according to general procedure 3. The obtained crude mixture was purified by silica column chromatography ($CHCl_3$:MeOH = 10:0 → 50:1) and the product was obtained as violet solid (42.7 mg, 80%).

**$^1$H NMR** (500 MHz, $CDCl_3$): *δ* = 8.00 (d, *J* = 7.6 Hz, 1H), 7.63 (m, 1H), 7.58 - 7.54 (m, 1H), 7.14 (d, *J* = 7.7 Hz, 1H), 6.58 (d, *J* = 8.9 Hz, 2H), 6.47 (d, *J* = 2.5 Hz, 2H), 6.41 (dd, *J* = 8.9, 2.5 Hz, 2H), 2.99 (s, 12H) ppm.

**$^{13}$C NMR** (126 MHz, $CDCl_3$): $\delta$ = 166.6, 153.1, 152.6, 152.2, 135.7, 129.5, 128.3, 126.7, 124.9, 124.5, 109.3, 106.9, 104.3, 99.0, 68.8, 40.3 ppm.

**MS** (HR-ESI, pos): meas. *m/z* = 411.1814 calc. *m/z* = 411.1816 $[M+H]^+$.

**3-((Methylsulfonyl)carbamoyl) tetramethyl rhodamine (2)**

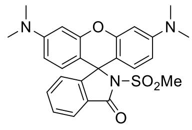


$C_{25}H_{25}N_3O_4S$
463.55 g/mol

Rhodamine **2** was synthesized from TMR and methanesulfonamide according to general procedure 3. The obtained crude mixture was purified by silica column chromatography ($CHCl_3$:MeOH = 10:0 → 50:1) and the product was obtained as violet solid (43.0 mg, 72%).

**$^1$H NMR** (500 MHz, $CDCl_3$): $\delta$ = 7.98 (d, *J* = 7.6 Hz, 1H), 7.58 (m, 1H), 7.52 (m, 1H), 7.08 (d, *J* = 7.7 Hz, 1H), 6.58 (d, *J* = 8.8 Hz, 2H), 6.47 (d, *J* = 2.5 Hz, 2H), 6.36 (dd, *J* = 8.8, 2.5 Hz, 2H), 2.97 (s, 12H), 2.93 (s, 3H) ppm.

**$^{13}$C NMR** (126 MHz, $CDCl_3$): $\delta$ = 167.5, 153.7, 153.1, 151.6, 135.3, 129.1, 128.2, 128.1, 124.7, 124.1, 108.5, 106.8, 99.1, 69.2, 42.1, 40.4 ppm.

**MS** (HR-ESI, pos): meas. *m/z* = 486.1417 calc. *m/z* = 486.1458 for $C_{25}H_{25}N_3NaO_4S$ $[M+Na]^+$.

**3-((*N,N*-Dimethylsulfamoyl)carbamoyl) tetramethyl rhodamine / SpyRho (4)**

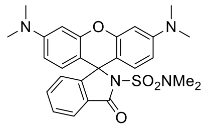


$C_{26}H_{28}N_4O_4S$
492.59 g/mol

SpyRho (**4**) was synthesized from TMR and *N*,*N*-dimethylsulfamide according to general procedure 3. The obtained crude mixture was purified by silica column chromatography ($CHCl_3$:MeOH = 10:0 → 50:1) and the product was obtained as violet solid (53.2 mg, 83%).

**$^1$H NMR** (500 MHz, $CDCl_3$): $\delta$ = 7.93 (d, *J* = 7.5 Hz, 1H), 7.57 – 7.48 (m, 2H), 7.06 (d, *J* = 7.6 Hz, 1H), 6.58 (d, *J* = 8.8 Hz, 2H), 6.48 (s, 2H), 6.35 (d, *J* = 8.4 Hz, 2H), 2.96 (s, 12H), 2.72 (s, 6H) ppm.

**$^{13}$C NMR** (126 MHz, $CDCl_3$): $\delta$ = 167.5, 154.0, 153.4, 151.5, 134.7, 128.8, 128.8, 128.4, 124.9, 123.7, 108.2, 107.6, 99.0, 68.9, 40.4, 38.0 ppm.

**MS** (HR-ESI, pos): meas. *m/z* = 493.1915; calc. *m/z* = 493.1904 for $C_{26}H_{29}N_4O_4S$ $[M+H]^+$.

**3-((2,2,2-Trifluoroethyl)carbamoyl) tetramethyl rhodamine (5)**

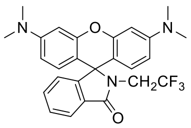

$C_{26}H_{24}F_3N_3O_2$
467.49 g/mol

Rhodamine **5** was synthesized from TMR and 2,2,2-trifluorethylamine according to general procedure 3. The obtained crude mixture was purified by silica column chromatography ($CHCl_3$:MeOH = 10:0 → 100:1) and the product was obtained as violet solid (36.5 mg, 60%).

**$^1$H NMR** (500 MHz, $CDCl_3$): $\delta$ = 7.95 (dd, $J$ = 5.8, 2.7 Hz, 1H), 7.52 – 7.43 (m, 2H), 7.06 (dd, $J$ = 5.7, 2.4 Hz, 1H), 6.49 – 6.41 (m, 4H), 6.35 (dd, $J$ = 8.8, 2.5 Hz, 2H), 3.72 (q, $^3J_{HF}$ = 9.3 Hz, 2H), 2.97 (s, 12H) ppm.

**$^{19}$F NMR** (282 MHz, $CDCl_3$) $\delta$ = −68.4 (t, $^3J_{HF}$ = 9.4 Hz) ppm.

**$^{13}$C NMR** (126 MHz, $CDCl_3$): $\delta$ = 169.4, 154.1, 153.1, 151.6, 133.5, 129.5, 128.9, 128.5, 124.2, 123.8 (q, $J_{CF}$ = 281 Hz)*, 123.5, 123.4, 109.0, 105.8, 98.7, 98.7, 65.6, 41.8*, 40.4 ppm.

**Note:** Highlighted peaks (*) were only detected in HSQC or HMBC experiments.

**MS** (HR-ESI, pos): meas. *m/z* = 468.1894; calc. *m/z* = 468.1893 for $C_{26}H_{25}F_3N_3O_2$ $[M+H]^+$.

**3-(Sulfamoylcarbamoyl) tetramethyl rhodamine (3)**

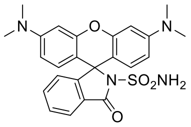

$C_{24}H_{24}N_4O_4S$
464.549 g/mol

A heat-dried *Schlenk* flask was charged with TMR (55.0 mg, 142 µmol, 1.00 eq), evacuated and backfilled with argon (3×). The solid was dissolved in anhydrous DCM (2 mL) and $POCl_3$ (266 µL, 436 mg, 2.85 mmol, 20.0 eq) was added dropwise. The solution was stirred at 60 °C for 3 h and the solvent was removed under reduced pressure to give the crude carboxylic acid chloride as a dark violet solid.

Next, sulfamide (136.8 mg, 1.42 mmol, 10.0 eq) and anhydrous DIPEA (372 µL, 276 mg, 2.13 mmol, 15.0 eq) were dissolved in anhydrous MeCN (4 mL) and added to the crude acyl chloride. The mixture was stirred at 70 °C for 1.5 h. The solvent was evaporated and the residue was taken up in chloroform (30 mL). The organic phase was washed with water (10 mL), dried over magnesium sulfate and filtered. After evaporation of the solvent, the crude product was purified by silica column chromatography ($CHCl_3$:MeOH = 10:0 → 50:1). The product was obtained as violet solid (23.1 mg, 35%)

**$^1$H NMR** (500 MHz, $(CD_3)_2SO$): $\delta$ = 7.89 (d, $J$ = 7.3 Hz, 1H), 7.62 – 7.58 (m, 1H), 7.56 (s, 2H), 7.55 – 7.52 (m, 1H), 6.92 (d, $J$ = 7.6 Hz, 1H), 6.51 – 6.47 (m, 2H), 6.40 (m, 4H), 2.91 (s, 12H) ppm.

**$^{13}$C NMR** (126 MHz, $(CD_3)_2SO$): $\delta$ = 166.2, 153.8, 151.9, 150.9, 134.8, 128.0, 127.2, 124.1, 123.1, 109.5, 108.5, 107.1, 98.3, 67.4, 39.9 ppm.

**MS** (HR-ESI, pos): meas. m/z = 487.1437; calc. m/z = 487.1410 for $C_{24}H_{24}N_4NaO_4S$ $[M+Na]^+$.

**Fluorescein ditriflate (S10)**

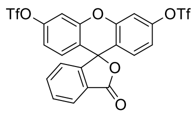


$C_{22}H_{10}F_6O_9S_2$
596.42 g/mol

A reaction vessel was charged with fluorescein (100 mg, 299 µmol, 1.00 eq), evacuated and backfilled with argon (3×). The starting material was suspended in anhydrous DCM (1.5 mL) and cooled to 0 °C. Next, Pyridine (193 µL, 189 mg, 2.39 mmol, 8.00 eq) was added followed by the dropwise addition of $Tf_2O$ (201 µL, 338 mg, 1.20 mmol, 4.00 eq). The solution was allowed to warm up to room temperature and was stirred for 6 h. The reaction was quenched with water and the obtained mixture was diluted with DCM (30 mL). The reaction mixture was washed with saturated $CuSO_4$ solution (25 mL) and brine (25 mL). The combined organic layers were dried over magnesium sulfate, filtered, and the solvent was removed under reduced pressure. The crude product was purified by silica column chromatography (cyclo-hexane:EtOAc = 10:0 → 3:1). The product was obtained as colorless solid (137 mg, 70%). The recorded spectrum was in accordance with literature.[12]

**$^1$H NMR** (300 MHz, $CDCl_3$): *δ* = 8.13 – 8.05 (m, 1H), 7.72 (m, 2H), 7.30 (d, *J* = 2.3 Hz, 2H), 7.22 – 7.16 (m, 1H), 7.08 – 6.92 (m, 4H) ppm.

**MS** (HR-ESI, pos): meas. *m/z* = 618.9574, calc. *m/z* = 618.9563 for $C_{22}H_{10}F_6NaO_9S_2$ $[M+Na]^+$.

**Janelia Fluor 549 ($JF_{549}$)**

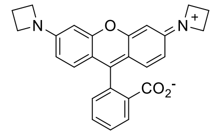


$C_{26}H_{22}N_2O_3$
410.47 g/mol

A reaction vessel was charged with fluorescein ditriflate (137 mg, 230 µmol, 1.00 eq), $Cs_2CO_3$ (210 mg, 643 µmol, 2.80 eq), $Pd_2(dba)_3$ (42.1 mg, 45.9 µmol, 0.20 eq) and 2-dicyclohexyl-phosphino-2′,4′,6′-triisopropylbiphenyl (XPhos; 32.9 mg, 68.9 µmol, 0.30 eq), sealed, evacuated and backfilled with argon (3×). The solids were suspended in dry dioxane (2 mL). Azetidine (31.5 mg, 551 µmol, 2.40 eq) was added and the mixture was stirred at 100 °C overnight. The mixture was diluted with MeOH and the solvent was evaporated under reduced pressure. The crude product was purified by silica column chromatography ($CHCl_3$:2 M $NH_3$ in MeOH = 10:0 → 9:1). The product was obtained as violet solid (19.0 mg, 20%). The recorded spectrum was in accordance with literature.[13]

**$^1$H NMR** (300 MHz, $CDCl_3$): *δ* = 8.00 (d, *J* = 7.6 Hz, 1H), 7.68 – 7.53 (m, 2H), 7.17 (d, *J* = 7.9 Hz, 1H), 6.57 (d, *J* = 8.6 Hz, 2H), 6.21 (d, *J* = 2.2 Hz, 2H), 6.09 (dd, *J* = 8.6, 2.3 Hz, 2H), 3.92 (t, *J* = 7.3 Hz, 8H), 2.38 (p, *J* = 7.5 Hz, 4H) ppm.

**MS** (HR-ESI, pos): meas. *m/z* = 411.1718, calc. *m/z* = 411.1703 for $C_{26}H_{23}N_2O_3$ $[M+H]^+$.

**3-((*N,N*-Dimethylsulfamoyl)carbamoyl) Janelia Fluor 549 (6)**

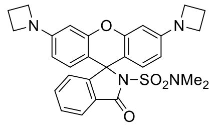


$C_{28}H_{28}N_4O_4S$
516.62 g/mol

A reaction vessel was charged with $JF_{549}$ (18.0 mg, 43.9 µmol, 1.00 eq), EDC hydrochloride (67.3 mg, 351 µmol, 8.00 eq), DMAP (42.9 mg, 351 µmol, 8.00 eq) and *N*,*N*-dimethylsulfamide (6.53 mg, 52.6 µmol, 1.20 eq) evacuated and backfilled with argon (3×). The solids were dissolved in anhydrous DCM (2 mL) and stirred at 60 °C overnight. The solvent was removed under reduced pressure and the crude product was purified by silica column chromatography ($CHCl_3$:MeOH = 10:0 → 50:1). The product was obtained as violet solid (18.9 mg, 83%).

**$^1$H NMR** (500 MHz, $CDCl_3$): *δ* = 7.92 (d, *J* = 7.4 Hz, 1H), 7.55 (m, 1H), 7.49 (m, 1H), 7.05 (d, *J* = 7.5 Hz, 1H), 6.53 (d, *J* = 8.5 Hz, 2H), 6.23 – 6.15 (m, 2H), 6.07 – 5.97 (m, 2H), 3.89 (t, *J* = 7.2 Hz, 8H), 2.71 (s, 6H), 2.36 (p, *J* = 7.2 Hz, 4H) ppm.

**$^{13}$C NMR** (126 MHz, $CDCl_3$): *δ* = 167.4, 154.1, 153.2, 153.0, 134.8, 128.8, 128.8, 128.4, 124.9, 123.7, 108.2, 107.1, 97.9, 69.0, 52.2, 38.0, 16.8 ppm.

**MS** (HR-ESI, pos): meas. *m/z* = 517.1900; calc. *m/z* = 517.1904 for $C_{28}H_{29}N_4O_4S$ $[M+H]^+$.

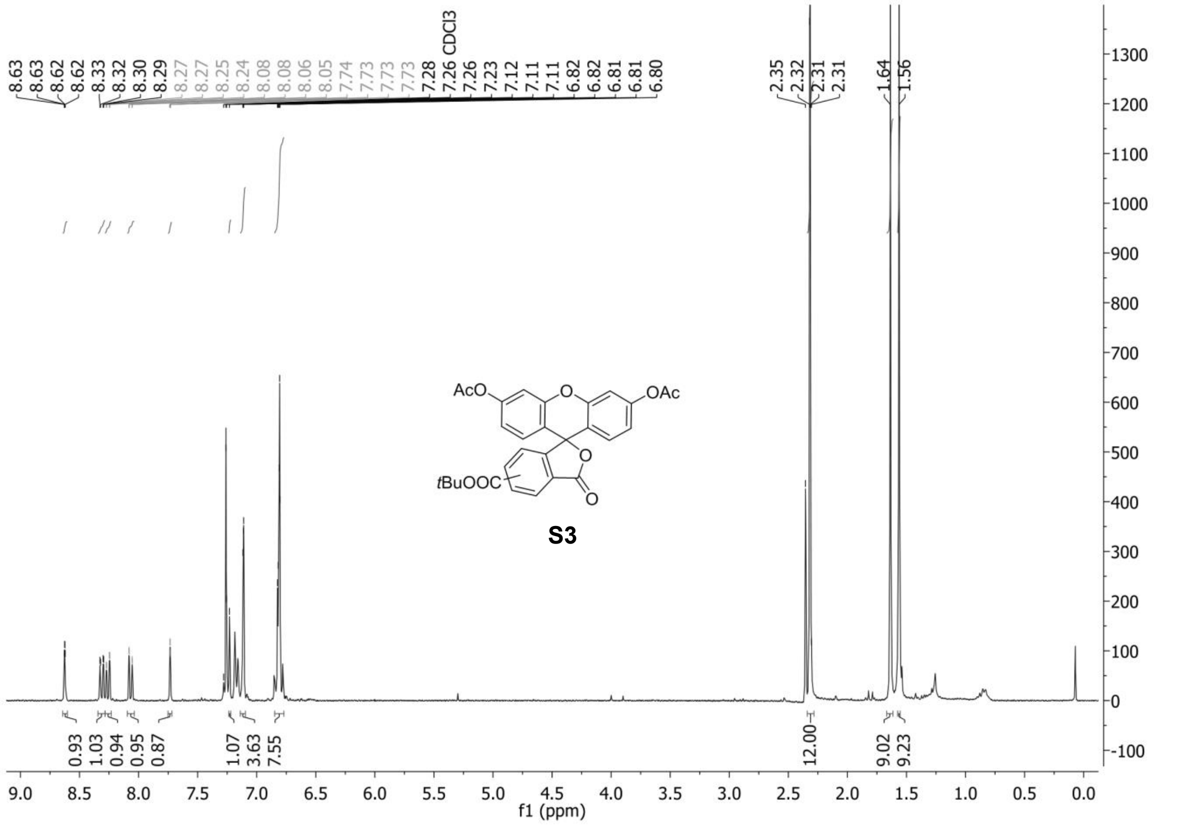
8.63
8.63
8.62
8.62
8.33
8.32
8.30
8.29
8.27
8.27
8.25
8.24
8.08
8.08
8.06
8.05
7.74
7.73
7.73
7.73
7.28
7.26 CDCl3
7.26
7.23
7.12
7.11
7.11
6.82
6.82
6.81
6.81
6.80
2.35
2.32
2.31
2.31
1.64
1.56
AcO
OAc
O
O
O
tBuOOC
S3
0.93
1.03
0.94
0.95
0.87
1.07
3.63
7.55
12.00
9.02
9.23
f1 (ppm)

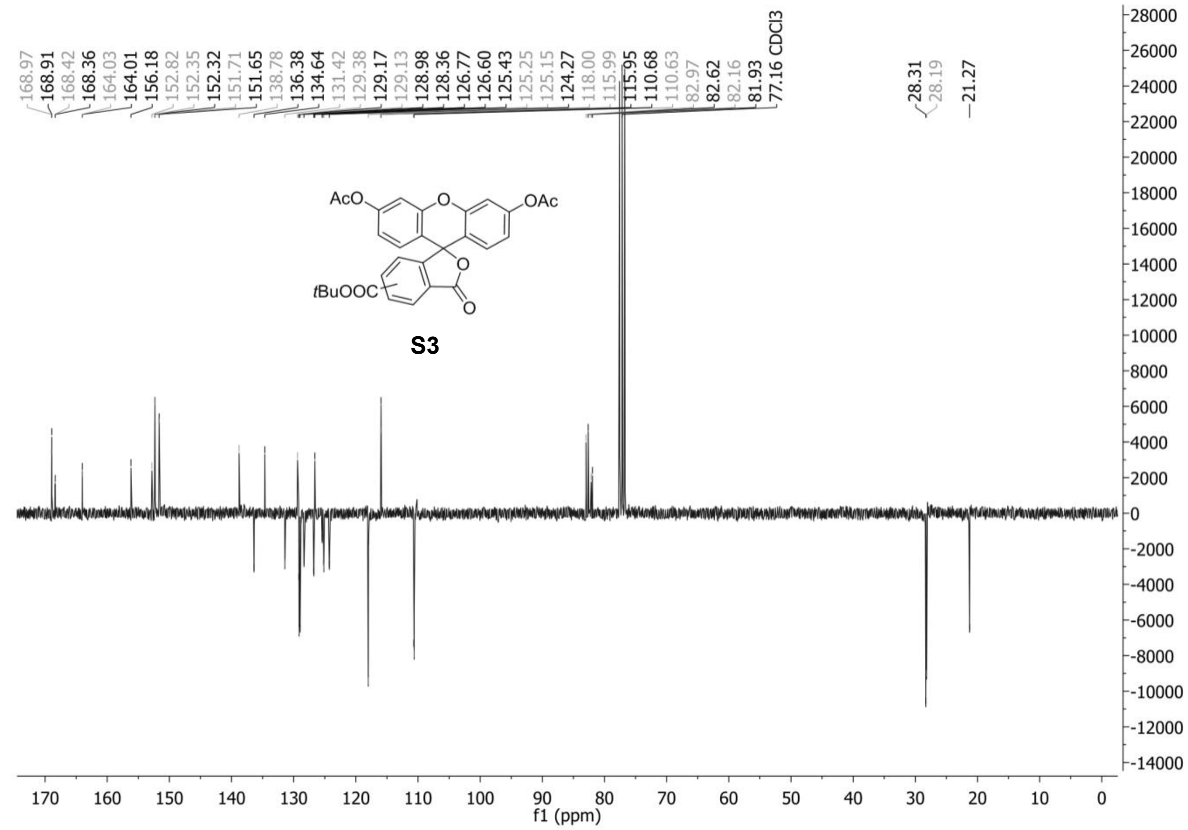
168.97
168.91
168.42
168.36
164.03
164.01
156.18
152.82
152.35
152.32
151.71
151.65
138.78
136.38
134.64
131.42
129.38
129.17
129.13
128.98
128.36
126.77
126.60
125.43
125.25
125.15
124.27
118.00
115.99
115.95
110.68
110.63
82.97
82.62
82.16
81.93
77.16 CDCl3
28.31
28.19
21.27
AcO
OAc
O
O
O
tBuOOC
S3
f1 (ppm)

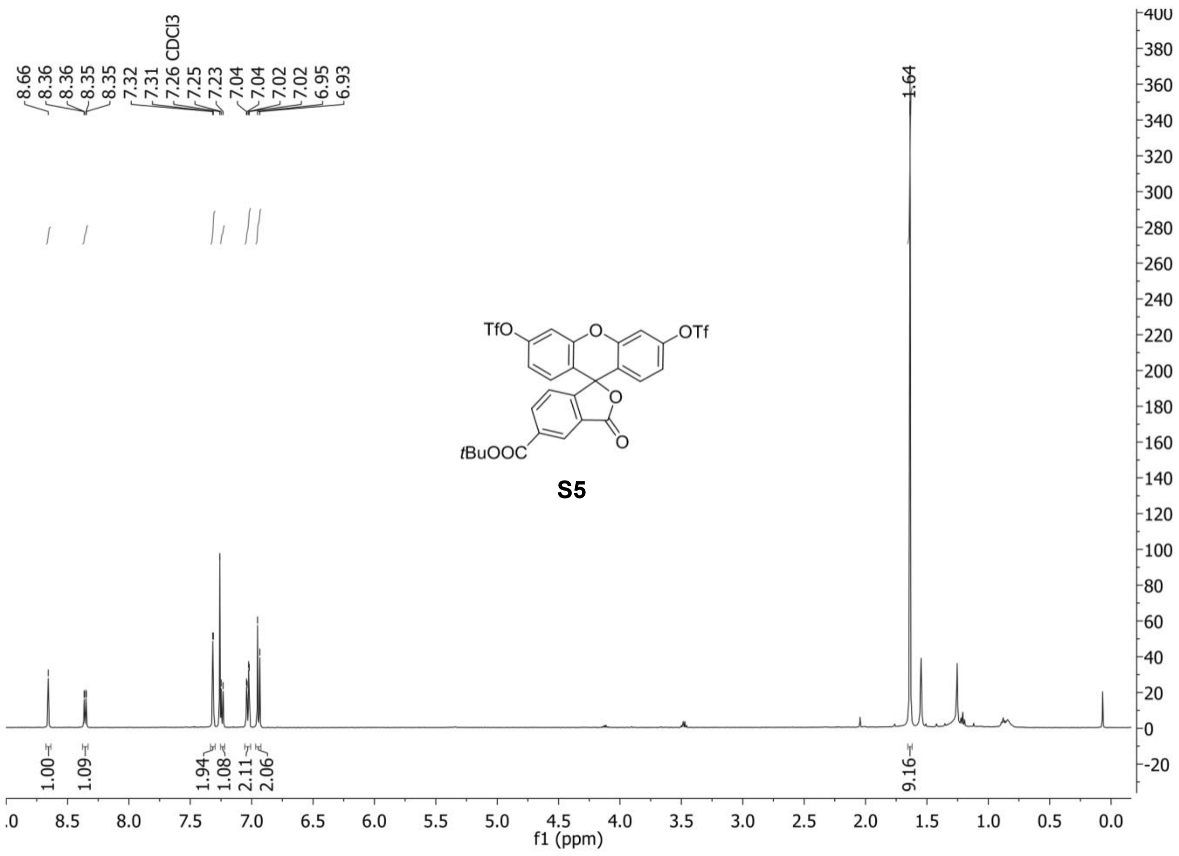
8.66
8.36
8.36
8.35
8.35
7.32
7.31
7.26 CDCl3
7.25
7.23
7.04
7.04
7.02
7.02
6.95
6.93
1.64
TfO
OTf
O
O
O
tBuOOC
S5
1.00
1.09
1.94
1.08
2.11
2.06
9.16
f1 (ppm)

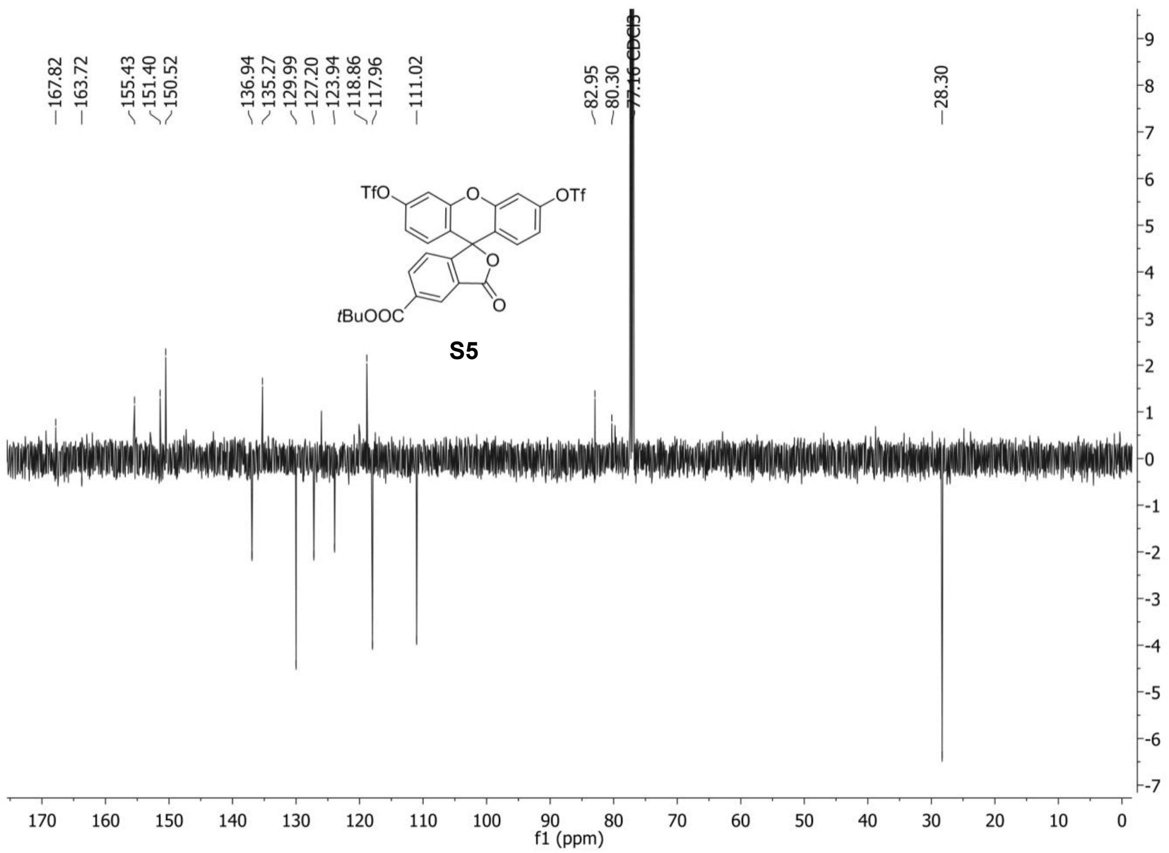
167.82
163.72
155.43
151.40
150.52
136.94
135.27
129.99
127.20
123.94
118.86
117.96
111.02
82.95
80.30
77.16 CDCl3
28.30
TfO
OTf
O
O
O
tBuOOC
S5
f1 (ppm)

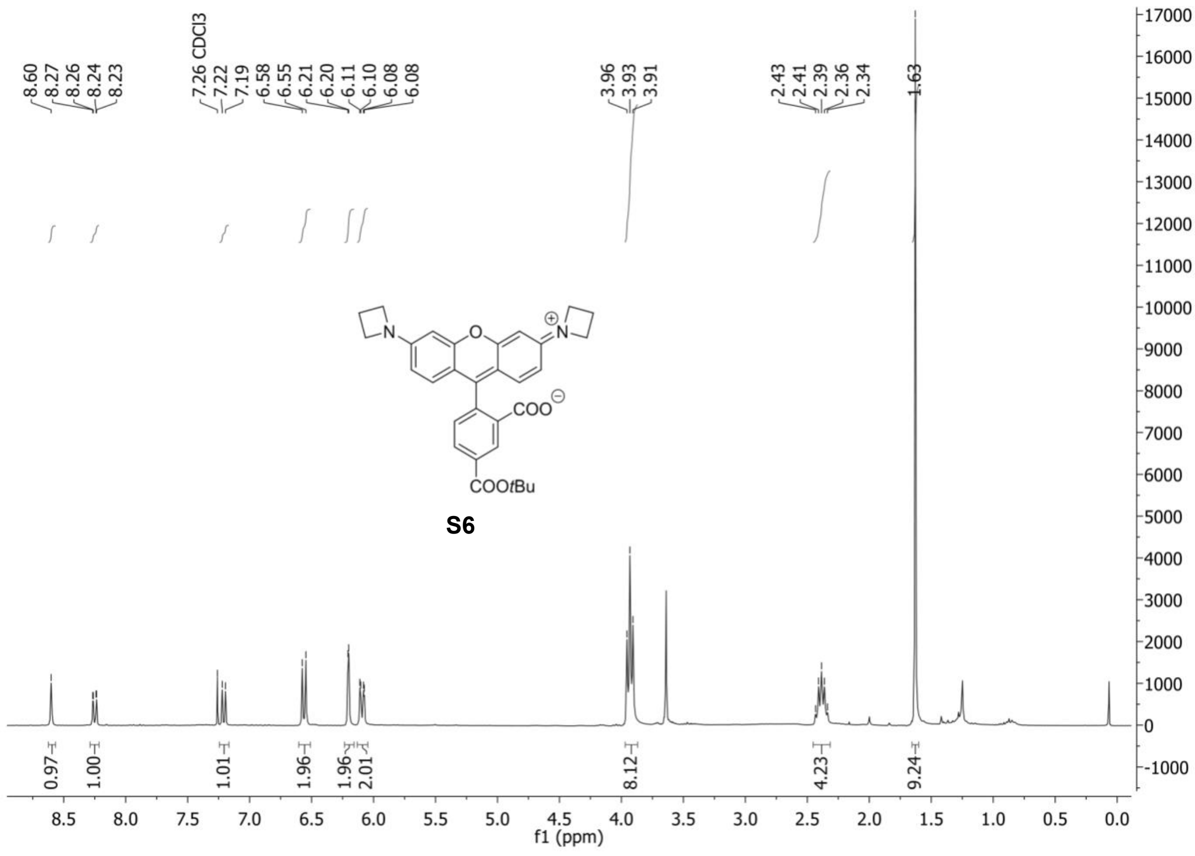
8.60
8.27
8.26
8.24
8.23
7.26 CDCl3
7.22
7.19
6.58
6.55
6.21
6.20
6.11
6.10
6.08
6.08
3.96
3.93
3.91
2.43
2.41
2.39
2.36
2.34
1.63
COO
COOtBu
S6
0.97
1.00
1.01
1.96
1.96
2.01
8.12
4.23
9.24
f1 (ppm)

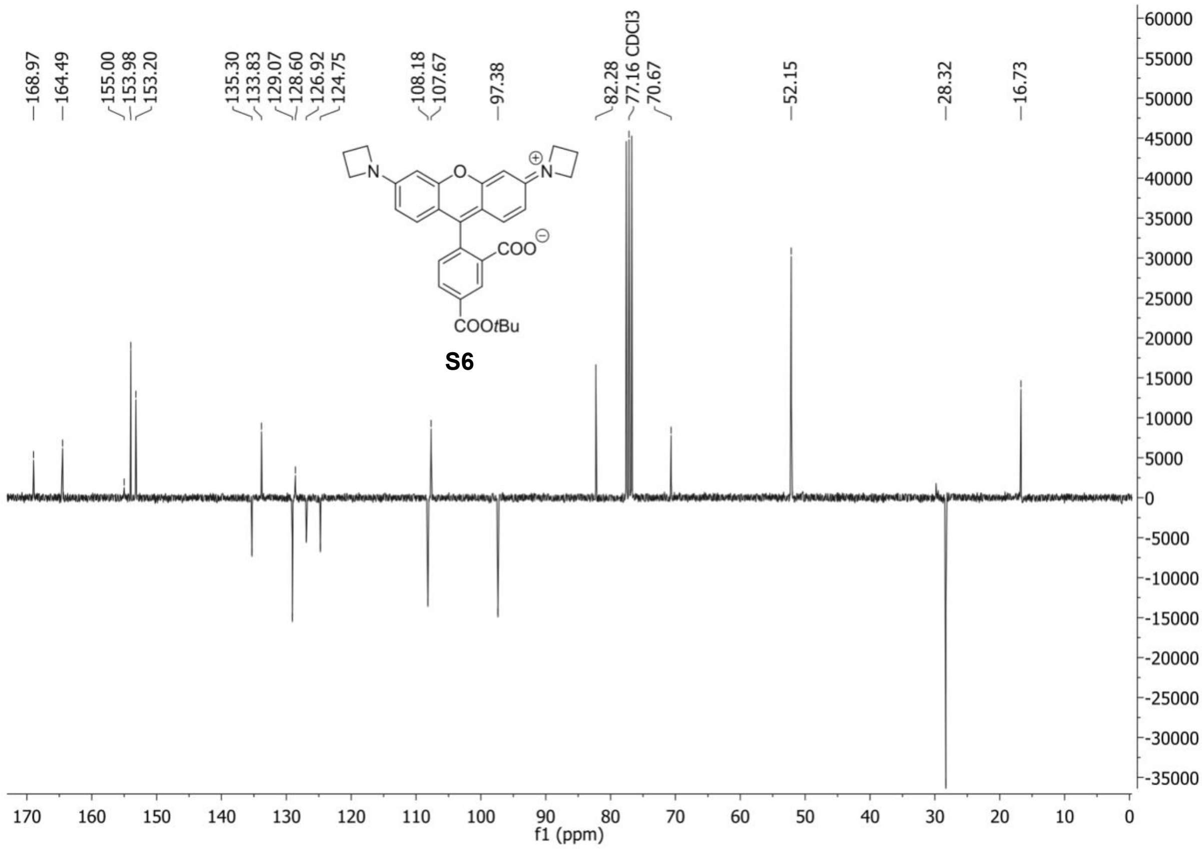
168.97
164.49
155.00
153.98
153.20
135.30
133.83
129.07
128.60
126.92
124.75
108.18
107.67
97.38
82.28
77.16 CDCl3
70.67
52.15
28.32
16.73
COO
COOtBu
S6
f1 (ppm)

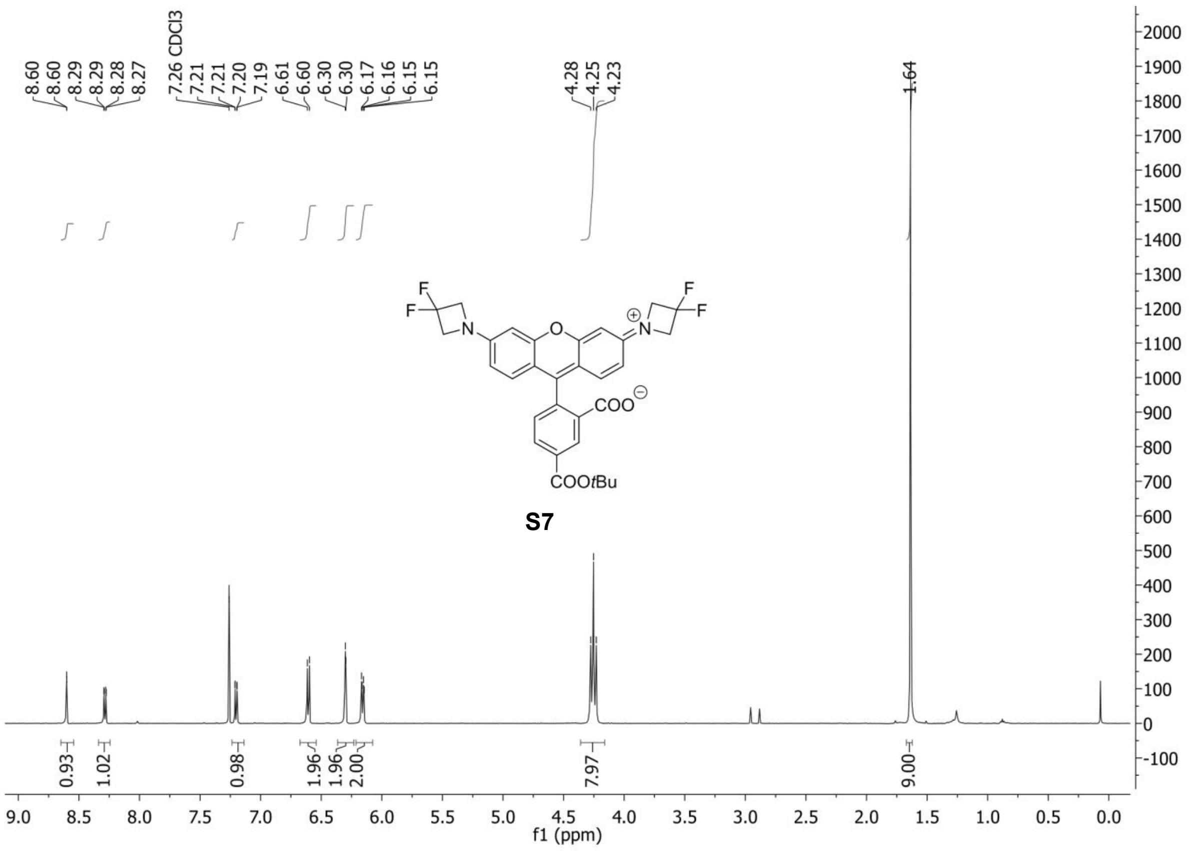
8.60
8.60
8.29
8.29
8.28
8.27
7.26 CDCl3
7.21
7.21
7.20
7.19
6.61
6.60
6.30
6.30
6.17
6.16
6.15
6.15
4.28
4.25
4.23
1.64
COO
COOtBu
S7
0.93
1.02
0.98
1.96
1.96
2.00
7.97
9.00
f1 (ppm)

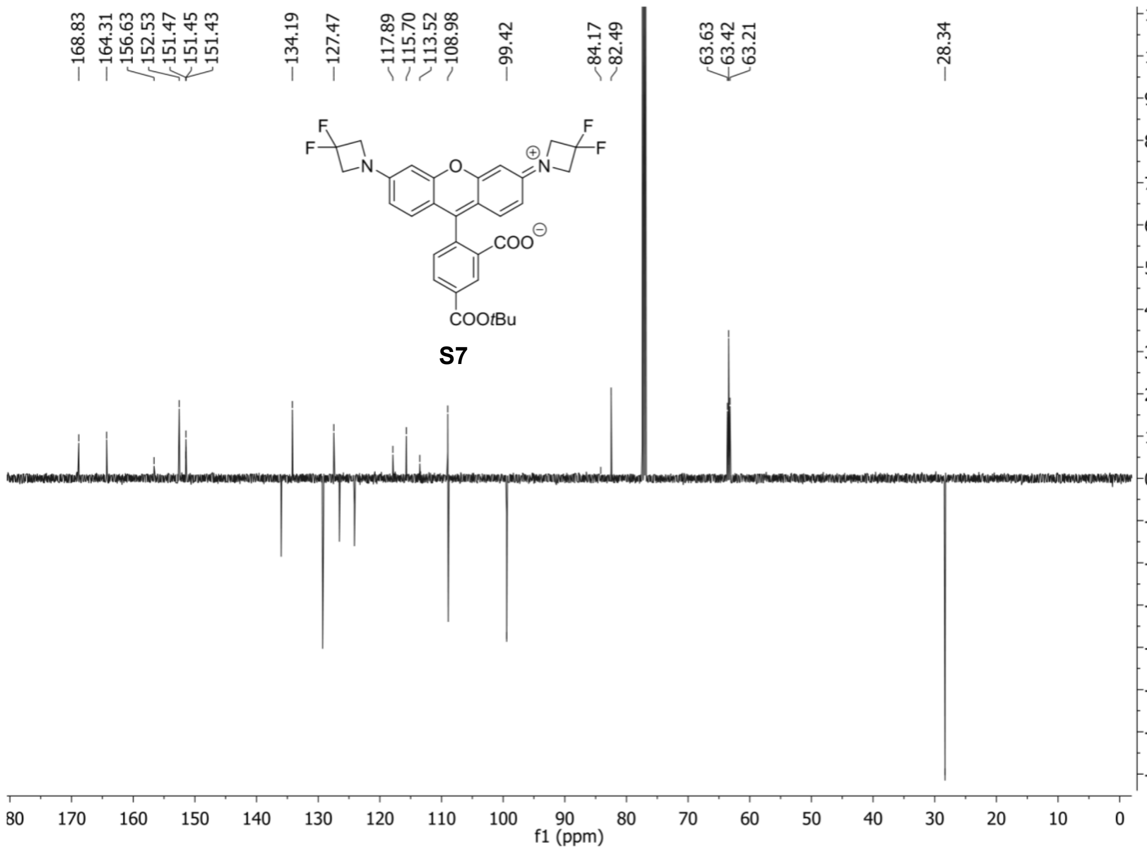
168.83
164.31
156.63
152.53
151.47
151.45
151.43
134.19
127.47
117.89
115.70
113.52
108.98
99.42
84.17
82.49
63.63
63.42
63.21
28.34
COO
COOtBu
S7
f1 (ppm)

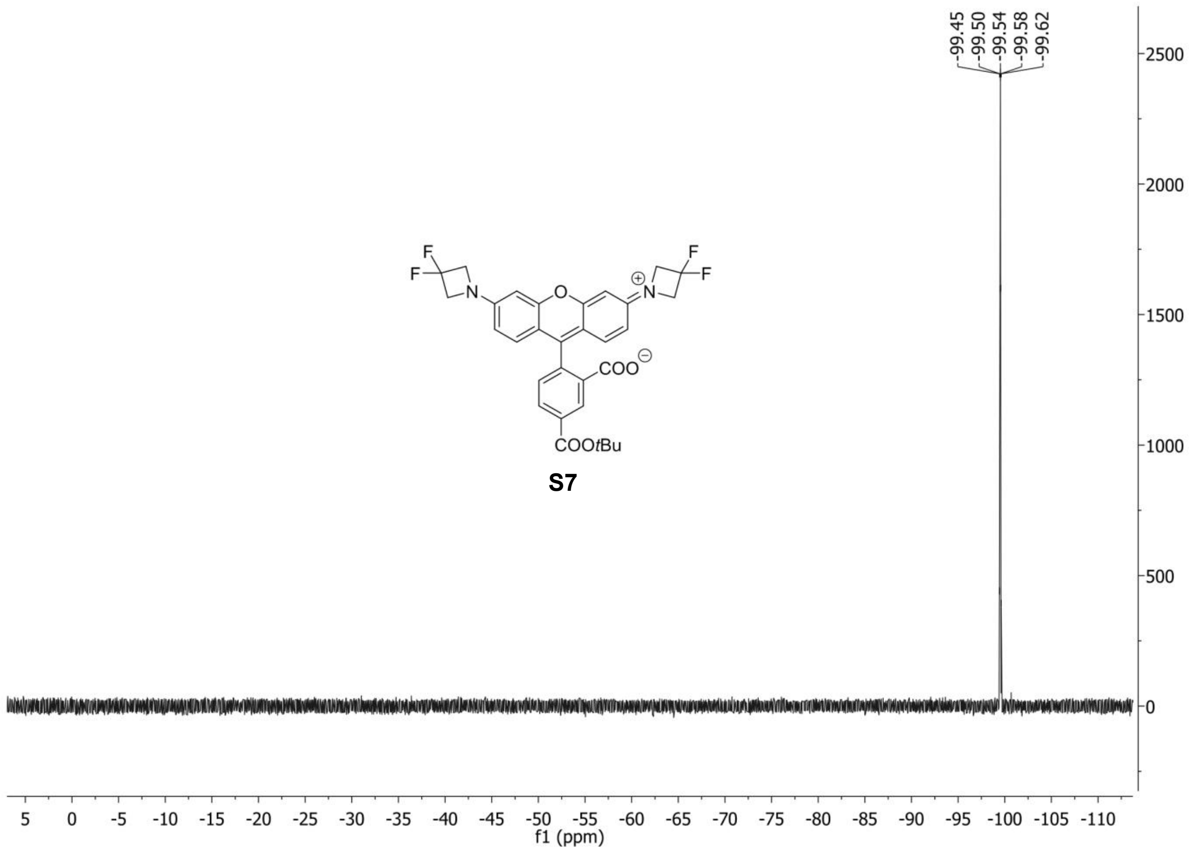
-99.45
-99.50
-99.54
-99.58
-99.62
F
F
N
O
N
F
F
COO
COOtBu
S7
2500
2000
1500
1000
500
0
5 0 -5 -10 -15 -20 -25 -30 -35 -40 -45 -50 -55 -60 -65 -70 -75 -80 -85 -90 -95 -100 -105 -110
f1 (ppm)

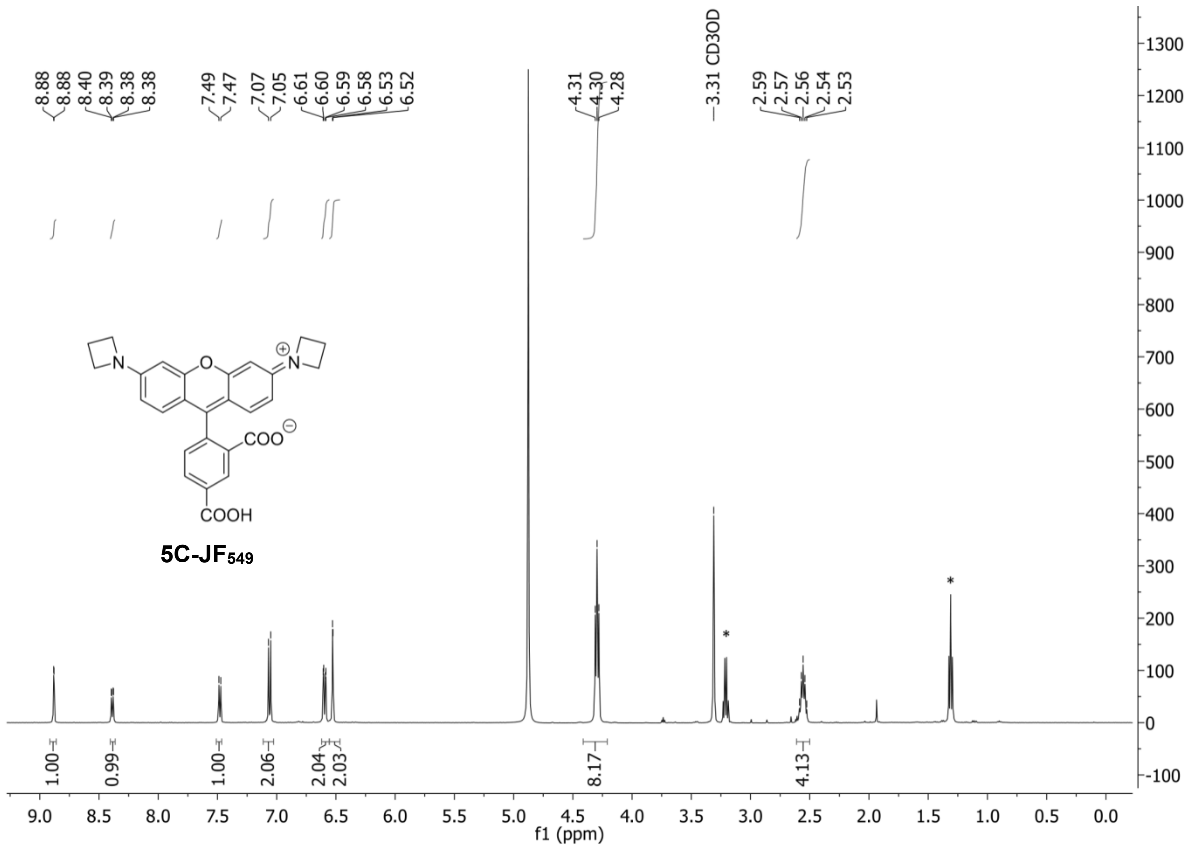

Signals indicated with * correspond to $Et_3N$.

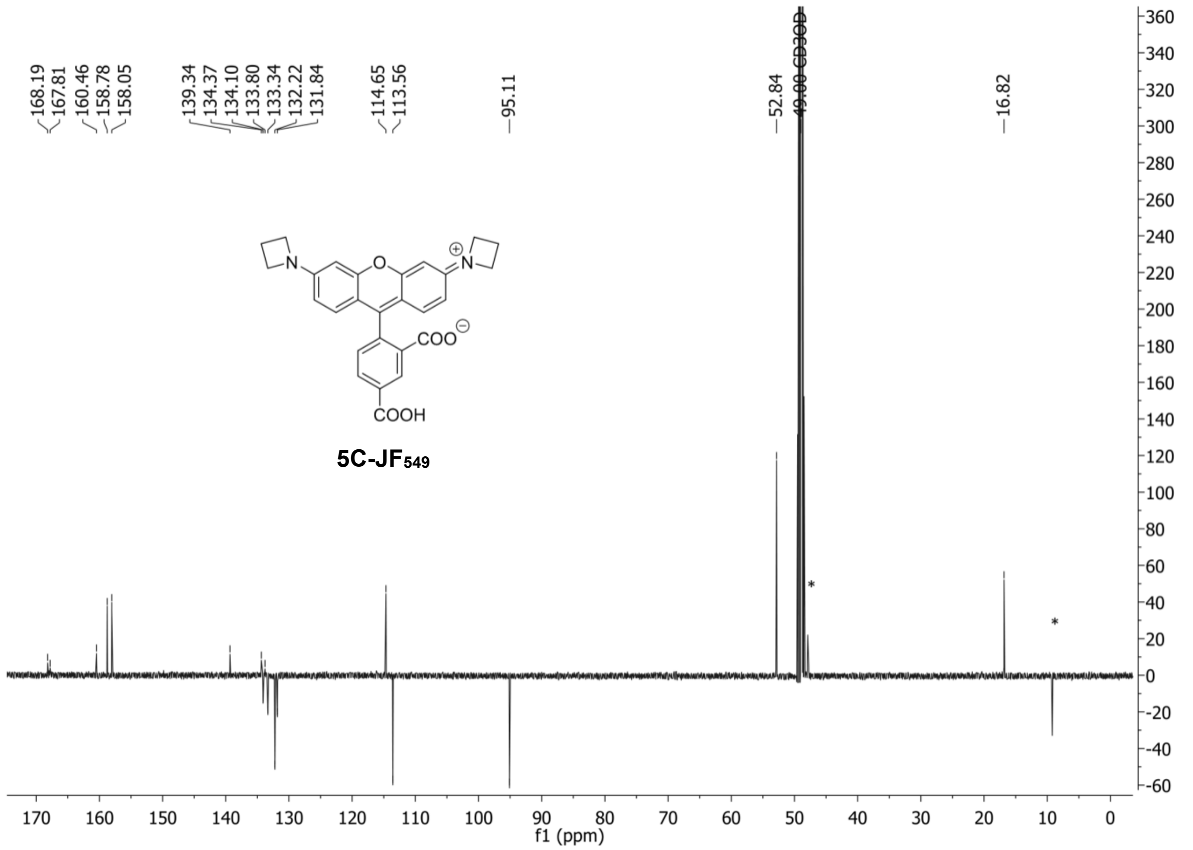

Signals indicated with * correspond to $Et_3N$.

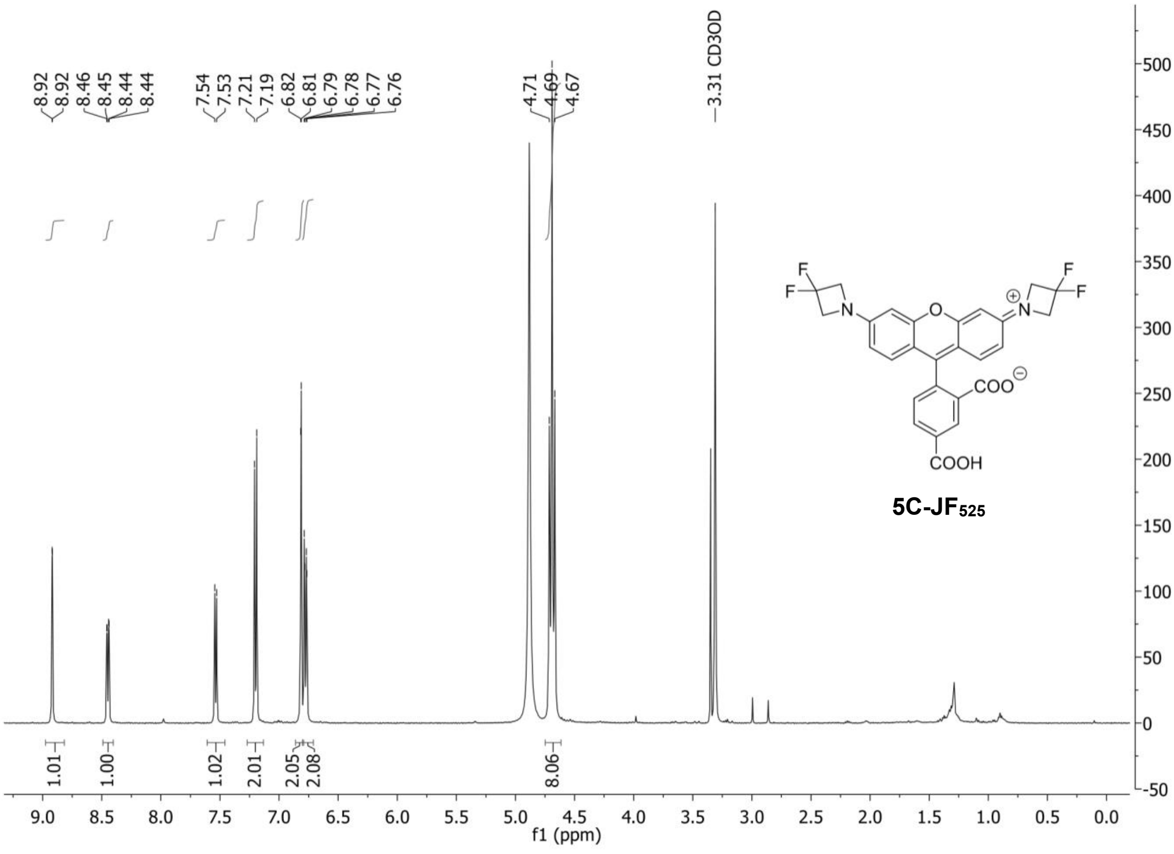
8.92
8.92
8.46
8.45
8.44
8.44
7.54
7.53
7.21
7.19
6.82
6.81
6.79
6.78
6.77
6.76
4.71
4.69
4.67
3.31 CD3OD
1.01
1.00
1.02
2.01
2.05
2.08
8.06
f1 (ppm)
5C-JF525

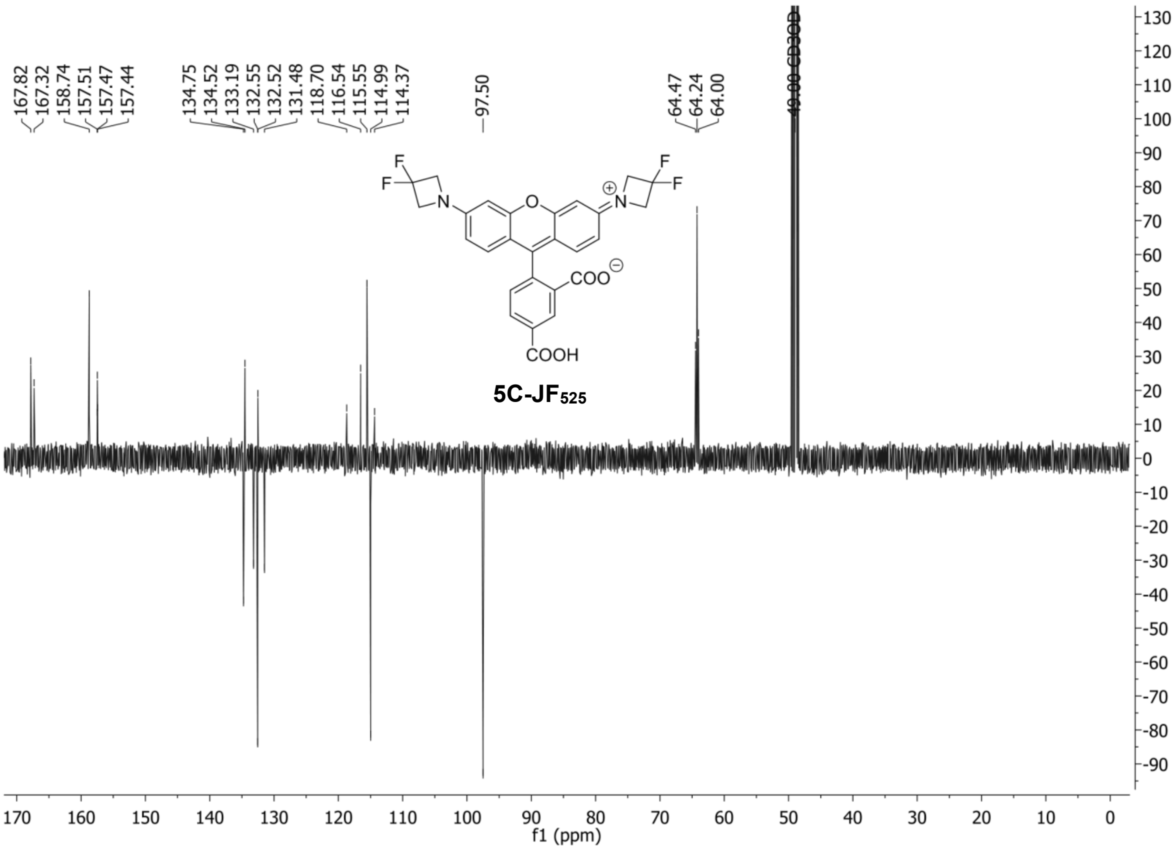
167.82
167.32
158.74
157.51
157.47
157.44
134.75
134.52
133.19
132.55
132.52
131.48
118.70
116.54
115.55
114.99
114.37
97.50
64.47
64.24
64.00
49.00 CD3OD
f1 (ppm)
5C-JF525

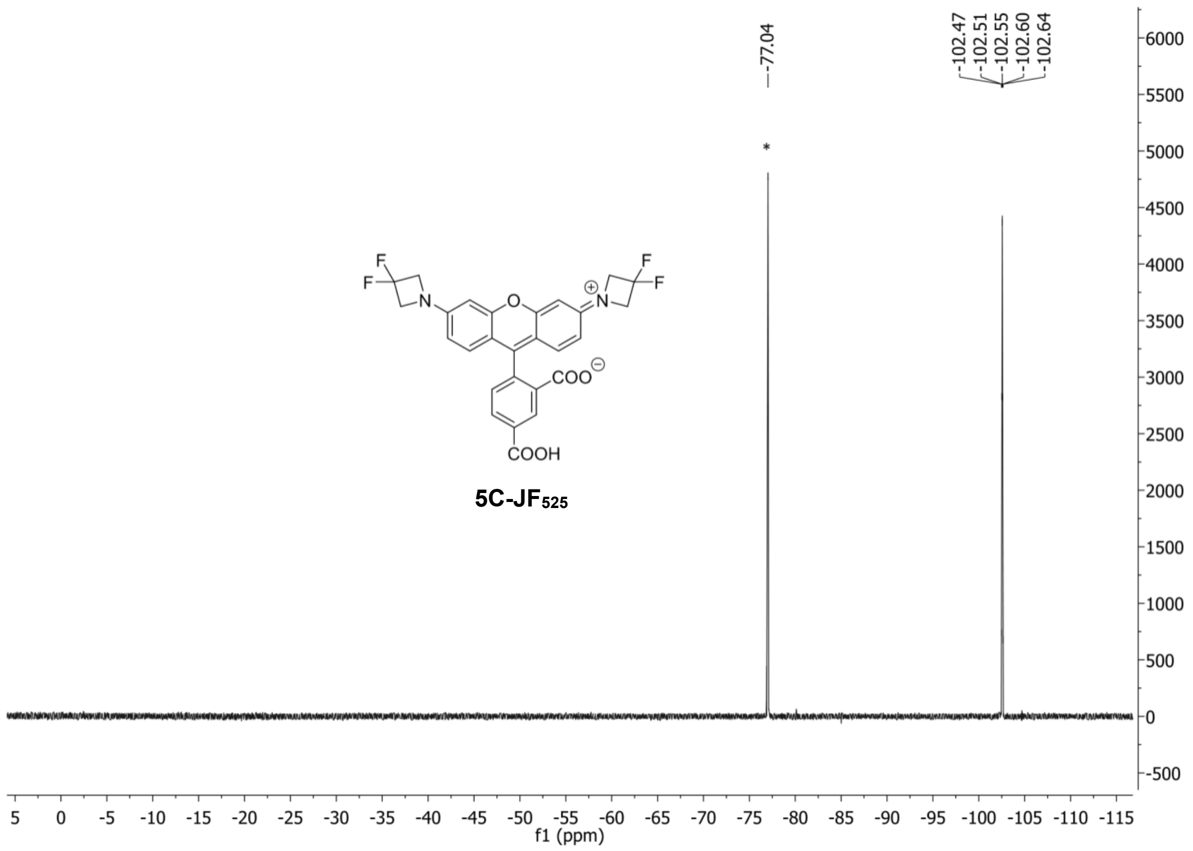


Signal indicated with * corresponds to TFA.

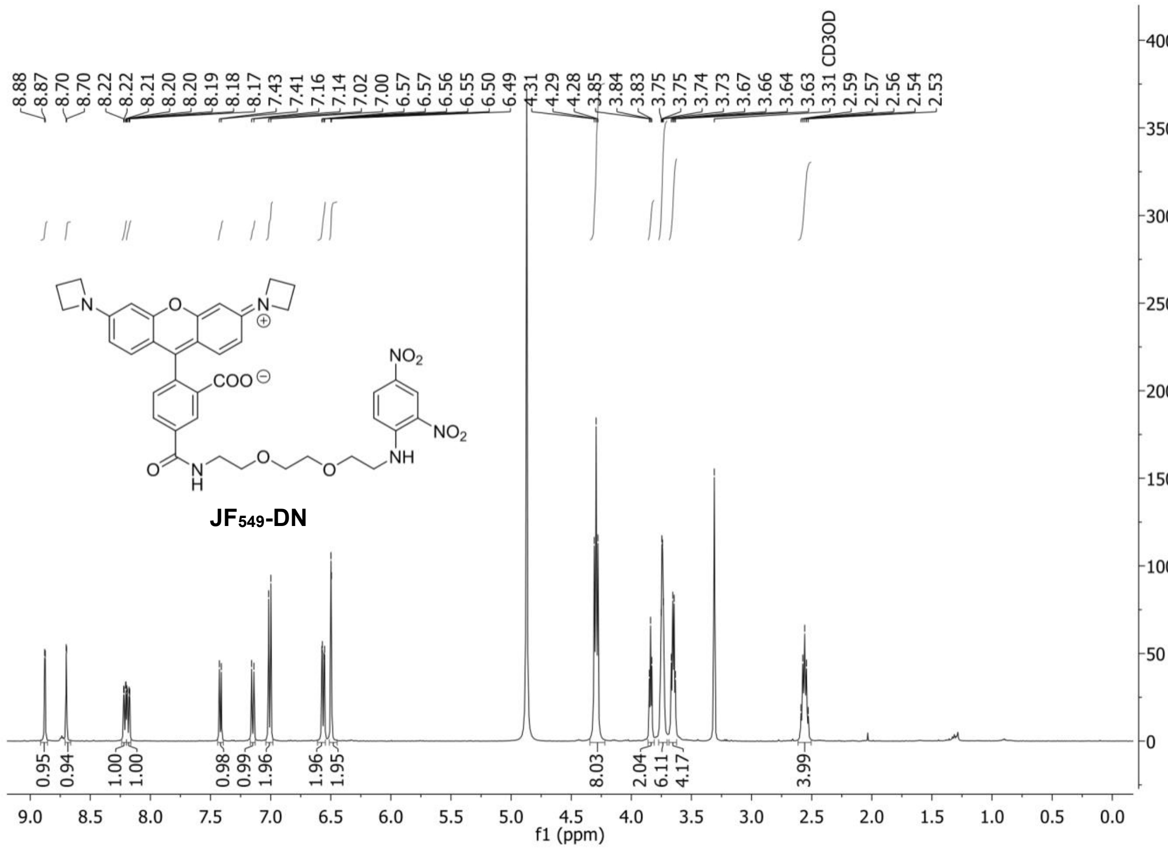
8.88
8.87
8.70
8.70
8.22
8.21
8.20
8.20
8.19
8.18
8.17
7.43
7.41
7.16
7.14
7.02
7.00
6.57
6.57
6.56
6.55
6.50
6.49
4.31
4.29
4.28
3.85
3.84
3.83
3.75
3.75
3.74
3.73
3.67
3.66
3.64
3.63
3.31 CD3OD
2.59
2.57
2.56
2.54
2.53
NO2
COO
NO2
NH
O
N
H
O
O
JF549-DN
0.95
0.94
1.00
1.00
0.98
0.99
1.96
1.96
1.95
8.03
2.04
6.11
4.17
3.99
f1 (ppm)

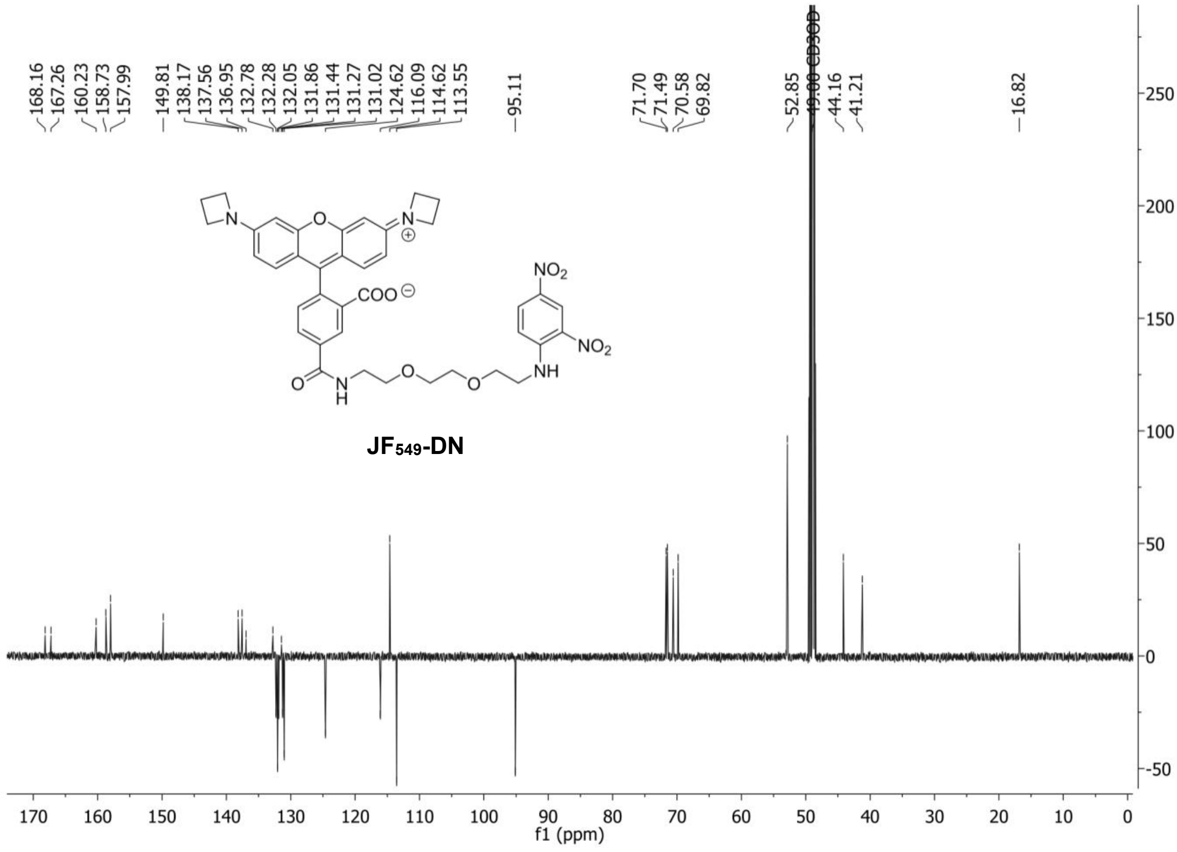
168.16
167.26
160.23
158.73
157.99
149.81
138.17
137.56
136.95
132.78
132.28
132.05
131.86
131.44
131.27
131.02
124.62
116.09
114.62
113.55
95.11
71.70
71.49
70.58
69.82
52.85
49.00 CD3OD
44.16
41.21
16.82
NO2
COO
NO2
NH
JF549-DN
f1 (ppm)

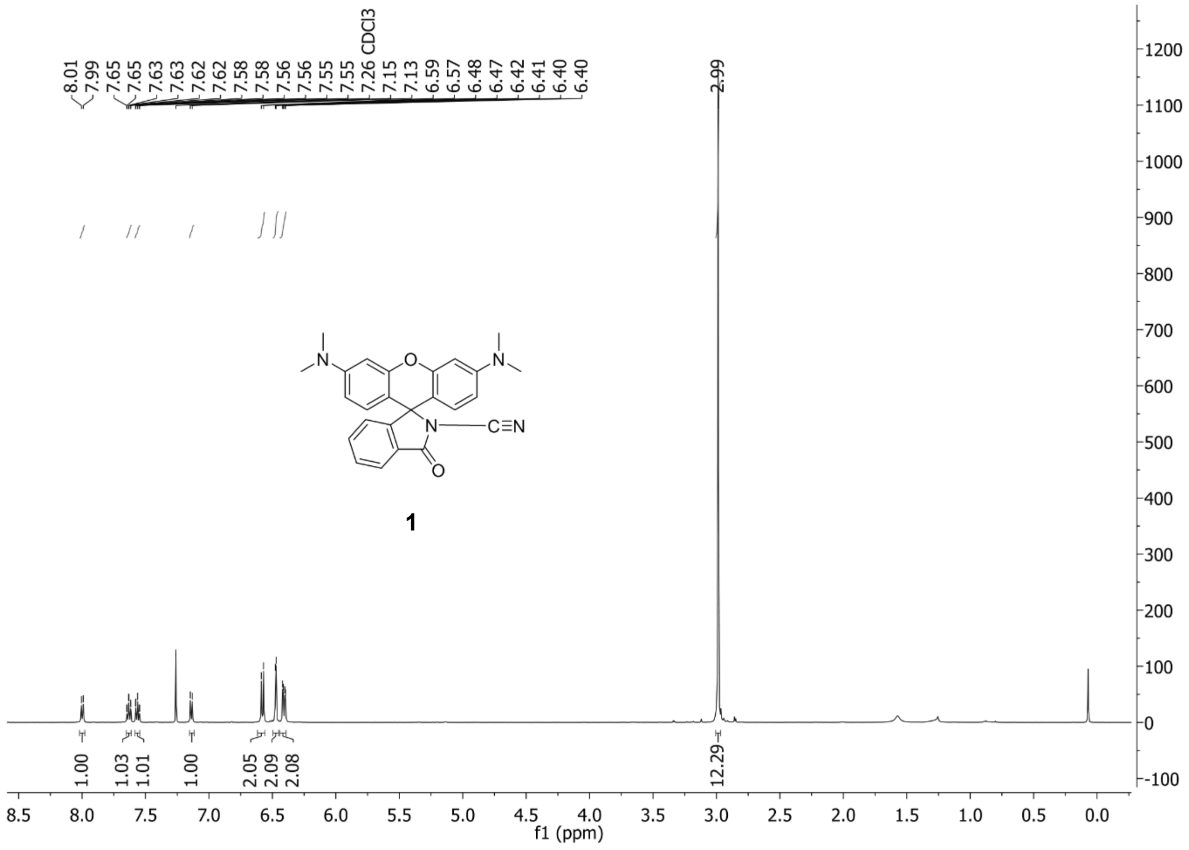
8.01
7.99
7.65
7.65
7.63
7.63
7.62
7.62
7.58
7.58
7.56
7.56
7.55
7.55
7.26 CDCl3
7.15
7.13
6.59
6.57
6.48
6.47
6.42
6.41
6.40
6.40
2.99
N
O
N
N
C≡N
O
1
1.00
1.03
1.01
1.00
2.05
2.09
2.08
12.29
8.5 8.0 7.5 7.0 6.5 6.0 5.5 5.0 4.5 4.0 3.5 3.0 2.5 2.0 1.5 1.0 0.5 0.0
f1 (ppm)
1200 1100 1000 900 800 700 600 500 400 300 200 100 0 -100

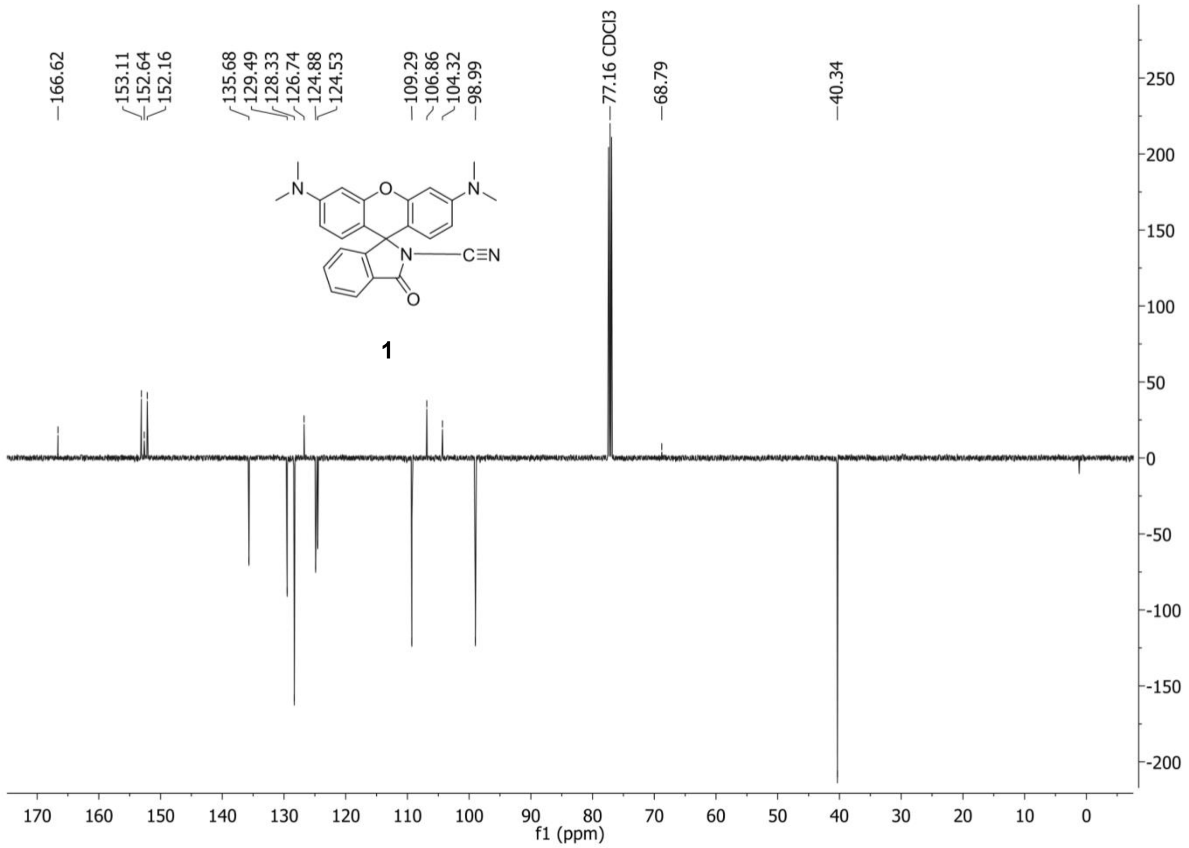
166.62
153.11
152.64
152.16
135.68
129.49
128.33
126.74
124.88
124.53
109.29
106.86
104.32
98.99
77.16 CDCl3
68.79
40.34
N
O
N
N
C≡N
O
1
170 160 150 140 130 120 110 100 90 80 70 60 50 40 30 20 10 0
f1 (ppm)
250 200 150 100 50 0 -50 -100 -150 -200

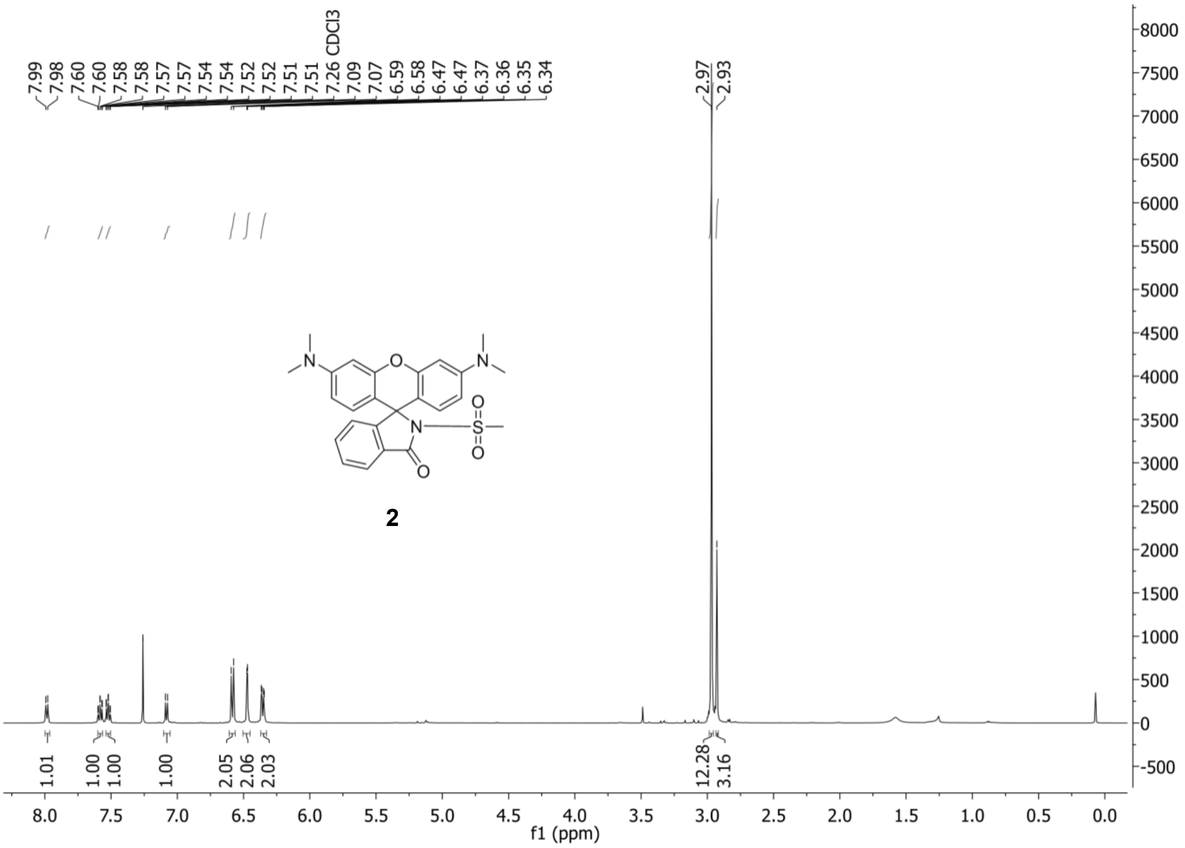
7.99
7.98
7.60
7.60
7.58
7.58
7.57
7.57
7.54
7.54
7.52
7.52
7.51
7.51
7.26 CDCl3
7.09
7.07
6.59
6.58
6.47
6.47
6.37
6.36
6.35
6.34
2.97
2.93
1.01
1.00
1.00
1.00
2.05
2.06
2.03
12.28
3.16
2
f1 (ppm)

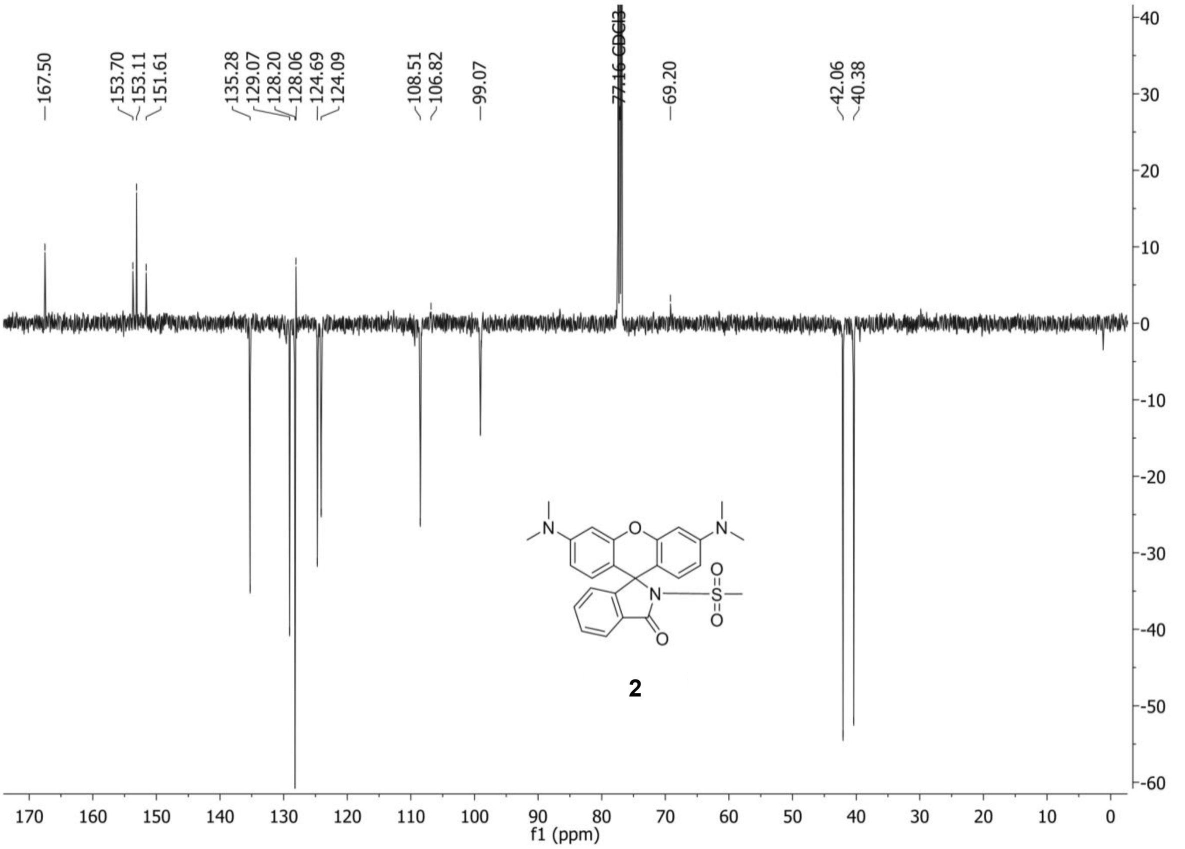
167.50
153.70
153.11
151.61
135.28
129.07
128.20
128.06
124.69
124.09
108.51
106.82
99.07
77.16 CDCl3
69.20
42.06
40.38
2
f1 (ppm)

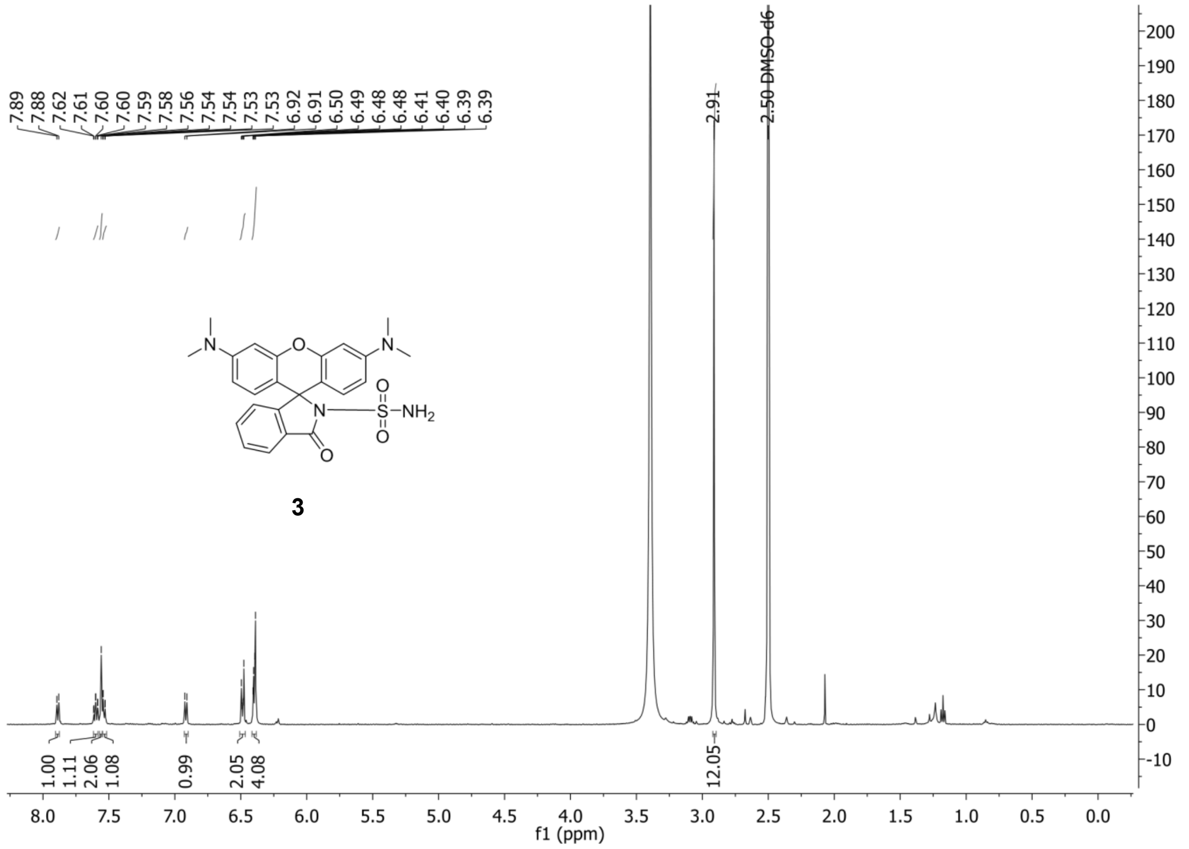
7.89
7.88
7.62
7.61
7.60
7.60
7.59
7.58
7.56
7.54
7.54
7.53
7.53
6.92
6.91
6.50
6.49
6.48
6.48
6.41
6.40
6.39
6.39
2.91
2.50 DMSO-d6
O
N
S
NH2
3
1.00
1.11
2.06
1.08
0.99
2.05
4.08
12.05
f1 (ppm)

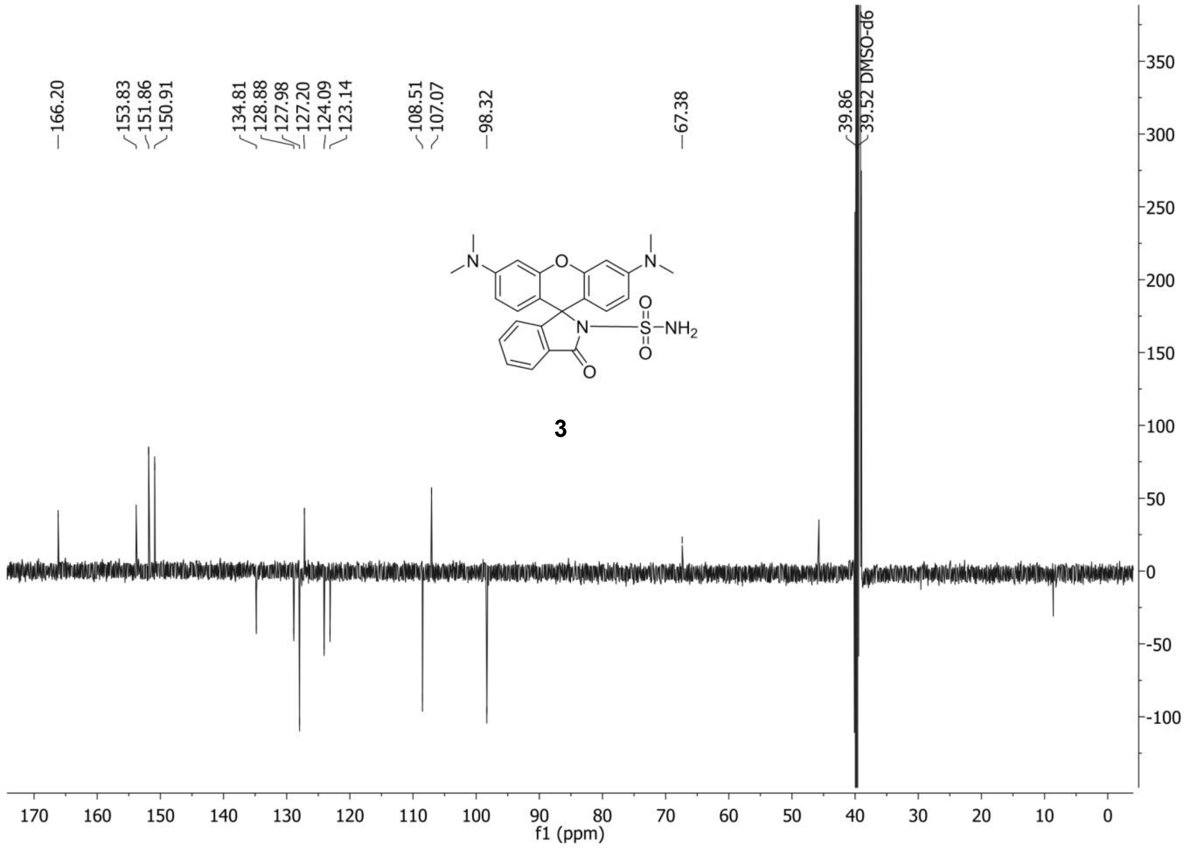
166.20
153.83
151.86
150.91
134.81
128.88
127.98
127.20
124.09
123.14
108.51
107.07
98.32
67.38
39.86
39.52 DMSO-d6
3
f1 (ppm)

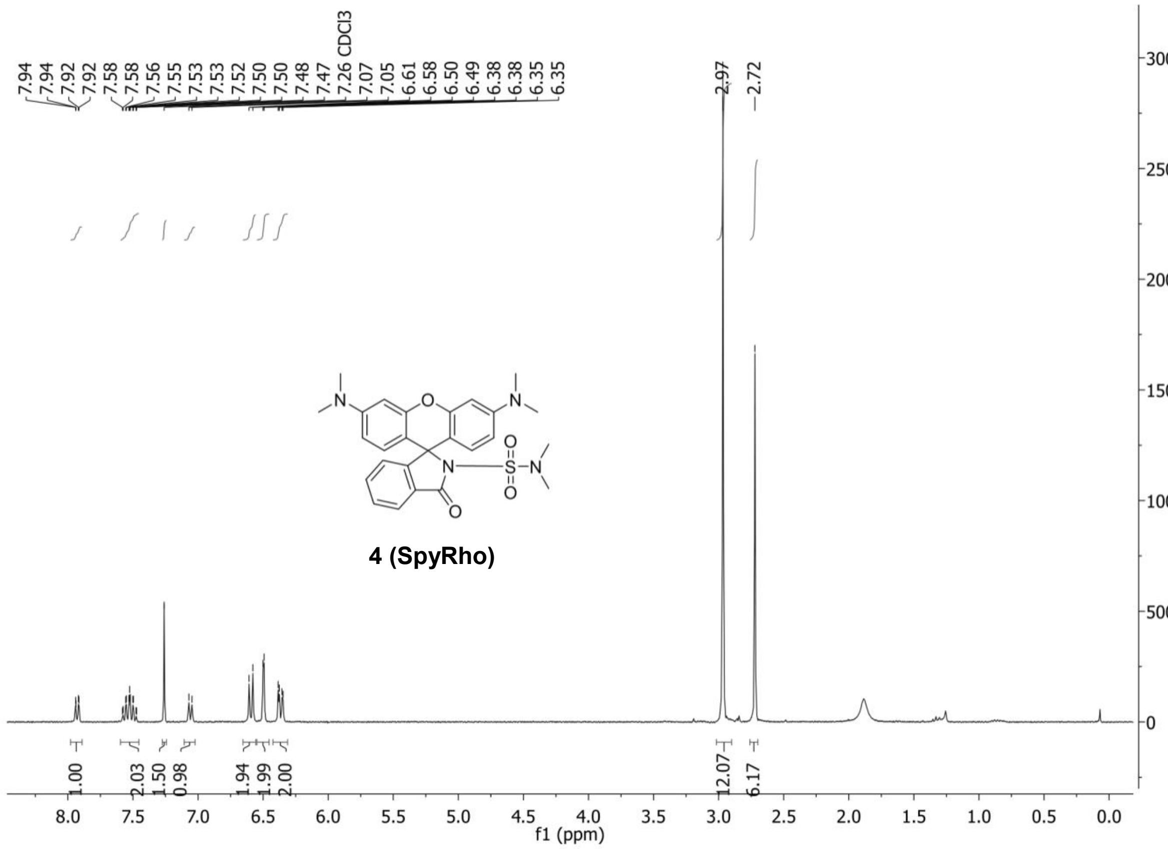
7.94
7.94
7.92
7.92
7.58
7.58
7.56
7.55
7.53
7.53
7.52
7.50
7.50
7.48
7.47
7.26 CDCl3
7.07
7.05
6.61
6.58
6.50
6.49
6.38
6.38
6.35
6.35
2.97
2.72
4 (SpyRho)
1.00
2.03
1.50
0.98
1.94
1.99
2.00
12.07
6.17
f1 (ppm)

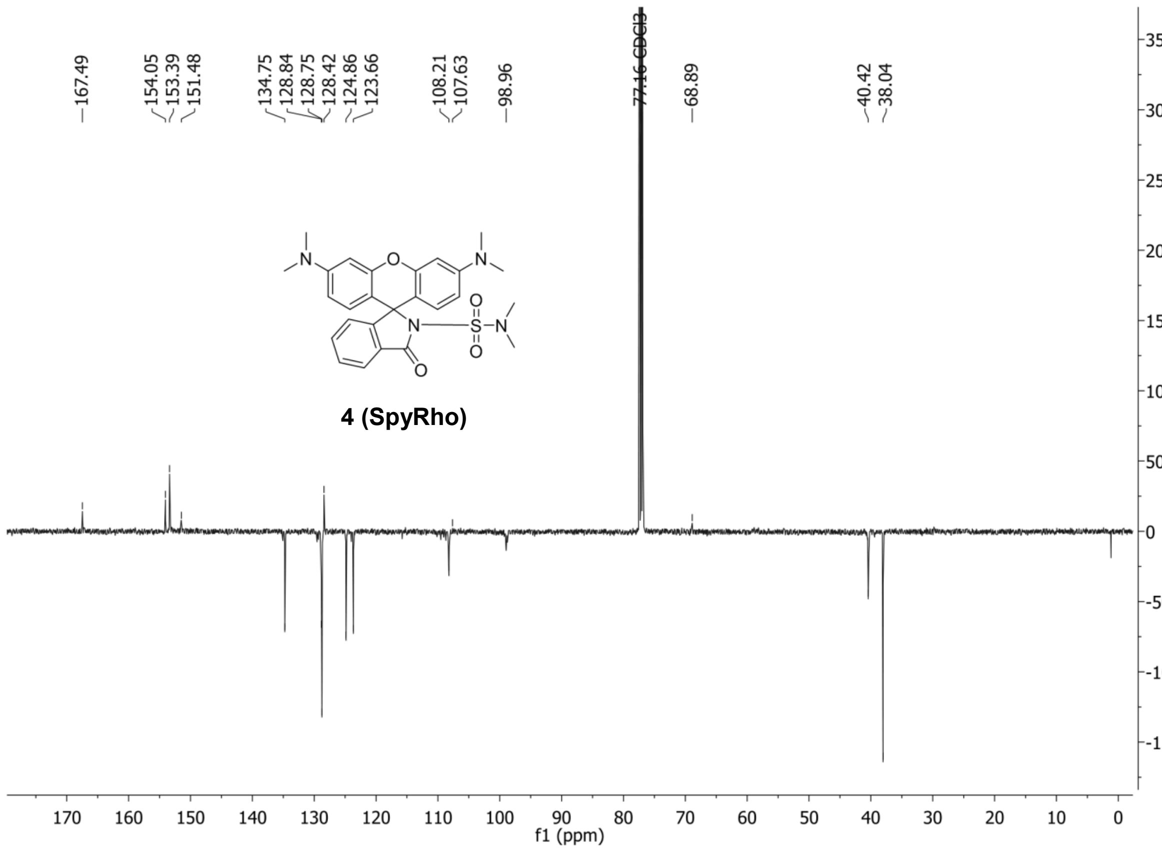
167.49
154.05
153.39
151.48
134.75
128.84
128.75
128.42
124.86
123.66
108.21
107.63
98.96
77.16 CDCl3
68.89
40.42
38.04
4 (SpyRho)
f1 (ppm)

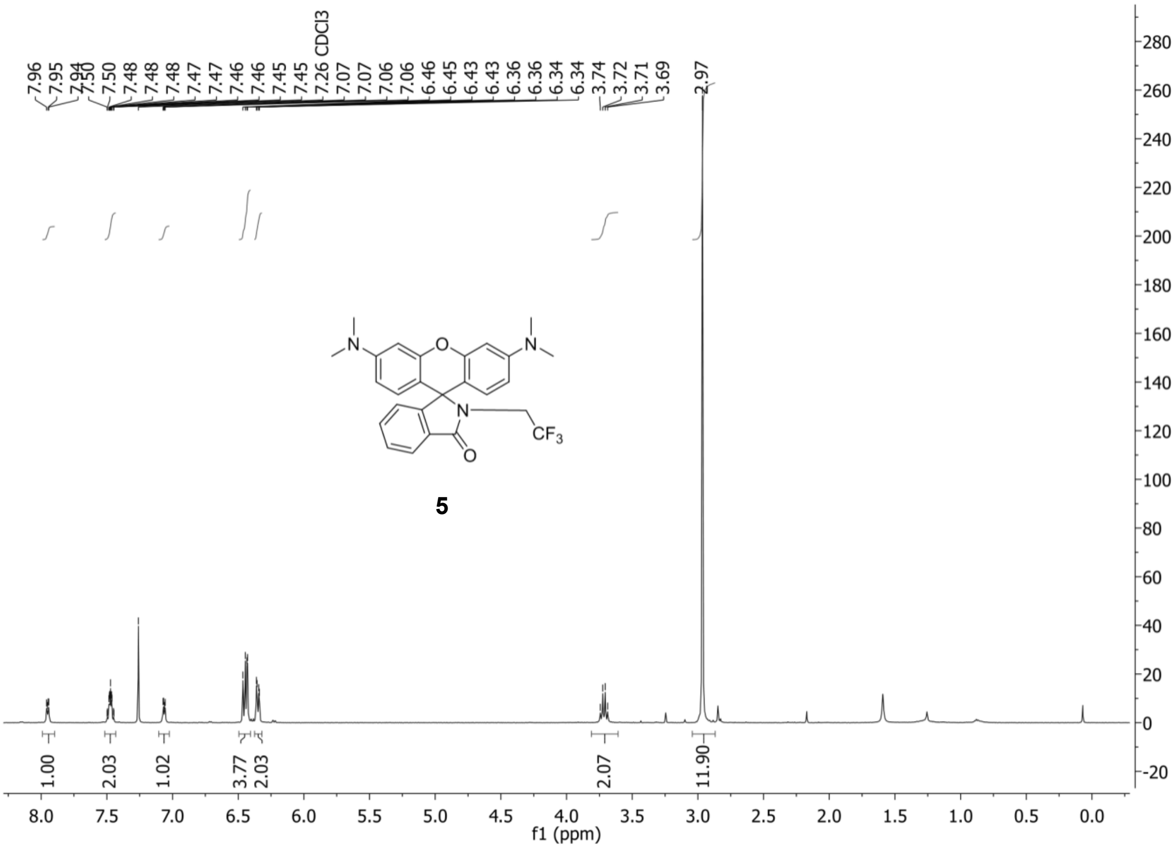
7.96
7.95
7.94
7.50
7.50
7.48
7.48
7.47
7.47
7.46
7.46
7.45
7.45
7.26 CDCl3
7.07
7.07
7.06
7.06
6.46
6.45
6.43
6.43
6.36
6.36
6.34
6.34
3.74
3.72
3.71
3.69
2.97
N
O
N
N
CF3
O
5
1.00
2.03
1.02
3.77
2.03
2.07
11.90
f1 (ppm)

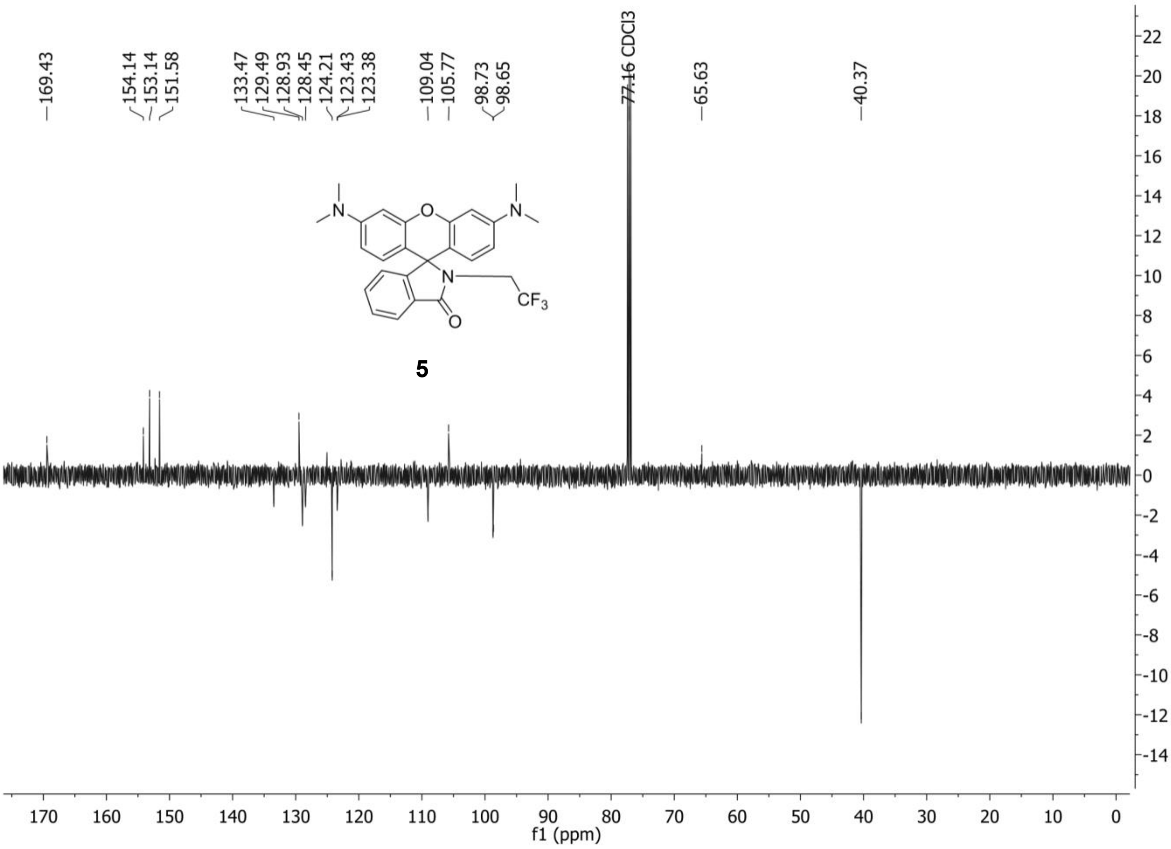
169.43
154.14
153.14
151.58
133.47
129.49
128.93
128.45
124.21
123.43
123.38
109.04
105.77
98.73
98.65
77.16 CDCl3
65.63
40.37
N
O
N
N
CF3
O
5
f1 (ppm)

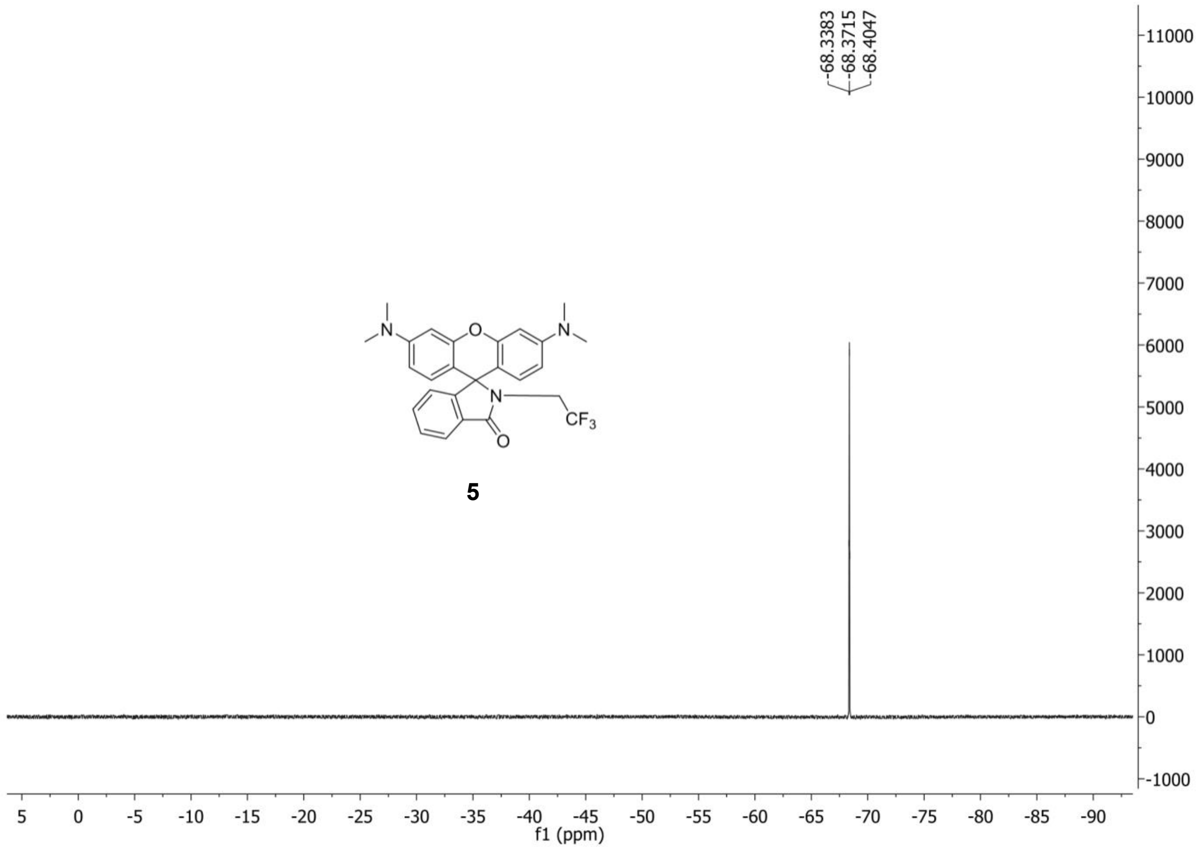
-68.3383
-68.3715
-68.4047
11000
10000
9000
8000
7000
6000
5000
4000
3000
2000
1000
0
-1000
5 0 -5 -10 -15 -20 -25 -30 -35 -40 -45 -50 -55 -60 -65 -70 -75 -80 -85 -90
f1 (ppm)
N
O
N
N
$CF_3$
O
5

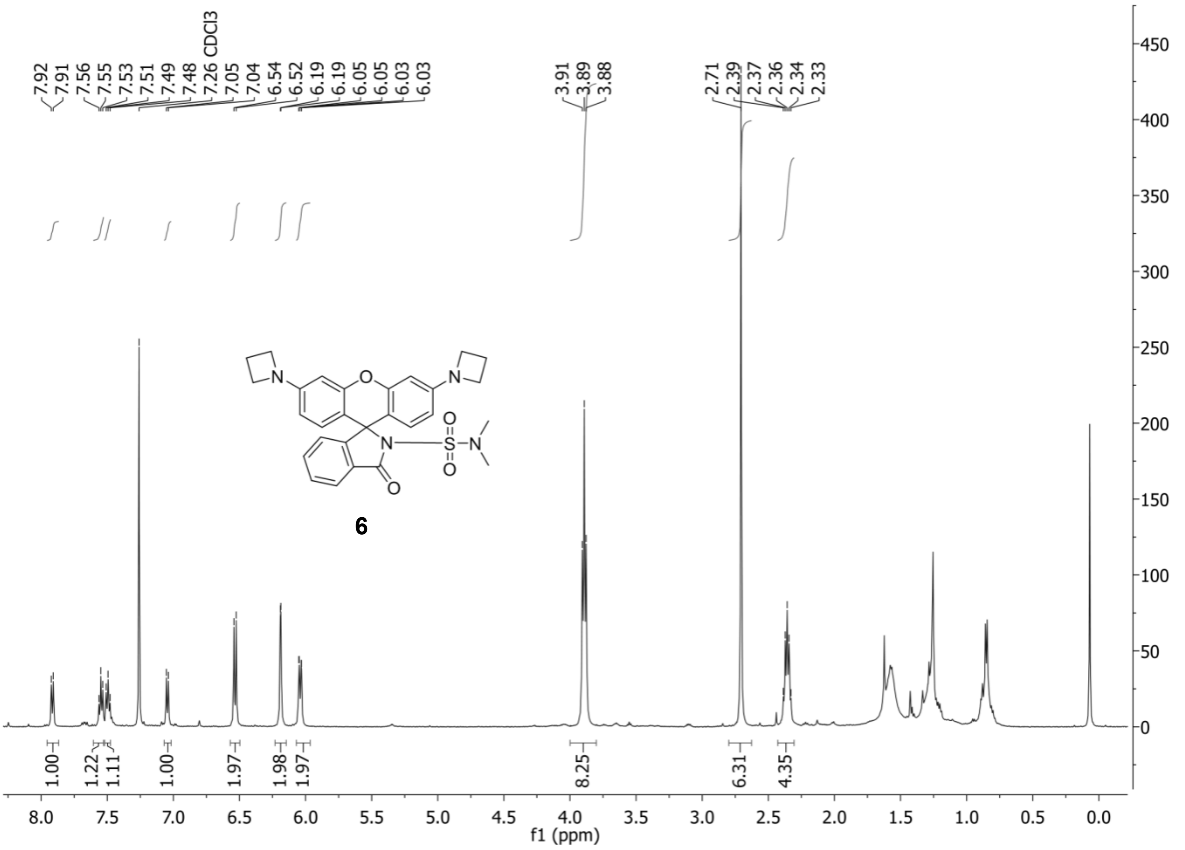
7.92
7.91
7.56
7.55
7.53
7.51
7.49
7.48
7.26 CDCl3
7.05
7.04
6.54
6.52
6.19
6.19
6.05
6.05
6.03
6.03
3.91
3.89
3.88
2.71
2.39
2.37
2.36
2.34
2.33
6
1.00
1.22
1.11
1.00
1.97
1.98
1.97
8.25
6.31
4.35
f1 (ppm)

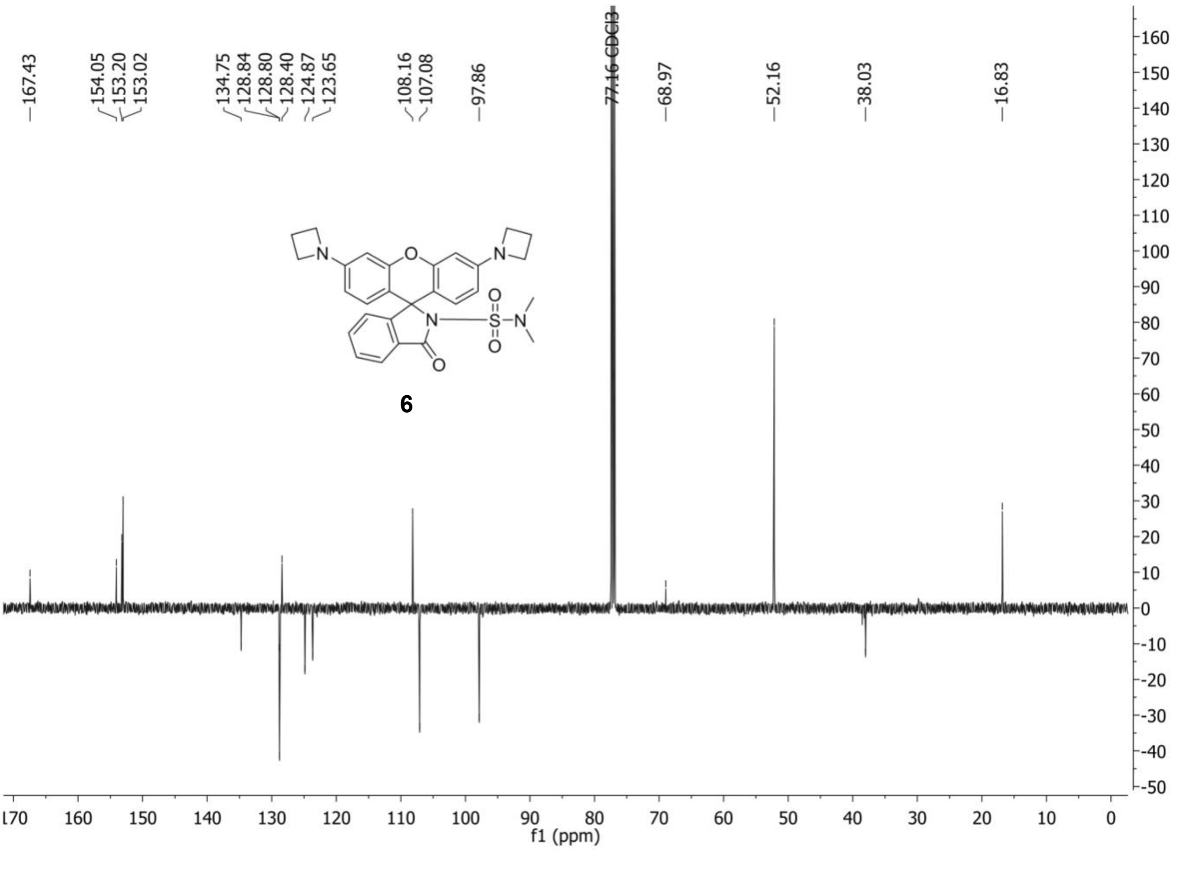
167.43
154.05
153.20
153.02
134.75
128.84
128.80
128.40
124.87
123.65
108.16
107.08
97.86
77.16 CDCl3
68.97
52.16
38.03
16.83
6
f1 (ppm)
